# Lateral inhibition in V1 controls neural & perceptual contrast sensitivity

**DOI:** 10.1101/2023.11.10.566605

**Authors:** Joseph Del Rosario, Stefano Coletta, Soon Ho Kim, Zach Mobille, Kayla Peelman, Brice Williams, Alan J Otsuki, Alejandra Del Castillo Valerio, Kendell Worden, Lou T. Blanpain, Lyndah Lovell, Hannah Choi, Bilal Haider

## Abstract

Lateral inhibition is a central principle for sensory system function. It is thought to operate by the activation of inhibitory neurons that restrict the spatial spread of sensory excitation. Much work on the role of inhibition in sensory systems has focused on visual cortex; however, the neurons, computations, and mechanisms underlying cortical lateral inhibition remain debated, and its importance for visual perception remains unknown. Here, we tested how lateral inhibition from PV or SST neurons in mouse primary visual cortex (V1) modulates neural and perceptual sensitivity to stimulus contrast. Lateral inhibition from PV neurons reduced neural and perceptual sensitivity to visual contrast in a uniform subtractive manner, whereas lateral inhibition from SST neurons more effectively changed the slope (or gain) of neural and perceptual contrast sensitivity. A neural circuit model identified spatially extensive lateral projections from SST neurons as the key factor, and we confirmed this with anatomy and direct subthreshold measurements of a larger spatial footprint for SST versus PV lateral inhibition. Together, these results define cell-type specific computational roles for lateral inhibition in V1, and establish their unique consequences on sensitivity to contrast, a fundamental aspect of the visual world.

## Introduction

Lateral inhibition is a core concept for sensory coding. It was discovered in the retina, where neural responses to a small spot of light became smaller when spatially adjacent regions were also co-illuminated^1^. Instead of increasing responses, the larger stimulus further restricted them in the effective region of visual space. Evidence for lateral inhibition shaping neural responses has since been found across sensory systems^2-7^. Nevertheless, the importance of lateral inhibition for sensory perception remains questioned, particularly from studies of primary visual cortex (V1).

One question concerns the perceptual effects of lateral inhibition in V1. Much work in V1 shows that lateral inhibition is unnecessary to explain several aspects of neural selectivity for stimuli in the central portion of the receptive field (RF)^8^. However, just like retinal neurons, V1 neurons improve their spatial sensitivity during costimulation of the RF plus the surrounding regions of space^9-13^. This modulation (or surround suppression^14^) is due to inhibition suppressing excitation within V1^10,11,15,16^. Remarkably, it is not known how lateral inhibition from a distant site in V1 affects perceptual sensitivity for stimuli appearing only in the RF.

A second question concerns how perceptual effects of lateral inhibition depend on cell types. Surround suppression in mouse V1 relies on somatostatin (SST) positive inhibitory neurons^15,17,18^, but the spatial scale of SST inhibition driving these effects remains unresolved^19^. Further, SST neurons strongly inhibit both excitatory neurons and parvalbumin (PV) interneurons^20-22^, which could counteract SST mediated suppression. PV neurons themselves show larger spatial integration than excitatory neurons^10,15,23,24^, and could also be involved in lateral suppression of excitation. Remarkably, the relationship between cell-type specific lateral inhibition and perception is unknown. To clarify this, it would be beneficial to activate lateral inhibition from PV or SST neurons independent from stimulus drive, and then measure how these activations propagate through V1 circuitry and affect stimulus perception. One possibility is that distant PV or SST neurons equally suppress excitation in the RF and equally alter perceptual sensitivity; an alternative is that anatomical and circuit properties of PV versus SST neurons confer unique consequences for perceptual sensitivity.

A third question concerns the computational role of lateral inhibition in V1. Several studies have examined computational roles for local inhibition (i.e., inhibition activated near the excitatory neurons that are driven by the visual stimulus). Moderate, sustained activation of local PV neurons (but not SST neurons) causes a divisive scaling of excitatory neuron responses to stimuli of different orientations^25-28^. Divisive scaling of the response curve –termed gain modulation– adjusts neural sensitivity without diminishing selectivity, a key neural computation underlying a variety of contextual effects^29-31^. Crucially, it is not known if scaling of V1 neural responses leads to scaling of perceptual responses. Furthermore, these prior studies were restricted to interactions within the local network; no studies have probed the computational role of lateral inhibition on both neural and perceptual response curves. This is important to clarify because lateral interactions among cortically (and retinotopically) distant sites have long been conjectured as a mechanism for behavioral and contextual modulation of perceptual sensitivity across the visual field^32,33^.

Here we tested how lateral inhibition from distant PV or SST neurons in mouse V1 modulates neural and perceptual sensitivity to stimulus contrast. We optogenetically activated inhibitory neurons that were laterally displaced from the task-relevant excitatory neurons by nearly 1mm in V1, along the gradient of horizontal visual space. Driving PV neurons reduced perceptual sensitivity uniformly across all contrasts, consistent with a subtractive shift of the psychometric contrast response curve. On the other hand, driving SST neurons caused a stronger divisive change of the contrast response curve. These perceptual effects of PV and SST lateral inhibition were mirrored by changes in neural contrast sensitivity on the same trials. A neural circuit model identified more extensive lateral projections from distal SST neurons – which we confirmed anatomically – as a key factor driving divisive scaling of response curves. Patch-clamp recordings in awake V1 directly revealed a larger spatial footprint for SST versus PV lateral inhibition. Together, these results define cell-type specific computational roles for lateral inhibition in V1, and establish their unique consequences on perception.

## Results

### SST lateral inhibition controls the slope of psychometric contrast sensitivity

To probe the perceptual effects of PV versus SST lateral inhibition, we trained water-restricted, head-fixed stationary mice to perform a visual detection task^34,35^. We expressed channelrhodopsin (ChR2) in PV or SST neurons by crossing Ai32 mice with PV- or SST-cre mice (abbreviated as PV-ChR2 or SST-ChR2 mice). Mice learned to report detection of small Gabor gratings (σ = 10°) appearing in discrete locations in the visual field, as in our prior studies^34,35^. We focused on blocks of trials where the stimulus appeared at the vertical meridian (defined as 0° azimuth) in the binocular region of greatest visual sensitivity^34,36^. Visual stimuli appeared at multiple contrasts (0-33%) so that we could probe how lateral inhibition impacted psychometric contrast sensitivity (Fig. 1A). V1 sites for silicon probe neural recordings and laser stimulation were precisely targeted in every experiment using hemodynamic imaging, per our prior studies^37^. Recording sites in V1 targeted the retinotopic location of the detected gratings (Fig. 1B, C; Fig. S1) while the laser spot (∼0.3mm in width; Fig. S1) was positioned far away from the recording site (∼0.8mm) and stimulated neurons with spatial receptive fields 70° away from the detected gratings (Fig. S1), a distance much larger than excitatory neuron RF size^23^. On a randomized subset of trials (25-33%), we activated PV or SST neurons at this distal location using moderate laser power (1.7mW) that modulated behavioral performance without abolishing it. Moderate and sustained laser activation lasted for the duration of the visual stimulus, as in prior studies of local inhibition^28,38^. This experimental design allowed us to isolate the effects of lateral inhibitory drive from stimulus drive, and then measure how lateral inhibition from distinct inhibitory neurons transforms neural and perceptual sensitivity to contrast.

**Figure 1.**
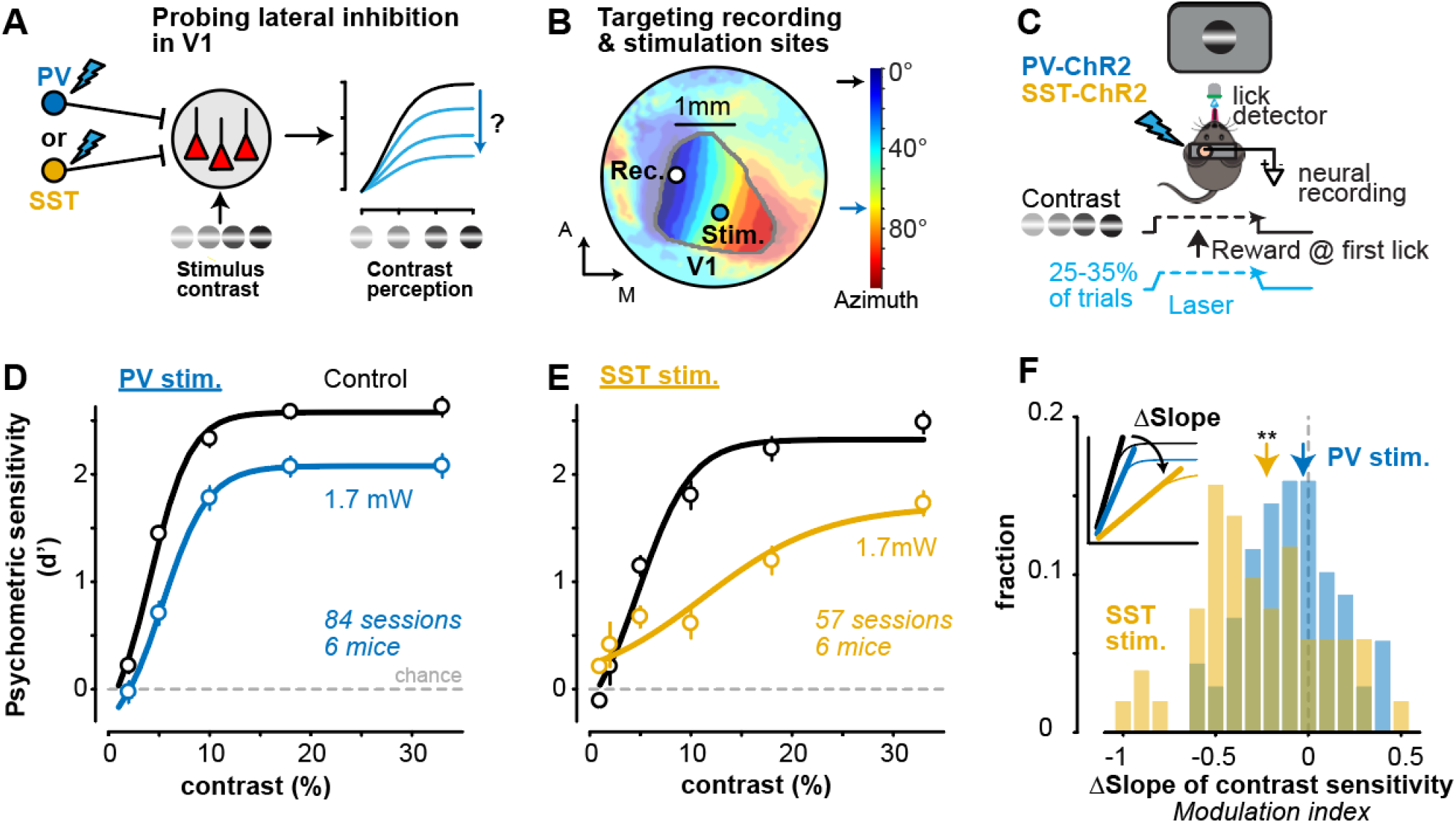
SST lateral inhibition controls the slope of psychometric contrast sensitivity. **A**. Experimental design to probe how lateral inhibition from PV (blue) or SST (gold) neurons in primary visual cortex (V1) affects neural and perceptual contrast sensitivity. **B**. Example hemodynamic response map of horizontal visual space (azimuth, colorbar). Visual field sign (Methods) used to delimit V1 (contour) and target recording sites (0°) and laser stimulation (70°, ∼0.8 mm away from recording site) in every experiment. See Fig. S1 for laser spot size (∼0.3mm) and recording locations across experiments. **C**. SST- or PV-cre x Ai32 channelrhodopsin (ChR2) mice were trained to detect static Gabor gratings (location = 0°; σ = 10°) at multiple contrasts (0-33%). PV or SST neurons were activated on 25% or 33% of trials with a laser ramp (1.7mW at peak) during grating appearance. Neural activity was recorded simultaneously from the V1 retinotopic site corresponding to grating location (as in B). **D**. Psychometric contrast sensitivity (d’) in PV-ChR2 mice (6 mice, 84 sessions) for interleaved control (black) and PV inhibition trials (blue). Mean ± SEM, fit with a sigmoidal equation (Methods). **E**. Same as D for SST-ChR2 mice (6 mice, 57 sessions). **F**. SST stimulation significantly decreased the slope of the contrast response curve (−0.23 ± 0.26, median ± MAD; *p* < 1e-3, Wilcoxon signed-rank test), but PV stimulation did not (−0.03 ± 0.20; *p* = 0.22). Change in slope quantified as modulation index (MI), defined as difference between control and laser conditions (inset) divided by sum (see Methods). Histogram shows distributions of MI calculated per session (same sessions in D, E). Changes not explained by differences in false alarms, pupil dynamics, or other behavioral factors in PV vs SST mice (Fig. S2). Affine model shows greater divisive gain change with SST stimulation, and greater subtractive offset change with PV stimulation (Fig. S4).

PV stimulation diminished detectability of all contrasts in a uniform subtractive manner; however, SST stimulation resulted in a divisive scaling of the slope of the contrast sensitivity function. Both forms of inhibition caused significant decreases in overall performance, and an increase in the contrast threshold (C_50_; Fig. S2), but the clearest effect was a difference in scaling of the entire contrast response function (dynamic range). We quantified these effects in individual sessions by calculating a psychometric slope modulation index (MI), defined as the difference in contrast response function slopes for control and laser trials, divided by their sum (Fig. 1F). SST stimulation significantly decreased the slope of psychometric sensitivity (Fig. 1F; *p <* 1e-3), but PV stimulation did not (*p* = 0.22). These different forms of SST vs PV modulation were not explained simply by overall differences in behavioral performance or by differences in pupil area (linked to arousal^39^) or pupil position across groups (Fig. S3). Finally, fitting an affine model to the control psychometric functions — then transforming them with a gain and offset parameter to explain photostimulation trials — revealed that SST stimulation caused a significantly greater change in the gain factor (division), while PV stimulation caused a significantly greater change in offset (subtraction; Fig. S4). Thus, lateral inhibition from SST neurons was more effective in driving divisive gain modulation of the contrast sensitivity function.

### Cell-type specific lateral inhibition adjusts neural contrast sensitivity during perception

Are these perceptual changes in contrast sensitivity mirrored by changes in V1 neural activity? We performed extracellular silicon probe recordings in V1 during the task, and found that lateral inhibition controlled the contrast sensitivity of putative excitatory neurons. We optically targeted recordings to the retinotopic site of the visual stimulus, and stimulated PV or SST neurons far away (0.8mm) from the retinotopic representation of the visual stimulus (Fig. 1B). We classified neurons with broad waveforms as regular spiking (RS) putative excitatory neurons (Methods; Fig. S5C), and focused on those with contrast dependent responses measured in control conditions (Fig. 2; r^2^ > 0.25 with Naka-Rushton fit; 312/657 or 47% of neurons in both PV and SST mice). Distal activation of PV (Fig. 2A-B) or SST neurons (Fig. 2D-E) reduced the visually-evoked transient response of RS neurons, but in different ways: PV stimulation reduced spiking by a similar factor across contrasts (Fig. 2C), while SST activation more effectively reduced low and medium contrast responses to near baseline levels (Fig. 2F), more consistent with a change in slope. Both conditions caused overall reductions in firing rates (R_max_), and increased neural contrast thresholds (C_50_; Fig. S6). We further dissected these average effects session-by-session to determine the strength and correlation of neural and behavioral contrast sensitivity.

**Figure 2.**
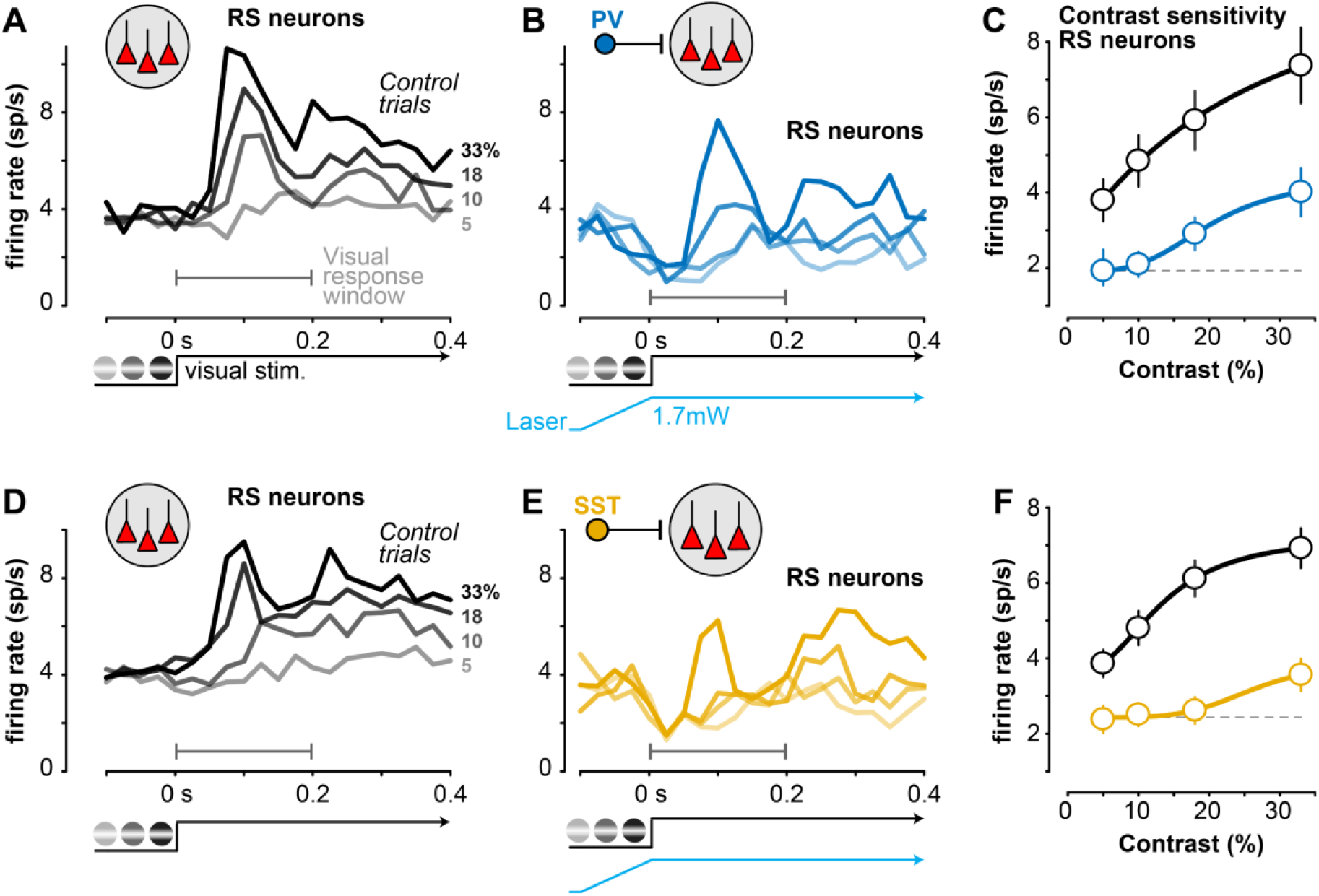
Cell-type specific lateral inhibition adjusts neural contrast sensitivity during perception. **A**. Average responses of V1 putative excitatory regular spiking (RS) neurons (n=166) during contrast detection (sorted by contrast, right) in PV-ChR2 mice (4 mice, 26 sessions). Recordings at V1 retinotopic location of stimulus (Fig. 1B). **B**. Same sessions and neurons as A during interleaved trials of distal PV stimulation, sorted by contrast. **C**. Contrast response curve (mean ± SEM) during control (black) and PV stimulation (blue, gray-dashed line shows firing rate at lowest contrast). Response calculated during visual response window (first 0.2s after stimulus in A and B). **D-F**. Same as A-C, for SST-ChR2 mice (n = 146 RS neurons; 4 mice, 18 sessions).

### SST inhibition drives correlated reduction of neural and perceptual contrast sensitivity

For all contrast tuned RS neurons recorded in the task, we first measured how contrast response functions changed with PV or SST distal inhibition, then compared this to the perceptual effects on the same trials. We again defined a modulation index (MI) to quantify changes in the slope of the RS neuron contrast response functions on control versus laser trials (Fig. 3A, inset). As suggested from the average responses, SST stimulation reduced the slope of contrast response functions in larger fractions of individual RS neurons than PV stimulation (Fig. 3A; *p* = 0.032). To control for potential confounds of unreliable slope estimates in weakly responsive neurons, we next examined only neurons with high firing rates (those firing more than the population median, >3 spikes/s). Again, SST stimulation evoked larger reductions in the slope of contrast sensitivity than PV stimulation (*p* = 0.010); further, a simple linear fit of the control versus photostimulation responses^40^ (across contrasts) also revealed a smaller y-intercept (more subtractive relationship) with PV versus SST stimulation (*p* =0.02). During SST stimulation, a large fraction of RS neurons (38%) became completely insensitive to contrast (MI = -1; Fig. 3B inset), a significantly greater fraction than during PV stimulation (21%; SST vs. PV, *p* = 0.021). The strong effects in these neurons were masked in population averages (not shown), and were not due to poor curve fits: constraining fits to enforce shallower slopes (Methods) did not reduce the fraction of MI = -1 neurons, nor reduce effects on contrast response function slope (SST: -0.74 ± 0.50 MI; PV: -0.47 ± 0.53 MI; *p* = 0.02). Likewise, the results were not due to overrepresentation of high firing rate contrast tuned neurons in single sessions: hierarchical bootstrapping^41^ that resampled evenly across mice and sessions showed that, if anything, hierarchical resampling evenly across the population reduced the inherent noisiness of individual neurons and sessions (minimizing the influence of the neurons with increased slopes during photostimulation), but still preserved greater reductions in the slope of contrast sensitivity for RS neurons during SST stimulation (Fig. 3C). Lastly, an affine model revealed that RS responses during SST stimulation were explained by a significantly greater change in the gain, while responses with PV stimulation were explained by a significantly greater change in offset (subtraction; Fig. S4), mirroring the perceptual responses.

**Figure 3.**
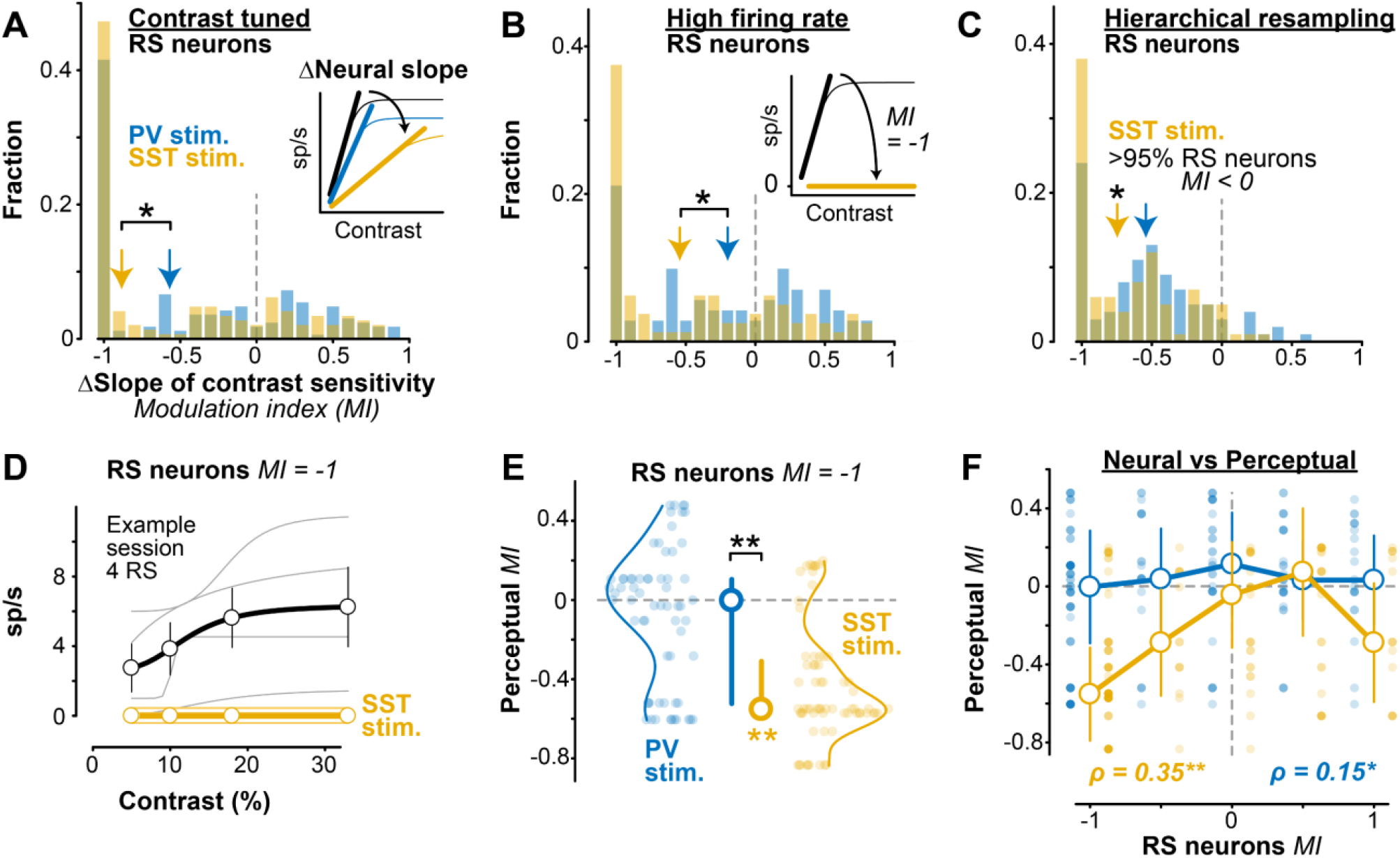
SST activation simultaneously reduces neural and perceptual contrast sensitivity. **A**. SST stimulation significantly decreased contrast response function slope (inset) in contrast tuned RS neurons (−0.88 ± 0.54 MI, median ± MAD, 146 RS neurons; *p <* 1e-13, Wilcoxon signed rank test), and significantly more than did PV stimulation (−0.57 ± 0.55 MI, *p <* 1e-11; 166 excitatory neurons; PV vs SST, *p =* 0.032, 1-tail Wilcoxon rank-sum test). **B**. Same as A for RS neurons with high firing rate (> 3 spikes/s, population median). Larger slope decrease with SST (−0.54 ± 0.53 MI, 80 RS neurons) versus PV stimulation (−0.20 ± 0.50 MI, 71 RS neurons; PV vs SST, *p = 0*.*010*). Inset shows schematic for neurons that become contrast insensitive (MI = -1) with SST stimulation (38% of neurons with SST stimulation; 21% of neurons with PV stimulation; *p =* 0.021, Fisher’s exact test). **C**. Hierarchical bootstrapping verified robustness of RS slope changes. SST stimulation significantly decreased the slope of RS neurons (−0.75 ± 0.31 MI, *p* < 0.05, >95% of bootstrapped samples had a decrease in slope), but distal PV stimulation did not significantly decrease the slope of RS neurons (−0.54 ± 0.31, *p* = 0.10). **D**. Response of 4 example simultaneously recorded RS neurons with a neural slope MI = -1 during distal SST stimulation. Spiking activity decreased to 0 across contrasts (individual curves offset from 0 for visualization). Corresponding behavioral slope MI = -0.57. Mean ± SEM. **E**. The corresponding perceptual MI for neural MI = -1 (68 RS neurons during SST stimulation, 64 RS neurons during PV stimulation). Significant decrease in perceptual MI with SST stimulation (−0.55 ± 0.24 MI, median ± MAD; *p < 1e-10*, Wilcoxon signed-rank test), but not PV stimulation (0.00 ± 0.29 MI; *p* = 0.12). Median ± IQR. **F**. Changes in neural and perceptual slope of contrast sensitivity more strongly correlated during SST stimulation (ρ = 0.35, *p* < 1e-4, Spearman’s rank correlation) than distal PV stimulation (ρ = 0.15, *p* = 0.048). Decoding perceptual performance from RS activity revealed stronger changes to psychometric sensitivity with SST stimulation (Fig. S8)

We further examined how loss of contrast sensitivity in RS neurons varied with perceptual effects. Remarkably, in sessions where SST stimulation evoked complete loss of contrast sensitivity in any single RS neuron (MI = -1, example in Fig. 3D), there was a simultaneous and significant decrease of perceptual contrast sensitivity (Fig. 3E; *p <* 1e-11), with no such effect during PV stimulation (*p* = 0.12). We expanded this approach to examine modulation of all contrast tuned RS neurons (not just those with total loss of contrast sensitivity). We found that there were significant session-by-session correlations between overall firing rate and overall d’ (Fig. S7). Moreover, SST stimulation drove significantly correlated reductions in both neural and perceptual contrast sensitivity slopes (Fig. 3F; ρ = 0.35; *p* < 1e-4), with a weaker relationship during PV stimulation (ρ = 0.15; *p* = 0.048). In other words, lateral SST inhibition drove stronger reduction in the gain of contrast tuning of RS neurons, and this also drove stronger reduction in the gain of perceptual sensitivity to the same stimuli on the same trials. Further, decoder analysis revealed that neural activity (all neurons, no tuning criteria) predicted behavioral performance, and recapitulated the larger changes in the slope of psychometric sensitivity during SST (but not PV) stimulation (Fig. S8); further, removing the neurons with total loss of contrast sensitivity (MI = 1) from the decoder abolished the difference in slope changes with SST versus PV stimulation. We next turned to a simple neural circuit model to probe potential mechanisms driving these effects of PV versus SST lateral inhibition.

### Network model with long-range SST projections replicates contrast tuning effects

We wondered if differing control of contrast sensitivity was due to the subcellular location of PV versus SST inhibition, or due to the larger spatial extent of SST versus PV projections, as suggested by recent experiments^42,43^. We constructed a leaky integrate-and-fire (LIF) circuit model of excitatory (E), PV, and SST neurons based on prior studies^44^ and updated this model in three key ways (Fig. 4A, B). First, we updated the inhibitory connectivity probabilities and weights to match recent experimental data^20,21^. Second, we approximated SST dendritic inhibition by including an interaction term between excitation and SST inhibition^45^; this approach captures arithmetic operations of dendritic inhibition in pyramidal neurons^46^. Third, we updated the spatial extent of SST projections to reflect greater spatial influence of SST inhibition^42,47^, particularly relevant in V1 where most SST neurons are Martinotti cells characterized by extensive lateral axons^48^. This V1 model was set up as a 1mm^2^ grid, with “visual” spiking (provided by an external thalamic layer) arriving in a 0.2mm region of the grid (Gaussian distribution, see Methods and Fig. S9 for spatial grid model properties). We simulated “optogenetic” activation in silico by injecting a conductance into PV or SST neurons ∼0.8mm away from the site receiving visual input. The model parameters were fit to the spontaneous and visually-evoked firing rates of single unit activity recorded in RS (putative E) cells and both PV and SST neurons (detailed later) in control conditions (Methods). This simple model replicated the overall experimental effects of lateral PV versus SST stimulation on E neurons (Fig. 4C, D; model at left, experiments at right). Furthermore, this model displayed many hallmarks of cortical network dynamics: inhibitory stabilization^49^ (Fig. S10), asynchronous irregular spontaneous activity^50^ (Fig. S11), and spontaneous firing variability that matched the data (Fig. S11).

**Figure 4.**
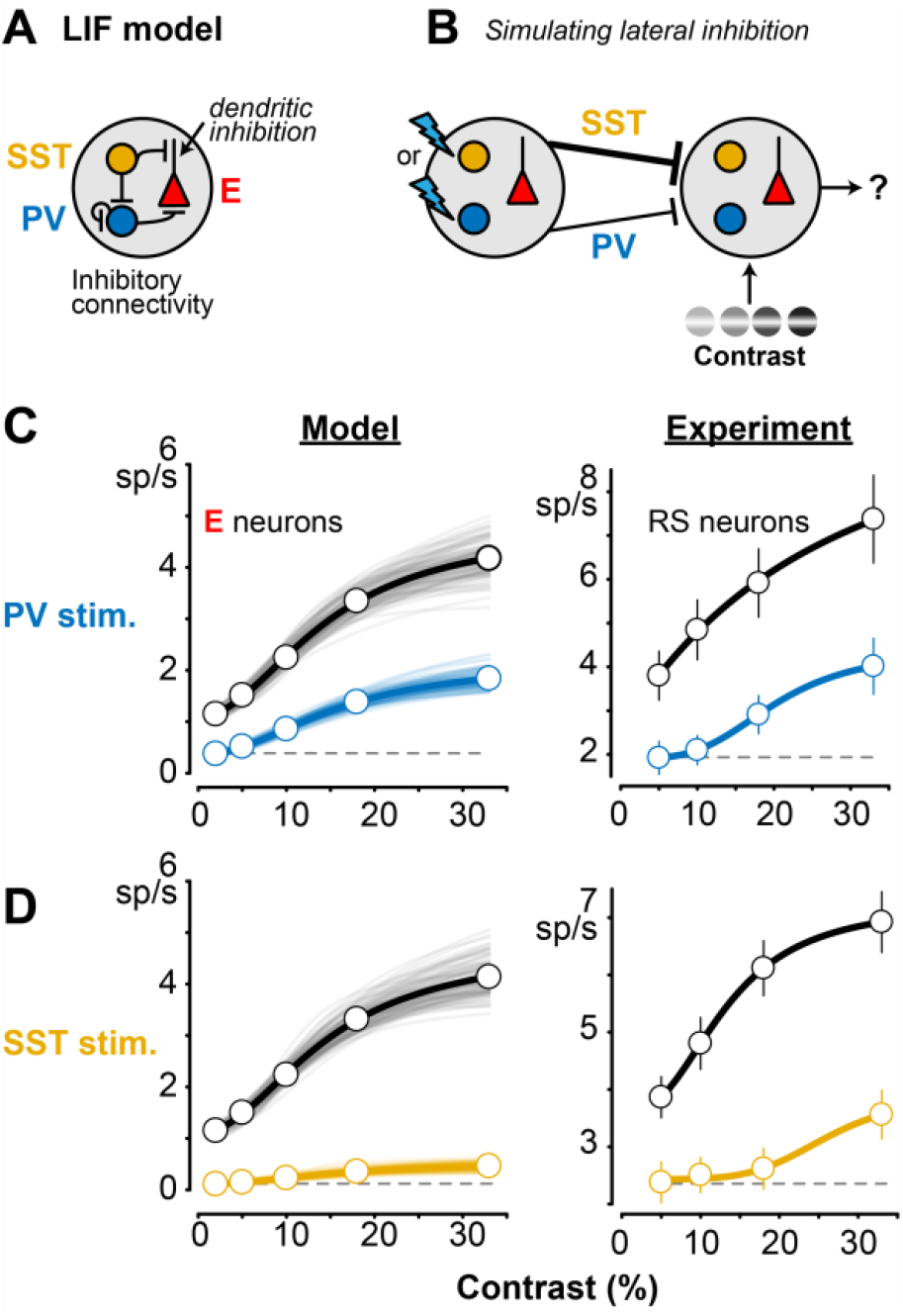
LIF network model replicates effects of PV or SST stimulation on contrast tuning. **A**. Conductance-based leaky integrate- and-fire (LIF) network model with PV, SST, and excitatory (E) neurons. SST dendritic inhibition of E neurons (arrow) modeled as multiplicative interaction with excitation per prior studies (see Results). Excitatory connectivity (Methods) not shown here to highlight inhibitory connectivity. **B**. Within a 1mm^2^ grid of neurons, a 0.2mm patch received contrast-modulated “visual” input from an external thalamic layer (right), while PV or SST neurons ∼0.8mm away were directly activated to mimic lateral inhibition (left). The model included a higher probability for long-range SST versus PV projections (thicker line), per experimental data (Results). **C**. Effects of distal PV stimulation in the model (left) and in the experiment (right, replotted from Fig. 2C). **D**. Same as C for distal SST stimulation. More divisive scaling of E responses in model and experiment, relative to PV stimulation.

Using this simple circuit model, we varied either the strength of SST dendritic inhibition, or the spatial extent of SST connectivity, to disentangle two potential mechanisms mediating the effects of SST lateral inhibition (Fig. 5A). We found that a higher probability of SST lateral connections strongly modulated the slope of contrast sensitivity (Fig. 5C, top to bottom), while the strength of dendritic inhibition played a smaller role (Fig. 5C, left to right). We ran multiple instances of the model in three fixed regimes (Fig. 5D-F), and found that a high probability of lateral SST connectivity drove the greater changes in the slope of contrast sensitivity (Fig. 5D), even when dendritic SST inhibition was weakened (Fig. 5E). However, with a low probability of lateral SST connectivity, strong dendritic inhibition by itself was unable to change the slope of contrast sensitivity during SST stimulation (Fig. 5F). Dendritic inhibition thus only moderately amplifies the effects driven by the greater SST connection probability, even at different stimulation distances (Fig. S12). In all of these model iterations, the effects of lateral PV stimulation were unaffected by changes in SST projection probability or dendritic strength (Fig. 5B, D-F). Furthermore, when we removed the long-range SST projections, and instead increased the local dendritic excitability of SST neurons, this by itself was unable to cause divisive changes in the slope (or gain) of the local RS neurons (Fig. S13). These findings identify long-range projections of SST neurons as a potentially critical factor for the effects of lateral inhibition on the gain of contrast sensitivity.

**Figure 5.**
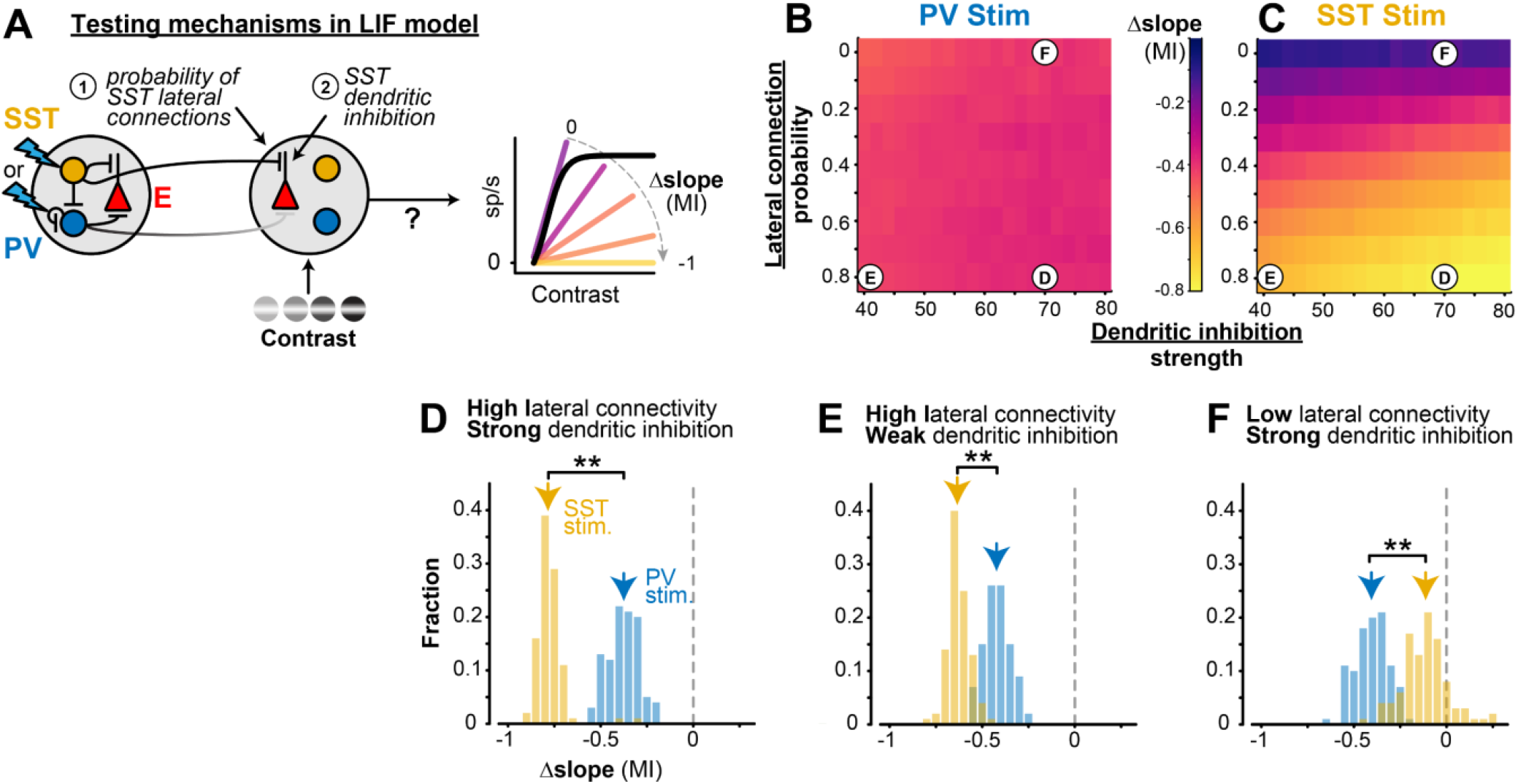
Long-range SST projections in LIF network model control slope of contrast sensitivity. **A**. The probability of SST lateral connections and SST dendritic inhibition strength was varied in the model to identify the source of the slope change with distal SST stimulation. **B**. Heat maps showing the average (100 iterations) slope MI of E neurons during distal PV stimulation for varied long-range SST projection probabilities (p^l^ _SST_) and SST dendritic inhibition strengths (α; see Methods). MI is constant during distal PV stimulation with any lateral connection probability or dendritic inhibition strength. Letters on heatmaps indicate various SST dendritic integration strengths and long-range SST probabilities depicted in D-F. **C**. Same as B for distal SST stimulation. Slope changes during distal SST stimulation are largely dependent on SST lateral connection probability (vertical gradient). Slope changes slightly varied with SST dendritic inhibition strength. **D**. For high SST lateral projection probability (p^l^ _SST_ = 0.8) and high SST dendritic inhibition strength (α = 70 MΩ), distal SST stimulation had larger divisive effects (−0.78 ± 0.05 MI, median ± MAD) than distal PV stimulation on E neurons across contrasts (−0.38 ± 0.07 MI; *p* < 1e-32, 1-tail Wilcoxon rank-sum test). **E**. For high SST lateral projection probability (p^l^_SST_ = 0.8) and low SST dendritic inhibition strength (α = 40 MΩ), distal SST stimulation has larger divisive effects (−0.63 ± 0.04 MI) than distal PV stimulation on E neurons (−0.42 ± 0.06; *p* < 1e-32). These effects are smaller than when the strength of SST dendritic inhibition is high (D). **F**. For low SST lateral projection probability (p^l^_SST_ = 0) and high SST dendritic inhibition strength (α = 70 MΩ), distal SST stimulation had a smaller divisive effect (−0.11 ± 0.09 MI) than distal PV stimulation (−0.41 ± 0.08 MI; *p* < 1e-30), indicating long-range SST projections are essential for eliciting strong divisive effects.

We next anatomically confirmed extensive lateral projections from SST but not PV neurons, using multiple approaches. First, we analyzed the Allen Brain Atlas mouse V1 single neuron morphology database^51^ (*in vitro* patch-seq recordings). Here, SST neurons showed significantly more extensive lateral axonal projections than PV neurons, with many SST neuron axons extending 0.4mm-0.8mm laterally (Fig. 6); importantly, axonal spread was much greater than dendritic spread. Second, in another study of mouse V1 SST neurons^48^, we found several examples of Martinotti cells with extensive axonal projections (>0.4 mm laterally), consistent with the findings above (Fig. S14). Third, we performed anatomical tracing of SST and PV axonal spread laterally across the V1 retinotopic map of horizontal visual space. We targeted viral injections to the same monocular region of V1 that was illuminated in behavioral experiments (Fig. S15), and found SST axons extending laterally >0.4mm (and up to 1mm) into the binocular regions of V1 (the site of stimulus drive); this was not the case for PV axons, which remained confined to the injection site. (Fig. S16). Due to tissue shrinkage and axonal slicing, the projection distances *in vitro* are likely smaller than the actual distances *in vivo*, but the relative differences between PV and SST neurons are unaffected. Together, these findings provide multiple lines of anatomical evidence that SST axons are more laterally widespread than PV axons across V1, and could thus mediate the greater divisive inhibitory effect of SST inhibition, as predicted from the model.

**Figure 6.**
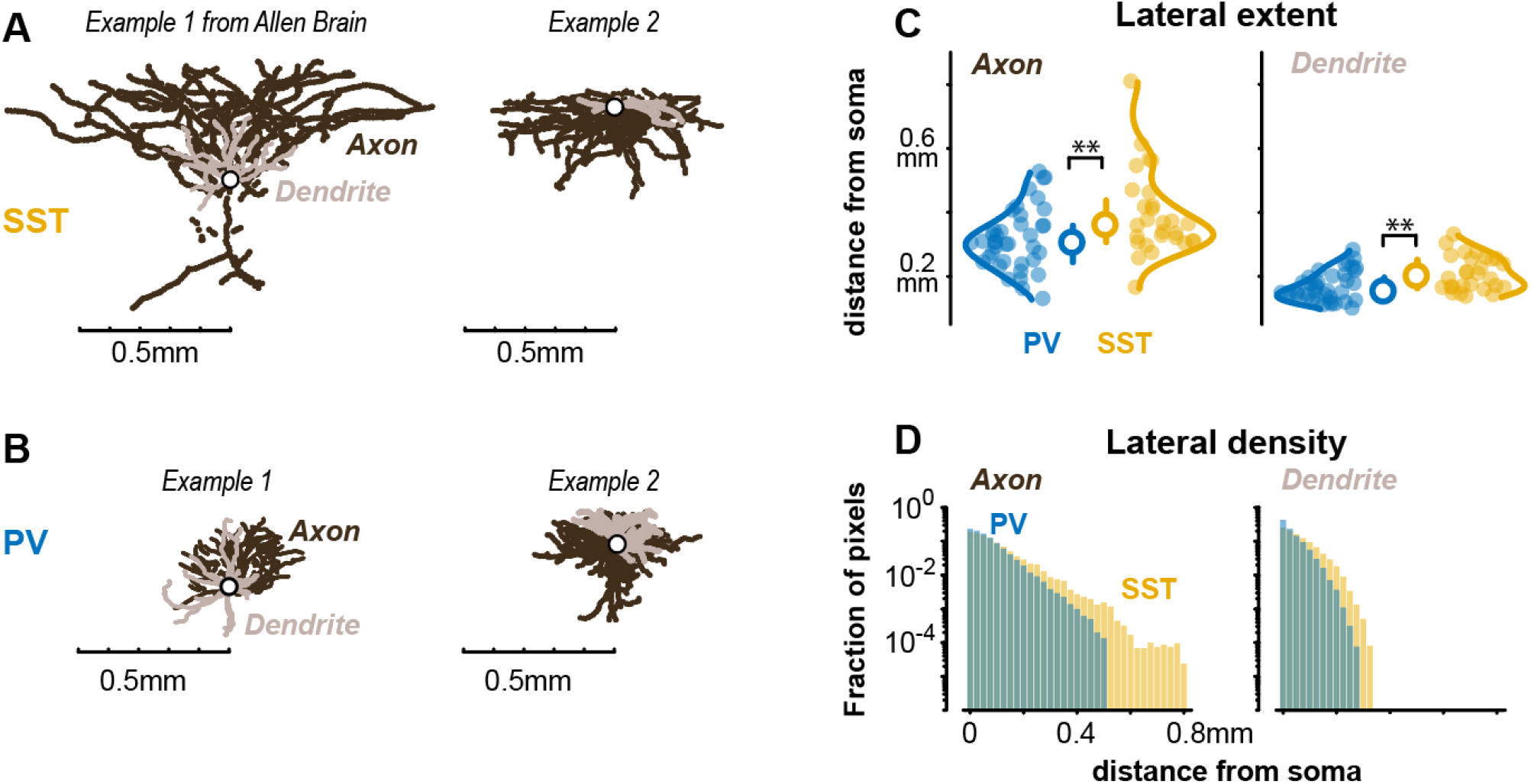
More extensive lateral axonal projections in SST than PV neurons. **A**. Two example mouse V1 SST neurons reconstructed from *in vitro* patch clamp recordings (from Allen Brain Atlas Cell Types database^51^). Scale bar aligned to soma. **B**. Same as A, for two PV neurons. **C**. *Left*, the maximum lateral extent of axonal projections is significantly greater in SST (n=28) versus PV (n=39) neurons. SST = 0.36 ± 0.10 mm, PV = 0.30 ± 0.07mm (median ± MAD; *p* < 0.01, Wilcoxon rank sum test). *Right*, SST neuron dendrites extend laterally more than PV neurons (SST = 0.20 ± 0.05 mm, PV = 0.15 ± 0.04 mm, *p* < 0.01, Wilcoxon rank sum test). **D**. SST neurons had greater axonal and dendritic lateral density than PV neurons (*p* < 0.01, Kolmogorov-Smirnov test). In both cell types, axons projected significantly greater distances than dendrites (*p* < 1e-8) by nearly 200 µm. See S14 and S16 for further anatomy.

Do we find functional evidence for a larger spatial footprint of SST activity? We reasoned that if SST neurons have a larger spatial extent of projections than PV neurons, SST neurons may spike more readily when stimulated at sites distal to the recording, potentially by direct activation of long-range axons^52-55^. We tested these predictions in awake mice (outside of the task) by first identifying PV and SST neurons with brief square pulses (0.04s) of light directly at the recording site (“optotagging”^56^; Fig. S5), then moving stimulation to a distal site (0.8mm away, as in behavioral experiments). Local stimulation in PV-ChR2 mice rapidly increased population activity of fast-spiking (FS) putative PV inhibitory neurons (identified by waveform) and suppressed RS activity (Fig. S17A). The majority of FS neurons (n = 368) were also significantly optogenetically driven (n = 203 tagged PV+), while the statistically non-driven FS units (n = 165) were likely PV neurons that were themselves suppressed by driven PV units (Fig. S5D), consistent with strong self-inhibition among V1 PV neurons^20,21^. Local stimulation in SST-ChR2 mice identified significantly driven SST units (n = 52), and this strongly suppressed both RS (n = 784) and FS (n = 177) neurons (Fig. S17B). These effects on firing rates were present at the laser power used during the behavioral task, and increased at higher laser intensities at the recording site (Fig. S18). Importantly, the raw light-driven firing rates and latency to peak were not different between PV and SST neurons, suggesting similar somatodendritic photo-excitability (Fig. S19).

When we stimulated at the distal site far away from the recording, in SST-ChR2 mice we measured significant increases in local spiking of SST+ units at the recording site (Fig. S17D; Fig. S18D). On the other hand, in PV-ChR2 mice, we never observed local FS units that increased spiking with distal stimulation (Fig. S17C; Fig. S18C). The increase in local SST activity with brief square laser pulses at a distant V1 site (0.8mm away) is consistent with antidromic activation of SST neurons that have long-range axonal projections (Fig. S20), since the dendrites of both PV and SST neurons at the recording site extend only ∼0.2mm (Fig. 6). Importantly, RS (presumably excitatory) units were always suppressed at the recording site in these experiments, and both RS and FS neurons were always suppressed with SST stimulation during the task (Fig. S21), indicating the overall suppressive effect of lateral inhibition with no evidence of disinhibition of RS neurons. These overall suppressive effects of strongly driving the SST populations at the distal site could be due to withdrawal of excitation, increased inhibition, or a combination of both^9,10,16,53^. We next sought direct evidence for changes in synaptic excitation and inhibition in excitatory neurons during distal activation of SST versus PV neurons.

### Synaptic inhibition from SST neurons underlies changes in contrast sensitivity

Whole-cell patch-clamp recordings revealed that distal SST stimulation evoked greater hyperpolarization than PV stimulation, and this was due to elevated inhibitory conductance in excitatory neurons. We performed both current and voltage clamp recordings in awake mice since anesthesia suppresses SST neuron activity^15^ and severely compromises the spatial and temporal functions of cortical inhibition during awake visual processing^57-59^. We first found that distal SST stimulation evoked significantly stronger hyperpolarization of membrane potential than distal PV stimulation (Fig. 7A-C). These measurements were biased towards L2/3 neurons, which have low spontaneous firing rates; thus, resolving the effects of distal stimulation solely from spiking could be subject to a “floor effect” (rates can’t go below zero). Indeed, silicon probe recordings of spontaneous activity in L2/3 showed PV and SST stimulation decreased spike rates to a similar degree at these laser intensities (*p* = 0.62; Fig. S18). Importantly, subthreshold measurements are not subject to this floor effect, and reveal clear differences in the underlying synaptic input. We then performed voltage clamp experiments to determine if this greater hyperpolarization was due to reduced excitation, elevated inhibition, or a combination of the two. Distal PV or SST stimulation both reduced excitatory conductance, but to a similar degree (Fig. 7D). However, there was significantly greater inhibitory conductance evoked by distal SST versus PV stimulation (Fig. 7E), despite dendritic SST inhibition being further from the patch pipette than somatic PV inhibition. These effects were not due to differences in recording quality or electrical access in PV versus SST recordings (Methods). Within each neuron, the net suppression (ΔG_e_ – ΔG_i_) was significantly more intense with distal SST (−3.7 ± 1.5 nS) than PV stimulation (−1.9 ± 1.2 nS; *p =* 0.0310, Kolmogorov-Smirnov test). This combination of conductance effects likely underlies the stronger membrane potential hyperpolarization seen with distal SST stimulation. Taken together, these subthreshold recordings identify withdrawal of excitation and elevated synaptic inhibition in local excitatory neurons as a potential mechanism to adjust the slope of contrast sensitivity. We returned to the LIF model to test if directly modulating synaptic inhibition controls the slope of the contrast sensitivity in excitatory neurons.

**Figure 7.**
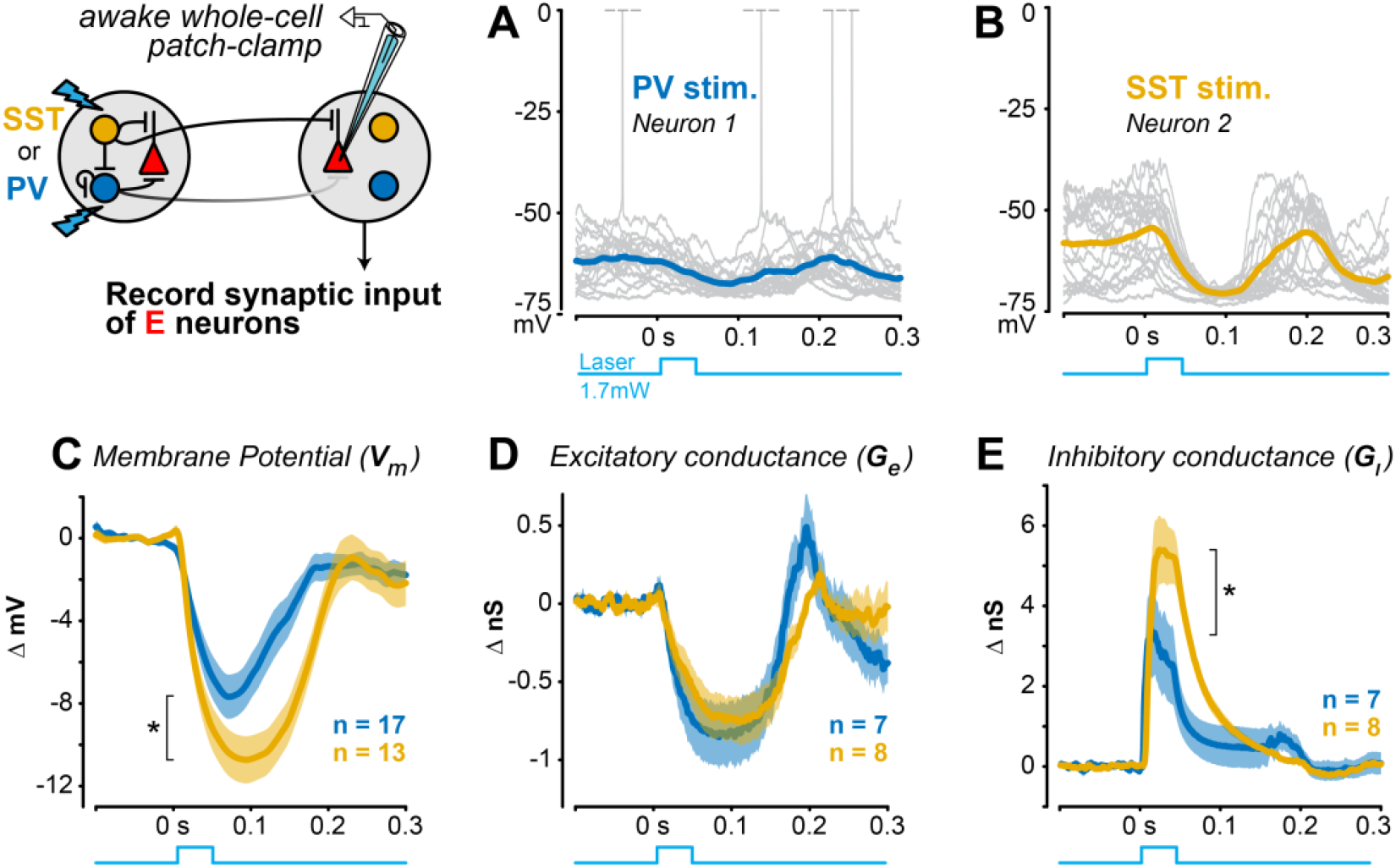
Stronger lateral inhibition from distal SST versus PV neurons. **A**. Example current clamp recording of membrane potential from an RS neuron during distal PV stimulation in awake V1 (20 trials). Spikes truncated at 0 mV. **B**. Same as A for an example RS neuron during distal SST stimulation. **C**. Distal SST stimulation causes greater hyperpolarization of excitatory neurons (ΔV_m_ = -7.72 ± 0.91 mV, mean ± SEM, 13 RS neurons; mean ΔV_m_ from 0 – 0.2s) than distal PV stimulation (ΔV_m_ = -4.52 ± 0.76 mV, 17 excitatory neurons; *p =* 0.008, 1-tail Wilcoxon rank-sum test). **D**. Distal PV or SST stimulation decreased excitatory conductance to a similar degree (PV stim: ΔG_e_ = -0.74 ± 0.19 nS, 7 RS neurons; SST stim: ΔG_e_ = -0.54 ± 0.13 nS, 8 RS neurons; *p* = 0.198; mean ΔG_e_ from 0 – 0.1s). **E**. Inhibitory conductance significantly larger with distal SST versus PV stimulation (SST stim: ΔG_i_ = 3.07 ± 0.40 nS, 8 RS neurons; PV stim: ΔG_i_ = 1.15 ± 0.52 nS, 7 RS neurons; *p =* 0.0047; mean ΔG_i_ from 0 – 0.1s).

We found that the intensity of inhibition in the V1 circuit model was the key factor driving changes in slope of contrast sensitivity of E neurons. We systematically varied the effective intensity of inhibition by moving the site of “optogenetic” stimulation in silico. During local stimulation directly at the site of “visual” input (where both PV and SST inhibitory connections to E neurons were prevalent), the slope of contrast sensitivity strongly decreased with stimulation of either PV or SST neurons (Fig. S22). However, as we moved the stimulation site towards distal locations, the effects of PV stimulation on the slope of contrast sensitivity rapidly decreased since the lateral connectivity of PV neurons to local E neurons also decreased. On the other hand, distal SST stimulation retained the ability to decrease the slope of contrast sensitivity because of the stronger lateral connectivity of SST to E neurons (Fig. S22).

We performed several control experiments and simulations to rule out potential “off target” photoexcitation of local SST neurons as a main factor underlying these effects. In one set of experiments, we expressed opsin only at the distal site with focal viral injections (as opposed to transgenic expression): this minimizes “off target” activation of the local SST neurons via their axons (or light spreading to dendrites); reassuringly, these experiments replicated the divisive gain change at the local site with distal SST (but not PV) stimulation (Fig. S23). Next, we similarly expressed opsin only at the local site, and measured the effects of distal photostimulation. This only activated local SST (not PV) neurons, and likewise only suppressed local activity in SST (not PV) mice; importantly, the magnitude of local suppression in these experiments (presumably mediated solely by the local SST neurons) was much smaller than when photostimulation strongly drove the SST populations directly at the distal site (Fig. S20). If these effects were simply because of light spread to the local site, then local PV neurons with opsin would also be driven (they were not). Lastly, we used the LIF model to restrict photostimulation only to SST neurons with cell bodies at the distal site (removing the small fraction of axonally activated neurons from the original model) – this caused no decrement in the magnitude of local divisive gain change (Fig. S24). We then simulated a greater spread of light (by activating more SST neurons towards the local site), but in the absence of greater SST lateral projections. This increase in SST population activation by itself was unable to cause divisive gain changes at the local site — these only emerged when we restored the higher probability of lateral SST connections.

The model makes a strong prediction: the effects of local PV and SST inhibition should be comparable, and both should cause divisive gain changes in E neurons (Fig. S22). We confirmed this with several lines of experiments. First, local optogenetic stimulation of either PV or SST neurons (at the recording site driven by the visual stimulus) showed similar amounts of photoexcitation (Fig. S19), and both decreased the contrast sensitivity slope of RS neurons, to a similar degree (Fig. S22E, S26C-D). Second, local stimulation of PV or SST neurons evoked similar amounts of hyperpolarization and inhibitory conductance (Fig. S25). Third, as predicted from the model, local PV stimulation significantly decreased the slope of both neural (–1 ± 0.41, median ± MAD) and perceptual contrast sensitivity (−0.53 ± 0.41), much more than distal PV stimulation did (neural: -0.57 ± 0.55, Fig. 3A; perceptual: -0.03 ± 0.20, Fig. 1D;), and to a similar degree as local SST stimulation (Fig. S22E; S26). These results provide a stark contrast to the effects of distal SST or PV stimulation, and from the exact same experimental conditions.

Taken together, the model and experiments show that when excitatory neurons receive strong inhibition –regardless of the presynaptic source– this divisively scales the neural and perceptual sensitivity to contrast. However, SST neurons possess a unique ability to modulate contrast sensitivity across large regions of cortical and visual space, by virtue of their more extensive lateral inhibitory footprint.

## Discussion

Here we have established that lateral inhibition in V1 exerts direct effects on neural and perceptual sensitivity to stimulus contrast. Lateral inhibition from SST neurons caused greater divisive scaling of perceptual response curves than did PV lateral inhibition. The perceptual effects of SST lateral inhibition were correlated with the strength of divisive modulation in putative excitatory neuron responses on those same trials. A simple circuit model predicted long-range SST inhibition as a key driver of these effects, as confirmed with direct measurement of a larger spatial footprint of SST versus PV synaptic inhibition. Taken together, our findings establish a mechanistic basis for cell-type specific computations performed by lateral inhibition in V1. These operations flexibly modulate behavioral sensitivity to contrast across large regions of the visual field.

Driving PV and SST neurons far from the site of visual input altered perception in unique ways. PV lateral inhibition uniformly reduced contrast sensitivity but preserved its steepness. SST lateral inhibition more effectively scaled down sensitivity to low and medium contrast, linearizing the dynamic range^60^. Importantly, the behaviorally relevant visual stimulus did not itself confound the effects of driving distant PV or SST neurons. These two modes of lateral inhibition provide flexible ways to control behavioral sensitivity to visual contrast spanning extensive retinotopic space (∼70° in our experiments). On the other hand, driving either SST or PV neurons directly at the retinotopic site of visual input (where both SST and PV connections to excitatory neurons are dense) invariably caused divisive effects on perceptual responses, consistent with prior studies of local inhibition on contrast perception^61^ and neural selectivity^25,28,62,63^. Although some studies have shown that local SST inhibition preceding visual input can cause subtractive effects on V1 firing rates^27,64^, divisive effects on firing rates dominate when visual excitation and local SST inhibition overlap in time^25,26^, as was the case here. Taken together, our results show how the perceivability of visual stimuli can be adjusted with distinct computations via cell-type specific lateral inhibition. These operations in V1 could be readily accessed by top-down inputs.

The cell-type specific effects of lateral inhibition depended on its spatial footprint. Using a V1 circuit model, we found that the critical feature driving greater divisive scaling of excitatory firing by SST rather than PV neurons was the reach and strength of inhibition, not its subcellular location (SST dendritic inhibition vs PV somatic inhibition). Preserving lateral projections of SST neurons but eliminating their dendritic effects still produced greater divisive scaling of excitatory neuron firing. We confirmed this critical feature with direct measurements of hyperpolarization driven by synaptic inhibition from distal SST neurons in awake mice. Further, we were able to resolve these differences even though dendritic SST inhibition is electrotonically attenuated compared to somatic PV inhibition. Our direct measurements of a larger spatial footprint for SST inhibition provide experimental confirmation of theoretical predictions^19^. Three additional pieces of experimental evidence support our findings. First, SST neurons in V1 fire most strongly to large visual stimuli (>70°)^15^, and are involved in synchronizing long-range (>0.5mm) visual oscillations^65^, both consistent with the spatial scale of effects on perception shown here. Second, most SST neurons in V1 are Martinotti subtypes defined by their extensive lateral axonal arbors^20,48^. Third, in slices of auditory cortex, the functional spatial footprint of SST inhibition also extends further than PV inhibition, and this lateral inhibition underlies the sharpness frequency tuning^42^. To our knowledge, our findings are the first to directly measure the subthreshold extent of cell-type specific lateral inhibition in awake V1, and identify this as a key mechanism enabling adjustment of perceptual sensitivity.

Lateral inhibition in mouse V1 suppressed firing of local excitatory and PV neurons, consistent with stabilized supralinear network architecture^16,49,66-68^. Our measurements of both cell-type specific spiking and subthreshold synaptic activity provide deeper insights for this framework. Activating distant PV neurons suppressed excitatory neuron firing and elicited a “paradoxical” decrease in the firing of local PV neurons, likely due to the combined loss of local excitation and increase of direct inhibition among PV neurons^16,21^. However, subthreshold recordings revealed that despite a relative lack of extensive lateral spread of PV axons, and without local PV neuron spiking, distal PV activation still hyperpolarized local excitatory neurons. We speculate that specializations in PV networks (gap junction and synaptic coupling, strong synchronization, strong directed connectivity with excitatory neurons^69^) may have additional network consequences (via effects on reduced lateral excitation rather than direct inhibition), a topic for future work. SST lateral inhibition also strongly suppressed both local excitatory and PV neuron firing, but local SST neurons were not suppressed, consistent with connectivity data^20^. We performed several control experiments and simulations that showed the main effects of SST stimulation were due to the more extensive lateral projections of SST populations at the distal site, causing greater inhibition of local excitatory neurons, and greater scaling of perceptual contrast sensitivity. These findings highlight that lateral interactions in networks with realistic connectivity and multiple inhibitory neuron subtypes can support both “paradoxical” suppression of some inhibitory neurons but activation of others^70^, leading to differing impact on stimulus responses and behavior.

Our study focused on PV and SST neurons for several reasons. First, they comprise ∼70% of cortical inhibitory neurons^21,71^, and constitute the strongest sources of direct synaptic inhibition to cortical excitatory (E) neurons in mouse V1^20^. Second, numerous studies have established computational transformations of visual responses in E neurons by local PV and SST inhibition^18,19,22,25-28,38,44^, but with far less known about their lateral interactions and consequences for visual perception. For these reasons, our study focused on measuring the direct effects on subthreshold and spiking responses in E neurons, along with their perceptual consequences. Now that we have established that SST lateral inhibition plays a key role for divisive gain modulation, it will be important to examine if VIP interneurons gate and shape SST lateral inhibition. VIP interneurons comprise ∼15% of cortical interneurons^21,71^, but respond to small stimuli^19,72^ as shown in our task, and they preferentially inhibit SST neurons, thus influencing cortical dynamics and visual responses via their “disinhibitory” effect on E neurons (reviewed by^73^). However, interactions between SST and VIP cells in V1 could be much more bidirectional than previously thought^20^. The role of VIP interneurons for the effects described here is an important topic for future experiments; these should monitor spatially extended populations of these sparse cell types simultaneously with SST and E neurons; likewise, future computational models that investigate spatially extended cortical inhibition should incorporate both local and distant connectivity and synaptic dynamics of multiple interneuron types.

Our findings carry some limitations that can be addressed with future investigations. Firstly, we focused here on the divisive aspects of modulation (changes in slope) because these have been most consistently implicated in many studies of local inhibition and gain modulation^29^. Our study revealed that SST rather than PV lateral inhibition causes relatively greater divisive modulation. However, this does not imply that SST lateral inhibition is *purely* divisive, or that PV lateral inhibition is *purely* subtractive – both aspects co-occur, as we show here, but the relative contributions of divisive and subtractive modulation show cell-type specificity. Further, a change in psychometric slope (scaling the dynamic range) is just one aspect of response modulation that may led to perceptual consequences, and likely other aspects contribute to neural^22,28,31^ and perceptual^61^ modulation, an important topic for future studies. Second, for experimental clarity we drove distant PV or SST neurons optogenetically, but without also driving them visually. Future work could explore effects of suppressing lateral inhibition^40^ as well as detailing its recruitment during spatially extensive visual stimulation. A related limitation is that photostimulation in transgenic-expressing mice necessarily drove synchronous activation of all opsin-expressing cell bodies and neurites at the distal site; importantly, our control experiments with focal expression at the distal site replicated the main results, but it remains important to determine how SST lateral inhibition evoked optogenetically is impacted by the specific spatial and temporal patterns of photoactivation. Finally, it will be interesting to determine the spatial scale of lateral projections from SST neurons across different cortical areas (including Martinotti and Chodl/nNOS subtypes^74,75^), and to examine how lateral interactions are recruited by cortical and subcortical inputs in a variety of visual spatial tasks.

## Acknowledgements

We thank members of the Haider lab for feedback, Anthony Lien and Pilar Rubio Beltran for help with viral injections, and the anonymous reviewers for constructive suggestions. This work was supported by the Alfred P. Sloan Foundation’s Minority Ph.D. (MPHD) Program Fellowship (to J.D.R.), the National Institute of General Medical Sciences (T32GM142616 to Z.M.), the National Eye Institute (R00 EY030840 to H.C.), Alfred P. Sloan Foundation Fellowships in Neuroscience (to B.H and H.C.), the Whitehall Foundation (to B.H.), National Institute of Neurological Disorders and Stroke and NIH BRAIN Initiative (NS107968, NS109978 to B.H.), and the Simons Foundation (SFARI 600343, B.H.).

## Author Contributions

J.D.R., A.J.O., K.P., B.W., K.W., L.T.B., and L.L. trained mice; J.D.R., A.J.O., K.P., and B.W. performed optogenetic experiments; B.W. designed and implemented optical hardware; J.D.R and K.P. performed silicon probe recordings with optogenetics; S.C. and B.H. performed patch clamp experiments; A.D.C.V. processed pupil data; J.D.R. wrote code and analysed data; S.H.K., Z.M., and H.C. designed the circuit model and performed simulations; S.H.K. and Z.M. wrote the modelling methods; J.D.R., B.H. wrote the manuscript with feedback from all authors.

## Inclusion and Diversity Statement

We worked to ensure sex balance in the selection of non-human subjects. One or more of the authors of this paper self-identifies as an underrepresented ethnic minority in science. One or more of the authors of this paper received support from a program designed to increase minority representation in science. The author list of this paper includes contributors from the location where the research was conducted who participated in the data collection, design, analysis, and/or interpretation of the work.

## Methods

### Experimental model and subject details

All experimental procedures were approved by the Institutional Animal Care and Use Committee (IACUC) at the Georgia Institute of Technology.

### Subject details

B6 PV[cre] (RRID: IMSR_JAX:017320) or Sst-IRES-Cre (RRID: IMSR_JAX:013044) mice crossed with Ai32(RCL-ChR2(H134R)/EYFP) (RRID: IMSR_JAX:024109) mice (crossed mice referred to as PV-ChR2 or SST-ChR2 respectively in this study) were used to optogenetically activate either PV or SST inhibitory neurons. Mice were individually housed under reverse light cycle and bred in house.

### Implant surgeries

Details of headplate / cranial window implants are described previously ^34,35,37,76^. Briefly, 4–10-week-old male and female SST- or PV-ChR2 mice were chronically implanted with a stainless steel headplate with recording chamber (11mm diameter) and a cranial window (5mm diameter, intact skull prep) during isoflurane anesthesia (3% induction, 1-2% maintenance). The cranial window was placed over the visual cortex to map the retinotopy and visual cortices through hemodynamic measurements (see Intrinsic Signal Imaging). Mice recovered for 3 days after implantation before experimentation.

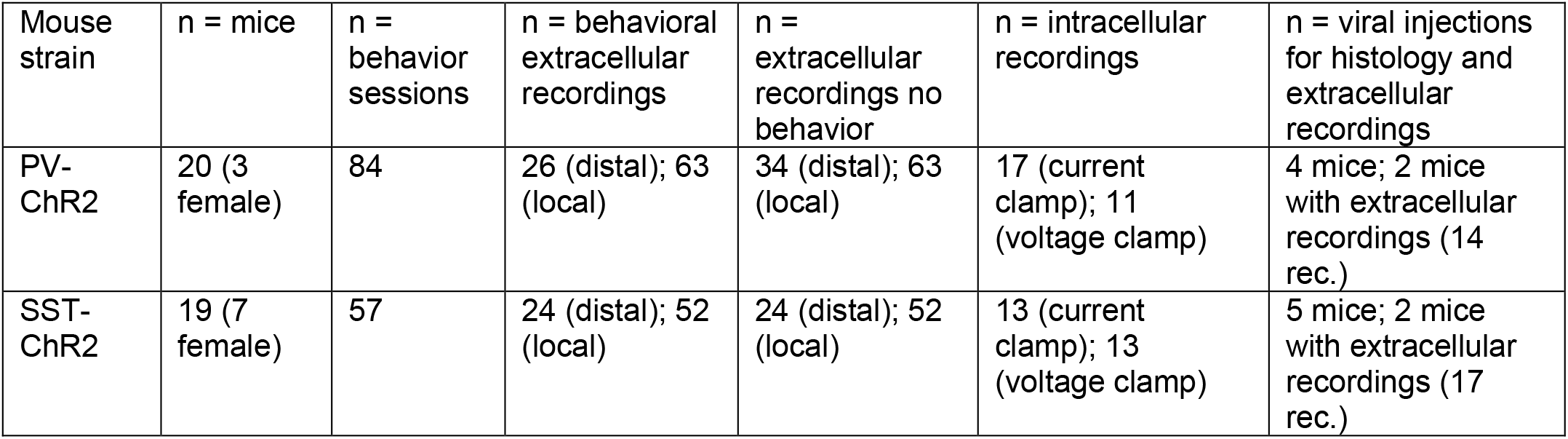

| Mouse strain | n = mice | n = behavior sessions | n = behavioral extracellular recordings | n = extracellular recordings no behavior | n = intracellular recordings | n = viral injections for histology and extracellular recordings |
| --- | --- | --- | --- | --- | --- | --- |
| PV-ChR2 | 20 (3 female) | 84 | 26 (distal); 63 (local) | 34 (distal); 63 (local) | 17 (current clamp); 11 (voltage clamp) | 4 mice; 2 mice with extracellular recordings (14 rec.) |
| SST-ChR2 | 19 (7 female) | 57 | 24 (distal); 52 (local) | 24 (distal); 52 (local) | 13 (current clamp); 13 (voltage clamp) | 5 mice; 2 mice with extracellular recordings (17 rec.) |

### Intrinsic signal imaging

Intact intrinsic signal imaging (ISI) method details are described previously ^37^. Briefly, mice were anesthetized (0.7-1% maintenance) and sedated (10^-5^ mg/kg Chlorprothixene) then head-fixed and positioned in front of 2 monitors spanning 150° (horizontal) by 48° (vertical) of the visual field. Contact lenses (3 mm diameter) were inserted to maintain ocular clarity during imaging. A camera was positioned over the cranial window and focused to ∼0.5 mm below the vasculature. Green (525 nm) and red (700nm) light were used to image vasculature and hemodynamic responses to visual stimuli drifting across the visual field. Azimuth (horizontal) and elevation (vertical) retinotopic maps were constructed from the hemodynamic responses. The visual field sign map was calculated from the sine angle difference between the azimuth and elevation maps, and used to delimit the extent of primary visual cortex (V1). Subject-specific retinotopic maps of V1 were used to target extracellular electrophysiological recordings and optogenetic stimulation. Optically targeted sites were confirmed via electrophysiological receptive field mapping (See “Recordings: Receptive field mapping”).

### Retinotopically targeted laser stimulation

Prior to laser stimulation experiments, mice were briefly anesthetized and a small cortical site (0.1 – 0.3mm) was either thinned to translucency, or opened into a small craniotomy over a retinotopically targeted site in V1. Mice recovered for >3 hours, and the skull was sealed between experiments by covering the site with an elastic polymer (Kwik-cast).

A blue laser (∼473 nm) was used for optogenetic stimulation in PV- or SST-ChR2 mice. Using custom optics, the laser was focused to a spot that restricted most of the power to the target site (Gaussian profile, 0.33mm full-width at half-max with 1.7mW power). Areas around the thinned skull / craniotomy were covered with opaque polymer (Kwik-cast) or dental cement (Metabond) to further restrict light spread.

A galvanometer precisely positioned the laser spot to retinotopic locations of V1 using ISI maps aligned to vasculature landmarks. In most experiments we stimulated V1 far away from the representation of detected stimulus (stimulus at 0°, laser at 70°; 0.8mm apart; Fig. 1). This experimental design isolates the effects of lateral inhibition from stimulus-driven activity. In a few experiments we also tested effects of laser stimulation directly at the recording site / V1 site representing the visual stimulus (Figs. S3-S5, S7, S8). Details of optogenetic stimulation during the behavioral task are described below (“Visual detection task with optogenetic perturbation”).

### Identification of laser-activated PV and SST neurons

After every electrophysiological experiment the laser was positioned directly over the recording site to measure single unit responses to brief (∼40 ms duration) laser pulses. This allowed us to statistically identify ChR2 expressing PV or SST neurons (“optotagging”^56^; Fig. S5). In some experiments, these brief pulses were also repeated at the distant manipulation site (Fig. S17; Fig. S18). Laser intensity in optotagging experiments was 0.5, 1.7, or 6.5 mW at the surface of the skull. These experiments allowed us to ensure that the moderate power we used during behavioral experiments (1.7 mW) avoided “paradoxical” or disinhibitory effects^49,77^ while also only modulating performance, rather than abolishing it (as we^34,35^ and others^78-80^ have shown with higher laser powers).

### Visual detection task with optogenetic perturbation

Water-restricted PV- and SST-ChR2 mice were trained to report visual detection by licking for water rewards (as detailed in prior studies^34,35,76^.Mice were head-fixed and stationary inside a plastic tube in front of 2 monitors spanning 160° in azimuth (from -37.8° to 115.8°; vertical meridian defined as 0°) and 48° in elevation (from -18.6° to 29.4°; horizontal meridian defined as 0°). Visual stimuli (static Gabor grating, *σ* = 8-12°) appeared without cueing and only after an enforced period of no licking (randomized per trial and drawn from a uniform distribution spanning 0.5s -7s). The first lick during grating presentation (1s response window) triggered water delivery. Visual stimuli appeared at a single location in blocks of 10 – 30 trials, either in the binocular (0°) or the monocular (70°) visual field, the same visual spatial detection task as our prior studies. Our main analysis here was restricted to blocks of trials with binocular stimuli and recordings in binocular V1, so that we could assess the modulatory effects of lateral inhibition on visual detection in the region of greatest visual sensitivity; however, we also analyzed detection of the monocular stimuli, and effects of PV or SST stimulation in monocular V1 during those trials (Fig. S22, S26). Michaelson contrast of the binocular stimuli ranged from 1-33% (typically sampling 4 contrasts) to capture the psychometric performance curve. 0% contrast stimuli were used to probe the false alarm rate. On a fraction (25 – 33%) of trials PV or SST neurons in monocular (∼70°) V1 were activated using a blue laser (∼473 nm, 1.7mW measured at the surface of the skull). The laser started ramping 0.1s before the appearance of the visual stimulus, and reached peak intensity at visual stimulus onset. The laser persisted throughout the duration of the visual stimulus. Visual stimuli (and laser stimulation, if present) terminated at the first lick on correct trials, or at the end of the response window (1 s) on incorrect trials. We analyzed the perceptual effects of optogenetic perturbations through either a thinned skull (6 PV-ChR2 mice, 30 sessions; 4 SST-ChR2 mice, 19 sessions) or craniotomy during recordings (6 PV-ChR2 mice, 54 sessions; 6 SST-ChR2 mice, 38 sessions). We observed no difference in the changes in psychometric slope of the contrast response curve for the thinned skull versus craniotomy preparation for distal SST stimulation (thinned skull, slope MI = -0.15 ± 0.31, median ± MAD; craniotomy, slope MI = -0.15 ± 0.31; *p* = 0.37, 1-tail Wilcoxon rank-sum test), or distal PV stimulation (thinned skull, slope MI = -0.16 ± 0.20; craniotomy, slope MI = -0.01 ± 0.19; *p* = 0.19). Behavioral data from all sessions were therefore combined in Fig. 1.

Perceptual performance analysis was primarily restricted to detection of stimuli presented in the binocular visual field to probe how PV or SST activation in site in V1 (monocular V1) not receiving task-relevant visual stimuli may influence perceptual behaviors. Perceptual performance was quantified using the sensitivity index d’ and response criterion (c) ^81^:

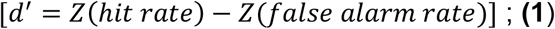

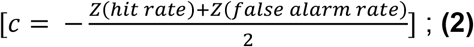

where Z represents the inverse of the normal cumulative distribution (MATLAB function, norminv). Task-irrelevant probe stimuli (5% contrast bars, 0.1s duration, 0.3s ISI, 9° width, appearing across randomly selected azimuth positions) were presented throughout the session, but not analyzed here; probe stimuli did not affect perceptual performance^35^. Mice typically performed hundreds of trials of detection in multiple spatial locations per day (307 ± 9 trials, mean ± SEM).

### Pupillometry analysis

We simultaneously recorded pupil position and diameter (a proxy for arousal ^39,82,83^) during all behavioral sessions. A camera (Imaging source DMK 21Bu04.H) with zoom lens (Navitar 7000) coupled with an infrared filter (Mightex, 092/52×0.75) was placed ∼22 cm from the animal’s right eye. The eye was illuminated by a near-infrared LED (Mightex, SLS-02008-A). Video files were acquired using the Image Acquisition Toolbox in MATLAB with custom code. ∼74 pixels in each frame of the video was equal to 1 mm. We focused analysis on a high-quality subset of these sessions (Fig. S3; 3 PV-ChR2 mice, 27 sessions; 4 SST-ChR2 mice, 28 sessions).

Pupil position and area were acquired and analyzed as previously described^34,35,76^. Δ Pupil area was calculated as the percent deviation from the mean 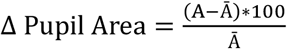, where A is the area in pixels and 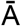 is the average area across all frames in a video. Δ Pupil position (in azimuth) was calculated as Δ Pupil position =x − x_Avg_, where x is position in degrees and x_Avg_ is average pupil position across all frames in a video. The pupil position in degrees was calculated assuming the eye was a sphere using the equation 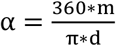, where m is the position in mm, d is the diameter of the eye (∼3.4 mm) based on a previous study^84^. Laser trials were subtracted from control trials to determine if pupil position or pupil area was different between the two conditions. Importantly, there were no differences in pupil metrics for PV versus SST mice (Fig. S3).

### Recordings: Visual detection behavior

A small craniotomy (0.1 – 0.3mm) was performed in either binocular (∼0.5 mm anterior to lambda, ∼2.5 – 3 mm lateral to central) or monocular V1 (∼0.5 mm anterior to lambda, ∼2 – 2.5 mm lateral to central). Craniotomy sites were confirmed with ISI and receptive field mapping. Mice recovered for at least 3 hours before recording. After recordings the recording chamber was covered with an elastic polymer (KwikCast) to preserve the skull and craniotomy health between consecutive recording days (2-5 days). Recordings were performed using multi-site silicon electrodes (NeuroNexus A1×32) consisting of a single shank with 32 linear channels. Electrodes were inserted ∼1mm below the cortical surface. During visual detection behavior, targeted recordings were performed in binocular V1 (0 – 20°) while optogenetically stimulating PV or SST neurons in monocular V1 (∼70°).

### Recordings: Receptive field mapping

After every recording the receptive field of the recording site was mapped by presenting vertical flashing bars one at a time (9° in width, duration 0.1 s, inter-stimulus interval 0.3 s, 5-100% contrast; Randomized position) that tiled the entire visual display. These mapping sessions were used to confirm retinotopy identified by ISI (Fig. S1).

### Recordings: Whole-cell patch-clamp

We performed whole-cell current clamp (n = 30 neurons) or voltage clamp (n = 24 neurons) recordings from regular spiking (RS) putative excitatory neurons in awake mice as detailed in past studies ^57,85^. RS identity was determined by spike width and spike frequency adaptation to current pulses. We simultaneously optogenetically stimulated PV or SST neurons, either at a site ∼0.8mm distant from the recorded neuron (exactly as described above, see “Retinotopically targeted laser stimulation”), or directly at the recording site. The differing effects of distal PV versus SST stimulation were not explained by differences in the quality of electrical access. Series resistance was comparable across groups, both in current clamp (PV: 33.9 ±2.9 MΩ, n = 17; SST: 33.1 ± 2.9 MΩ, n = 13, *p* = 0.71) and voltage clamp recordings (after partial compensation, PV: 19.1 ±1.9 MΩ, SST: 25.6 ±2.2 MΩ, mean ± SEM, *p* = 0.1).

### Focal viral experiments

In some experiments, we used a viral strategy to express ChR2 and EYFP in the presence of Cre in PV or SST neurons. Using PV-Cre or SST-Cre mice, we injected virus (AAV5-EF1a-dblFlox-hChR2(H134R)-EYFP, addgene; 50 nL, 2.2e12 GC/mL) into monocular V1 at 0.4 and 0.8 mm beneath the cortical surface. In electrophysiological experiments (Fig. S20, S23), mice were implanted with a cranial window to use ISI to target monocular V1 (Fig. S15). Otherwise, monocular V1 was targeted using anatomical measurements based on ISI maps across multiple mice (∼0.5 mm anterior to lambda, 2.5 mm lateral to central suture). We allowed for ∼3 weeks of expression before experimentation.

For anatomical experiments, after the expression period, we perfused the mouse to extract the brain and sectioned using a vibratome (coronal sections, 0.1 mm width). We then used a confocal microscope to image the sectioned slices (at 20x) using a z-stack tile scan. Using a rotated image aligned to the pia and white matter tract, we took the maximum fluorescence intensity across layers, and measured the pixels with fluorescence greater than the background across cortical distance (medial to lateral), normalized to the injection site. This measurement allowed us to assess the projection density of PV and SST neurons.

In some mice with injections, we performed electrophysiological experiments similar to the ones performed in PV-ChR2 and SST-ChR2 transgenic mice. These experiments allowed us to better measure how SST and PV neurons provide lateral inhibition without potential artefacts from off-target laser stimulation. We first performed recordings in monocular V1 (the viral injection site), while activating V1 (1.7 and 6.5 mW) at various distances away from the recording site. We then measured how PV and SST neurons responded to the laser stimulation across various distances away from the recording site. In other experiments, we recorded in binocular V1 while activating monocular V1 (the injection site) to determine how distal activation of focally-expressing PV or SST neurons affects the neural activity and contrast tuning of RS neurons. For this analysis we only analyzed RS neurons with control responses fit well to a Naka-Rushton curve, and RS neurons that showed overall activity reductions with distal stimulation.

### PV and SST neuron identification and laser stimulation (optotagging)

Extracellular spikes were sorted using KlustaViewa Suite or Kilosort^86,87^. To identify PV or SST neurons, we measured the short latency responses to a strong brief laser pulse (0.04s, 6.5mW) positioned over the recording site and used the stimulus-associated spike latency test (“SALT”)^56^. Neurons that increased their activity rapidly (<10 ms) during the pulse were identified as tagged PV or SST neurons in PV-ChR2 or SST-ChR2 mice respectively. All neurons were additionally identified by waveform width and classified as fast-spiking (FS) putative PV neurons (narrow width) or regular-spiking (RS) putative excitatory neurons (broad width), per prior studies^23,34,35,76^. Optotagged PV neurons had narrow waveforms and were combined with non-tagged FS neurons for analysis (Fig. S5).

### Laminar Identification

In some analysis (Fig. S18) we looked at L2/3 neurons during distal stimulation. L2/3 was identified using the current source density (CSD) of local field potential (LFP) as in our prior studies^34,35,76^. The channel with the earliest visually-evoked CSD sink was identified, with neurons that fell within ±100 microns of that channel identified as being in L4. Neurons above the L4 channels were identified as L2/3. We identified RS neurons in L2/3 and measured how PV or SST stimulation changed the spiking activity (modulation index; Fig. S18).

### Perceptual contrast sensitivity analysis

The sensitivity index (d’) was measured across contrasts for control (no laser) and stimulation (PV or SST) conditions. A Boltzmann’s sigmoidal equation was fit to the data points to generate psychometric contrast response curves:

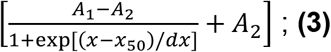

where A_1_ is the minimum d’, A_2_ is the maximum d’, x is the contrast, x_50_ is the contrast to reach 50% of the max d’, and dx is the steepness of the curve. The data was fit using the MATLAB function fit. From the Boltzmann’s sigmoidal fit, we quantified the maximum d’ (A_2_) and slope at the steepest point. A modulation index (MI) for the slopes was defined as 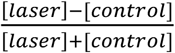 and used to calculate how PV or SST stimulation modulated the slope of the contrast sensitivity function relative to no laser (control). We also calculated the MI of the other parameters in the fits (e.g., A1 – minimum d’, A2 – maximum d’, x50 – contrast at 50% of max, or C_50_; Fig. S2).

### Neural contrast sensitivity analysis

To quantify neural contrast responses during perceptual behavior, the firing rate of RS neurons was calculated in the first 0.2s following stimulus onset for each contrast. We followed a similar strategy as behavioral data to estimate contrast response functions for firing rates, then assessed changes in the fit parameters during PV or SST lateral inhibition. Contrast response curves were fitted with Naka-Rushton equations ^88^:

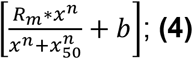

where R_m_ is the max firing rate, x is the contrast, n is the steepness, x_50_ is the contrast at 50% of the max firing, and b is the minimum firing rate. Only RS neurons that had clear contrast tuning and were well fit to the Naka-Rushton equation (r^2^ > 0.25) were included in analysis (146 RS neurons in SST-ChR2 mice, 166 RS neurons in PV-ChR2 mice). A modulation index (MI) was defined the same way as perceptual analysis 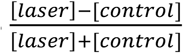 and used to calculate the relative changes in slope (slope at steepest point of the curve fits) elicited by PV or SST stimulation. We performed additional controls (Fig. 3B, C) to measure changes only in high firing RS neurons (>3 spikes/s), and also used a hierarchical bootstrapping method (100 samples) to confirm effects were not solely due to particular mice or sessions^41^.

We also used a hierarchical bootstrapping method to confirm effects were not solely due to particular mice or sessions^41^. It is typical to average stimulus evoked responses across neurons. However, this data is not strictly independent as multiple neurons are obtained within the same recording session, and sessions within the same mice. Averaging responses within sessions is also challenging as this greatly reduces the sample size. One approach to account for this issue is to use hierarchical bootstrapping, a statistical method to better quantify the uncertainty of observations in hierarchical datasets^41^. Using this approach we randomly resampled with replacement at each level. For each resample, we randomly selected: 5 recording sessions (based on average number of recordings per craniotomy, from a total of 26 sessions for PV and 18 for SST experiments); sampled the total number of neurons in that session (on average 15 neurons per session), 100 trials from each sampled session (on average 170 trials per session), and performed this resampling procedure 100 times for both the PV and SST stimulation groups. From each resampled population, we fit a Naka-Rushton equation to the control and stimulated (PV or SST) contrast responses for each neuron, and calculated the average slope MI across neurons well-fit by the Naka-Rushton equation (r2 > 0.25). This process provides estimates of the uncertainty around the population average. Should <95% of the bootstrapped average MI estimates fall below 0, that would indicate a significant decrease in slope.

In addition to measuring the slope, we calculated the MI of the other parameters in the fits (e.g., Rm – max firing rate, b – minimum firing rate, x50 – contrast at 50% of max response; Fig. S6). We also performed control analysis constraining the steepness parameter ranges (n, from range [0-30] to [0-5]) to determine any potential artefact from overly steep fits. We found restricting the parameter range did not affect the results.

### Affine model quantifying subtractive and divisive modulation

We tested whether a subtractive, divisive, or affine model (subtractive and divisive) would best fit the psychometric and neural contrast response curves during PV or SST stimulation. To implement the subtractive model, we fit the psychometric contrast responses during PV or SST stimulation to the Boltzmann’s sigmoidal equation (eq. 3), where we only allowed A_2_ and x_50_ to vary (taking the other parameters from the control fits). A similar approach was taken to fit a subtractive model to the neural data, except using the Naka-Rushton Equation, where we only allowed the b and x_50_ parameters to vary. For the divisive model, only A1 was varied for the psychometric responses, with the other parameters fixed (taken from the control fits); R_m_, n, and x_50_ was varied for the neural responses in the divisive model. Only sessions where PV or SST stimulation decreased the response were used for this analysis (decrease in d’ or firing rate). For PV and SST stimulation, both the perceptual and neural responses fit the affine model substantially better than purely subtractive or divisive models, indicating that PV and SST stimulation have a mixture of subtractive and divisive effects; this is somewhat expected since the affine model enables more parameters to fit the data.

In Fig. S4, we compared the subtractive and divisive effects in the perceptual and neural response by using an affine fit to the data and comparing the subtractive (i.e., changes in vertical offset; A_2_ for psychometric data, b for neural data) and divisive (changes in gain, maximum slope changes) components of the fits. All changes were quantified using the modulation index.

### Neural and perceptual correlation analysis

We assessed the strength of correlation between neural and perceptual changes to contrast sensitivity on the same trials. We calculated the changes in slope of the contrast sensitivity functions (MI, as described above), then calculated Spearman’s rank correlation coefficient between the neural MI and perceptual MI acquired during the same sessions (Fig. 3F; neural activity binned between -1 and 1, bin size = 0.5). Similar results were obtained without binning (distal PV stimulation, ρ = 0.125, *p* = 0.109; distal SST stimulation, ρ = 0.315, *p* < 1e-3).

In addition, to analyze the perceptual and neural contrast sensitivity, we also looked at the relationship between d’ and spiking activity, agnostic to contrast. We calculated a modulation index during PV and SST stimulation relative to the control condition, for both the perceptual (d’) and RS spiking activity. We used a Spearman’s correlation to measure the relationship between changes in neural and perceptual activity; there was a significant relationship between overall neural activity and overall performance for both PV and SST stimulation (Results).

### Neural decoder analysis

We performed neural decoding of perceptual responses using a linear classifier (Support Vector Machine, linear kernel, fitcsvm in MATLAB). The SVM reports binary classification of perceptual outcomes solely from the neural data. We identified a hyperplane (*y*) that best separates the different classes of behavioral responses (correct or incorrect trials), using the RS activity on a given trial in each session. The hyperplane is constructed as : *y = x′β+b*, where *x* is the firing rate, *β* is a vector of coefficients orthogonal to *y*, and *b* is the bias term. Given the importance of capturing the trial-by-trial relationships among simultaneously recorded neurons and behavioral performance, and the heterogeneous sampling of neurons at the individual session level, the classifier was trained for each recording session on either the PV (26 sessions) or SST (14 sessions) stimulated groups. All RS neurons within session were used regardless of contrast tuning (PV: 338 total RS neurons, SST: 319 total RS neurons). The SVM was trained on the control trials only from each session (PV: 2592 total trials, SST: 1652 total trials), with the laser stimulation trials held out (PV: 963 total trials, SST: 773 total trials). We then used the control or laser trials to predict a binary behavioral response (correct trial indicated by lick during response window, or not) for a given trial across each recording session. Importantly, there was no contrast information in the model. We assessed the classification accuracy by measuring the probability that the SVM correctly identified the behavioral responses from the RS activity during control or laser stimulation trials across different contrasts. The accuracy was above chance and contrast dependence of behavior emerged just from firing rates (Fig. S8). We then estimated the hit rates and false alarm rate (0% contrast trials) using the classifier predictions within each session. These estimated hit and false alarm rates were used to calculate the d’, which allowed us to reconstruct the psychometric contrast response curves. We fit these predicted psychometric contrast responses curves to a Boltzmann sigmoidal equation and we measured the slope at the inflection point. We then calculated the change in slope (MI) during PV and SST stim. In addition, we also constructed the classifier excluding RS neurons with strong sensitivity changes (MI = -1; PV: n = 94; SST: n = 98), and measured how excluding these neurons affected the slope of the psychometric curves. Importantly, constructing a single SVM trained on half the control trials from *all* sessions, then tested on held-out control & stimulation trials, showed similar results (Fig. S8C).

### Local and distal stimulation neural activity analysis

We measured the RS, FS, and SST responses to a brief laser pulse (0.04s duration, 0.5 – 6.5 mW) during local stimulation (recording and manipulation sites are the same) and distal stimulation (recording and manipulations sites are ∼0.8 mm apart) in V1. We calculated a modulation index of firing rates to measure the magnitude of change:

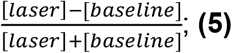

(baseline taken from the pre-stimulus laser period).

### Morphology Reconstructions

To analyze the morphology of PV and SST neurons, we analyzed publicly available data of single neuron patch clamp recordings with morphological fills from the Allen Brain Atlas Cell Database^51^. We reconstructed the morphology of these neurons, rotating the axes to align to the pia and white matter. The axonal and dendritic lateral extent, as well as overall lateral projection density (fluorescence levels across space, normalized to the injection site) were measured. We also reconstructed a set of publicly available SST Martinotti cells^48^ to verify long-range axonal projections (downloaded from data posted on Zenodo).

### Statistical analysis

In general, non-parametric Wilcoxon rank sum tests (unpaired data) or signed rank tests (paired data) were performed, unless otherwise noted. Significance was defined at α = 0.05, unless noted. No strategies were employed for randomization of subjects, recordings, data collection, or analysis, aside from the hierarchical bootstrap. Statistical details are described in Table S1.

### Circuit model: Network construction

A leaky-integrate-and-fire (LIF) spiking neural network model of the V1 circuit was used to test the effects of long-range SST projections and nonlinear dendritic integration on cell type-specific lateral inhibition. The basic construction of the network was based on prior models of cortical excitatory and SST and PV inhibitory cell types^44,89^. It consisted of *N*_Exc_ = 8,000 excitatory, *N*_PV_ = 1,000 PV, and *N*_SST_ = 1,000 SST neurons uniformly distributed on a 1*×*1 mm^2^ two-dimensional surface with periodic boundary conditions.

All neurons were modelled as conductance-based leaky integrate-and-fire (LIF) neurons. The membrane potential of the *j*th neuron *V*_*j*_ is described by

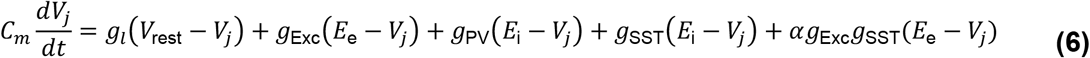

where the last term *αg*_Exc_*g*_SST_(*E*_e_ − *V*_*j*_) is only included for excitatory neurons. A neuron fires a spike when it reaches threshold *V*_th_ = −40 mV. During a refractory period of τ_ref_ = 5 ms, its voltage is then held at *V*_reset_ = −65 mV before resuming evolution described by Eqs. 6 & 7. Here *C*_*m*_ is the membrane capacitance, *g*_*l*_ is the leak conductance, *g*_Exc_, *g*_PV_, and *g*_SST_ are the synaptic conductances from excitatory, PV, and SST neurons, respectively, *V*_rest_ is the resting membrane potential, *E*_e_ and *E*_i_ are the reversal potentials of the excitatory and inhibitory synapses, and *α* is a constant determining the strength of dendritic integration effects, as in prior studies^45^. The first four terms on the right-hand side of Eq. 6 represent the leak current and the synaptic currents from excitatory, PV, and SST inputs, while the last term involving the product of *g*_Exc_ and *g*_SST_ describes the interaction of excitatory and SST synaptic inputs within the dendrite (see below).

The synaptic conductances evolve according to

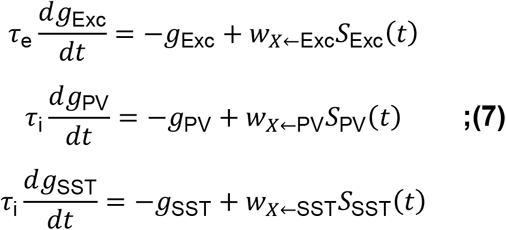

where τ_e_ and τ_i_ are excitatory and inhibitory synaptic time constants and *W*_*X*←Exc_, *W*_*X*←PV_, and *W*_*X*←SST_ are synaptic weights of synapses to population *X* (where *X* is the cell type of neuron *j*) from excitatory, PV, and SST neurons, respectively. *S*_Exc_, *S*_PV_, and *S*_SST_ are sums of Dirac delta functions representing incoming spike trains from excitatory, PV, and SST neurons, respectively. Neuron model parameters are based on the model by El-Boustani and Sur^44^ and are in the physiological range^90,91^.

Connection probabilities between neurons of each cell type were derived from published neuroanatomical data^20,21^. The density of connections from type *Y* to type *X* is given by *p*_*X*←*Y*_ whose values are listed in Table 1. We also incorporated long-range projections which are controlled by the parameters 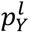. Specifically, each directed pair of neurons from type *Y* to type *X* was connected with a probability *P*_*X*←*Y*_ which depends on their distance *d*,

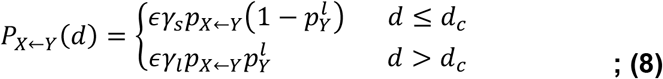

Here *d*_*c*_ is the cutoff distance given by 0.2 mm, *ϵ* controls the overall connectivity density of the network, and *γ*_*s*_ and *γ*_*l*_ are normalization parameters given by 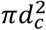 and 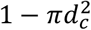, respectively. There are a total of *ϵp*_*X*←*Y*_*N*_*X*_ projections from a neuron of type *Y* to neurons of type *X*, where *N*_*X*_ is the total number of neurons in the network of type *X*. Among the *ϵp*_*X*←*Y*_*N*_*X*_ projections, 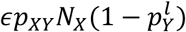 are short-range and 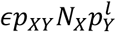 are long-range.

**Table 1:** Table of connection densities. The parameter *p*_*X*←*Y*_ determines the relative density of connections from neurons of type *Y* to neurons of type *X*. Values were based on prior study^**20**^.

| | | $p_{X \leftarrow Y}$ | | |
| --- | --- | --- | --- | --- |
| | | $X$ | | |
|  |  | Exc | PV | SST |
| $Y$ | Exc | 0.06 | 0.42 | 0.3 |
|  | PV | 0.35 | 0.39 | 0 |
|  | SST | 0.23 | 0.18 | 0 |

### Circuit model: Long-range projections

The parameters 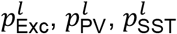 control the proportion of long-range excitatory, PV, and SST projections. Experimental evidence suggests that SST long-range projections are more prominent compared to those of PV neurons ^42^. Excitatory and SST neurons were given a high probability of long-range projections 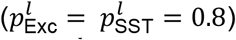, while PV neurons were assumed to have a smaller long-range probability 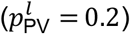. 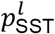 is varied between 0 and 0.8 to examine the effect of long-range SST projections. Varying 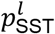 does not change the total number of synapses but only the proportion of those that are long-range (Eq. 8).

### Circuit model: Dendritic integration

PV neurons form perisomatic synapses on excitatory pyramidal cells, while SST neuron synapses target pyramidal cell dendrites^47,71,92,93^. The point-neuron LIF model implements SST dendritic inhibition of excitatory neurons by incorporating interactions of excitatory and SST inhibitory synaptic inputs that occur in the dendrites^94^. This approach has been validated against experimental data and a multicompartment biophysical model^94,95^. Eq. (6) mimics dendritic integration of excitatory synaptic inputs and SST inhibitory inputs via the multiplicative term *αg*_Exc_*g*_SST_(*E*_e_ − *V*_*j*_), while PV inputs sum linearly with excitatory inputs (mimicking linear somatic integration of excitation and inhibition). The parameter *α* determines the strength of the effect of dendritic inhibition on excitation. Generally, this parameter depends on the distances of the excitatory (E) and inhibitory (I) synapses from one another, and from the soma^45,46^. For example, E and I synapses on opposite ends of the dendrites have low interaction strength (*α* values –8 kΩ cm^2^), but E and I synapses proximal to each other but distant from the soma have much higher interaction strength (*α* values 20 kΩ cm^2^; all values from Fig. 4 in prior study^95^). Using the estimated surface area of the neuron, we converted this range into *α* values of –80 to 200 MΩ for our point models. For simplicity, it was assumed that all excitatory and SST synapses interact with a single constant *α* representing the average dendritic effect propagated to the soma and spike initiation site. We begin by assuming *α* = 70 MΩ (conservatively within the range of Li et al^45^) and 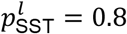 and tune a subset of parameters so that the spontaneous firing rates and evoked response without ChR2 stimulation approximately matched the data (see Circuit Model: Parameter Fitting). The parameter *α* was then varied in the range of 40-80 MΩ to explore the impact of dendritic inhibition on scaling the contrast dependence of firing rate. This range was chosen as it represents the values at which the spontaneous and evoked firing rates are within a realistic range (given the fixed parameter values in Tables 1 and 2). Because positive values of *α* result in stronger inhibition to excitatory neurons, the total inhibition from SST to excitatory neurons depends on a combination of *W*_Exc←SST_ and *α*. Decreasing *α* to values below 40 MΩ excessively weakens the strength of SST inhibition to excitatory neurons, leading to unrealistically high responses. For example, when *α* = 0 MΩ, firing rates for high contrast grating responses were 10-fold higher (∼75 spikes/s) than experimental data (∼7 spikes/s). However, one of our main results (Fig. 5) is that the dendritic integration term is not necessary to replicate the divisive effects of SST inhibition; even when *α* = 0 MΩ the effect is restored by increasing *W*_Exc←SST_ (not shown here). Thus, choosing *α* = 40 MΩ evoked 7 -10 spikes/s for high contrast responses, matching the control experimental results. Conversely, increasing dendritic effects beyond our upper bound of *α* = 80 MΩ suppressed excitatory neurons to an unrealistic degree: responses during the simulated ChR2 activation of SST neurons dropped to 0.1-0.3 spikes/s, 10-fold lower than the experimental range (2 – 4 spikes/s), and increasing *α* further suppresses the rates.

**Table 2:** Table of physiological parameters of the leaky integrate- and-fire (LIF) neuron models.

| Parameter | Description | Value | Units |
| --- | --- | --- | --- |
| $C_m$ | membrane capacitance | 0.25 | nF |
| $V_{\text{rest}}$ | leak potential | -70 | mV |
| $V_{\text{thr}}$ | threshold potential | -40 | mV |
| $V_{\text{reset}}$ | reset potential | -65 | mV |
| $\tau_{\text{ref}}$ | refractory period | 5 | ms |
| $E_e$ | reversal potential for excitatory synapses | 0 | mV |
| $E_i$ | reversal potential for inhibitory synapses | -75 | mV |
| $\tau_e$ | excitatory synaptic time constant | 6 | ms |
| $\tau_i$ | inhibitory synaptic time constant | 20 | ms |

### Circuit model: Simulating visual input and ChR2 stimulation

In all simulation conditions, the V1 network receives spontaneous inputs from 1,000 Poisson-spiking neurons with firing rate *r*^*b*^. Each neuron in the V1 circuit receives 1,000 *× ϵ*_ext_ incoming excitatory connections from the Poisson neurons with synaptic weight 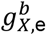 for neurons of type *X*. SST neurons receive an additional 1,000 *× ϵ*_ext_ inhibitory connections with synaptic weight 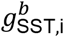.This is because SST neurons receive no other inhibitory input in this simple 3 neuron network, but experimental evidence suggests that they receive inhibitory inputs from multiple sources^20^.

Visual stimulation is implemented by an external thalamic layer (T) consisting of 1,000 neurons with inhomogeneous Poisson firing statistics, also uniformly distributed on a 1 mm *×* 1 mm surface. Each neuron in the thalamic layer projects to excitatory and PV+ neurons within a lateral distance of 0.2 mm with density *ϵ*_ext_ with synaptic weights *W*_Exc←*T*_ and *W*_PV←*T*_, respectively. At visual stimulus onset, the firing rates of the external thalamic layer neurons are activated with a spatiotemporal Gaussian pattern with spatial *σ* = 0.2 mm and temporal *σ* = 30 ms. The firing rate peaks 0.1s after stimulus onset with a peak average firing rate of *r*^*ext*^ which depends on the contrast *c* of the visual stimulus by the relation:

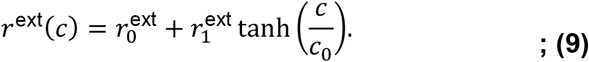

We model ChR2 activation of inhibitory cell types by stochastically injecting a conductance 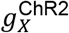 with a time-dependent rate to all neurons of the target cell type *X* inside a circular area with radius 0.3mm (matching experiments, Fig. 2; Fig. 4). The rate of activation is linearly ramped up starting from 0.1s before the visual stimulus onset at a rate such that it reaches 1000 Hz at the visual stimulus onset time, again matching experiments. The rate is then held constant for 0.3s, after which the ChR2 activation is turned off and the rate set to zero.

In most simulations, the center of the laser stimulus was located at a maximal distance ∼0.71 mm from the peak of the visual stimulus (except in Fig. S7 that measured the distance dependence of the effects). For simulations presented in Figs. S12 and S22, which examined the effects of laser distance, the position of the laser was varied over a straight line between the distal location (0.71 mm distance) and the center of the visual stimulus (0 mm). When targeting SST neurons, we assume that a randomly selected proportion *p*_*ChR*2_ of SST cells outside the targeted area also receive stimulation due to activation of long-range axonal projections. Including off-target SST activation was necessary for the response of local SST neurons to match those from recorded data. However, the divisive effects of distal SST stimulation did not depend on antidromic SST stimulation in the model (see Fig. S24). Parameters related to external inputs are given in Table 3.

**Table 3:** Table of parameter values used in network simulations.

| Parameter | Description | Value | Units |
| --- | --- | --- | --- |
| $r^b$ | Spontaneous input rate | 2 | Hz |
| $g_{\text{Exc},e}^b$ | Synaptic weight of spontaneous excitatory inputs to Exc | 13.5 | nS |
| $g_{\text{PV},e}^b$ | Synaptic weight of spontaneous excitatory inputs to Exc | 12.0 | nS |
| $g_{\text{SST},e}^b$ | Synaptic weight of spontaneous excitatory inputs to SST | 12.0 | nS |
| $g_{\text{SST},i}^b$ | Synaptic weight of spontaneous inhibitory inputs to SST | 3.0 | nS |
| $w_{\text{Exc} \leftarrow T}$ | Synaptic weight from thalamic layer to Exc neurons | 2.0 | nS |
| $w_{\text{PV} \leftarrow T}$ | Synaptic weight from thalamic layer to PV neurons | 1.0 | nS |
| $r_0^{\text{ext}}$ | Visual input parameter | 4 | Hz |
| $r_1^{\text{ext}}$ | Visual input parameter | 30 | Hz |
| $c_0$ | Contrast nonlinearity parameter | 16 | % |
| $g_{\text{PV}}^{\text{ChR2}}$ | ChR2 quantal conductance for PV | 15 | nS |
| $g_{\text{SST}}^{\text{ChR2}}$ | ChR2 quantal conductance for SST | 1.3 | nS |
| $p_{\text{off-ChR2}}$ | Proportion of “off-target” SST antidromic stimulation | 5 | % |
| $w_{\text{Exc} \leftarrow \text{Exc}}$ | Synaptic weight from Exc to Exc | 1.2 | nS |
| $w_{\text{PV} \leftarrow \text{Exc}}$ | Synaptic weight from Exc to PV | 0.1 | nS |
| $w_{\text{SST} \leftarrow \text{Exc}}$ | Synaptic weight from Exc to SST | 0.1 | nS |
| $w_{\text{Exc} \leftarrow \text{PV}}$ | Synaptic weight from PV to Exc | 1.5 | nS |
| $w_{\text{PV} \leftarrow \text{PV}}$ | Synaptic weight from PV to PV | 1.5 | nS |
| $w_{\text{Exc} \leftarrow \text{SST}}$ | Synaptic weight from SST to Exc | 0.8 | nS |
| $w_{\text{PV} \leftarrow \text{SST}}$ | Synaptic weight from SST to PV | 0.8 | nS |
| $\alpha$ | Dendritic interaction strength between E & SST input in E neurons | 70 | MΩ |
| $\epsilon$ | Connectivity density parameter | 0.2 | - |
| $\epsilon_{\text{ext}}$ | External connectivity density parameter | 0.05 | - |
| $p_{\text{Exc}}^l$ | Proportion of long-range Exc projections | 0.8 | - |
| $p_{\text{PV}}^l$ | Proportion of long-range PV projections | 0.2 | - |
| $p_{\text{SST}}^l$ | Proportion of long-range SST projections | 0.7 | - |

### Circuit model: Parameter fitting

We adapted the neuron model parameters from a prior study^44^ (Table 2). Parameters relating to spontaneous inputs (*r*^*b*^, 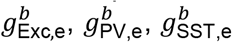 and 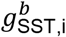; see Table 3), visual stimulus (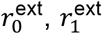, *W*_Exc←*T*_ and *W*_PV←*T*_), and synaptic weights (*W*_Y←X_ for all populations X and Y) were tuned to fit the model to the observed spontaneous firing rates and contrast curves measured in experiments. This was achieved with a combination of gradient descent minimizing the root mean square error of the average firing rates of each condition (spontaneous, contrasts 2-33%), followed by slight hand-tuning to capture both spontaneous and stimulus-driven regimes. During automated error minimization, the gradient of the error function was approximated using finite difference approximation. The fitting parameters were constrained to fit a physiologically plausible range, which in interval notation were [0.1 nS, 50 nS] for all synaptic weights and [1 Hz, 100 Hz] for spontaneous and thalamic layer firing rates. The ChR2 conductance parameters 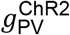 and 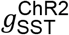 were chosen so that the mean firing rates of the targeted inhibitory neurons matched those from the observed data.

### Circuit model: Off-target dendritic stimulation

To rule out the possibility that a high dendritic excitability of SST neurons drove the main results, we performed additional simulations incorporating off-target dendritic activation of SST neurons. In this alternative model, random antidromic off-target activation was removed. Instead, SST neurons were assumed to have dendritic projections with a lateral radius of 0.3 mm. If the dendritic range overlapped with the laser stimulus, the neuron was stimulated with a reduced strength of 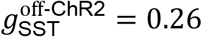 nS, whose value was determined by comparing the mean firing rate of SST neurons in the visual stimulus area with that from experimental data. As with the original model, we then varied 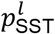 to determine if long-range lateral SST projections are necessary to reproduce the divisive effects (see Fig. S13).

### Circuit model: Properties of Cortical Networks

We tested whether the circuit model exhibited known properties of cortical networks. To test whether the network is inhibition-stabilized^49^, we removed inhibition by setting all synaptic weight values (i.e., *W*_Y←X_ where X is PV or SST) to zero. When removing inhibition locally, this change was confined to synapses from inhibitory neurons within a circle of radius 0.3 mm; when removing inhibition globally, this was done for all inhibitory synapses in the network. The resulting spontaneous and evoked firing rates were compared to those of the original model (see Fig. S10). We also examined whether the network exhibits asynchronous and irregular activity during spontaneous activity and under visual stimulus by computing the inter-spike interval (ISI) distributions of excitatory neurons during the specified stimulus conditions (Fig. S11). The coefficient of variation (CV) of an ISI distribution was defined as *σ*^2^/*µ*^2^, where *µ*^2^ is the mean and *σ*^2^ the variance of the distribution^50^.

### Circuit model: Simulation Details

Simulations were performed with PyNEST v. 3.3 ^96^ and NESTML v. 5.2.0 ^97^ with a time step of 0.1 ms. Tables 1 – 3 (following) describe all key model parameters

## Data and Code Availability

Data and code necessary to reproduce the main results will be made publicly available at DOI 10.6084/m9.figshare.24517960 upon publication.

All LIF simulation code is available at https://github.com/HChoiLab/CellTypeCircuit.

---

**Figure S1.**
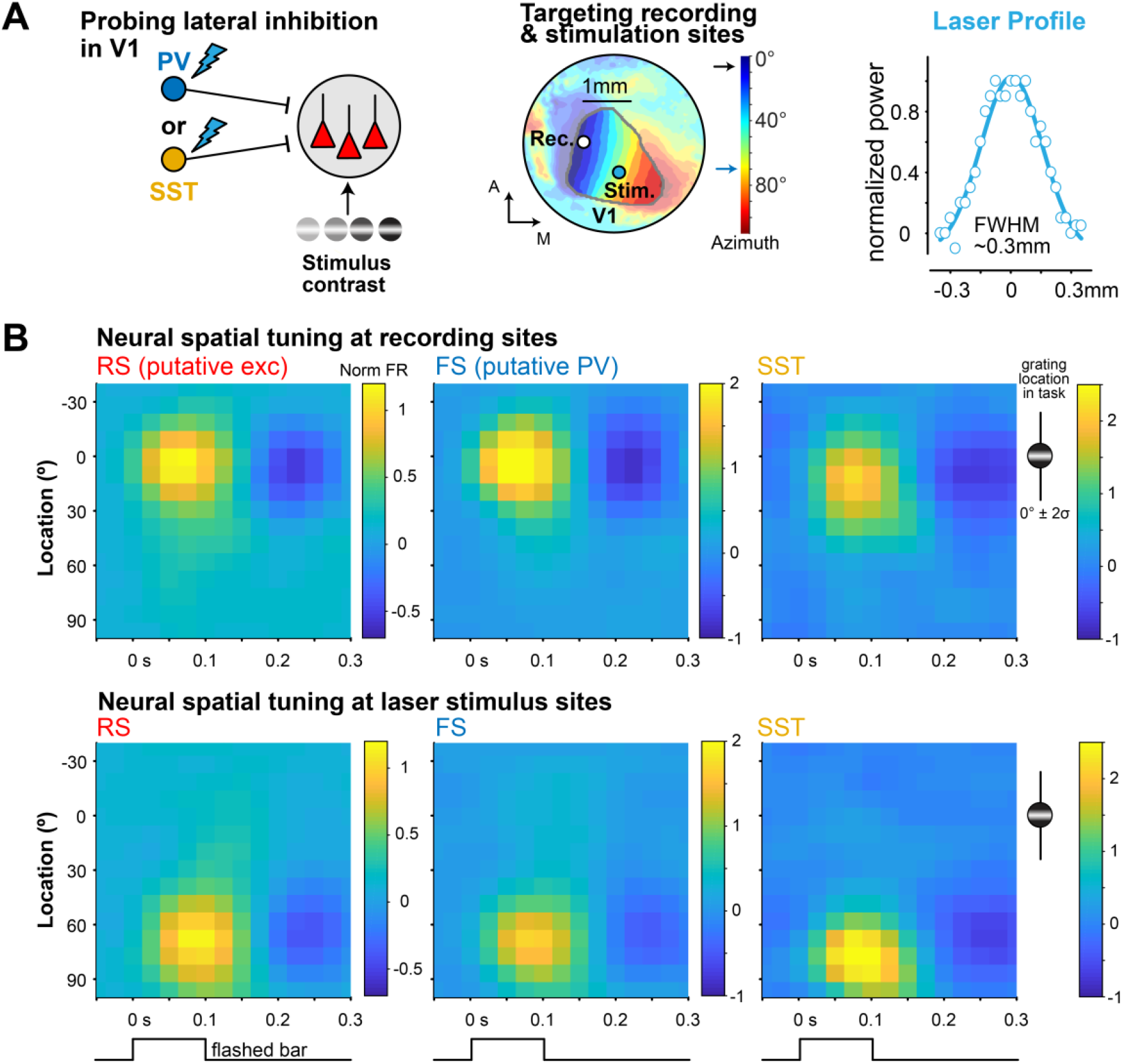
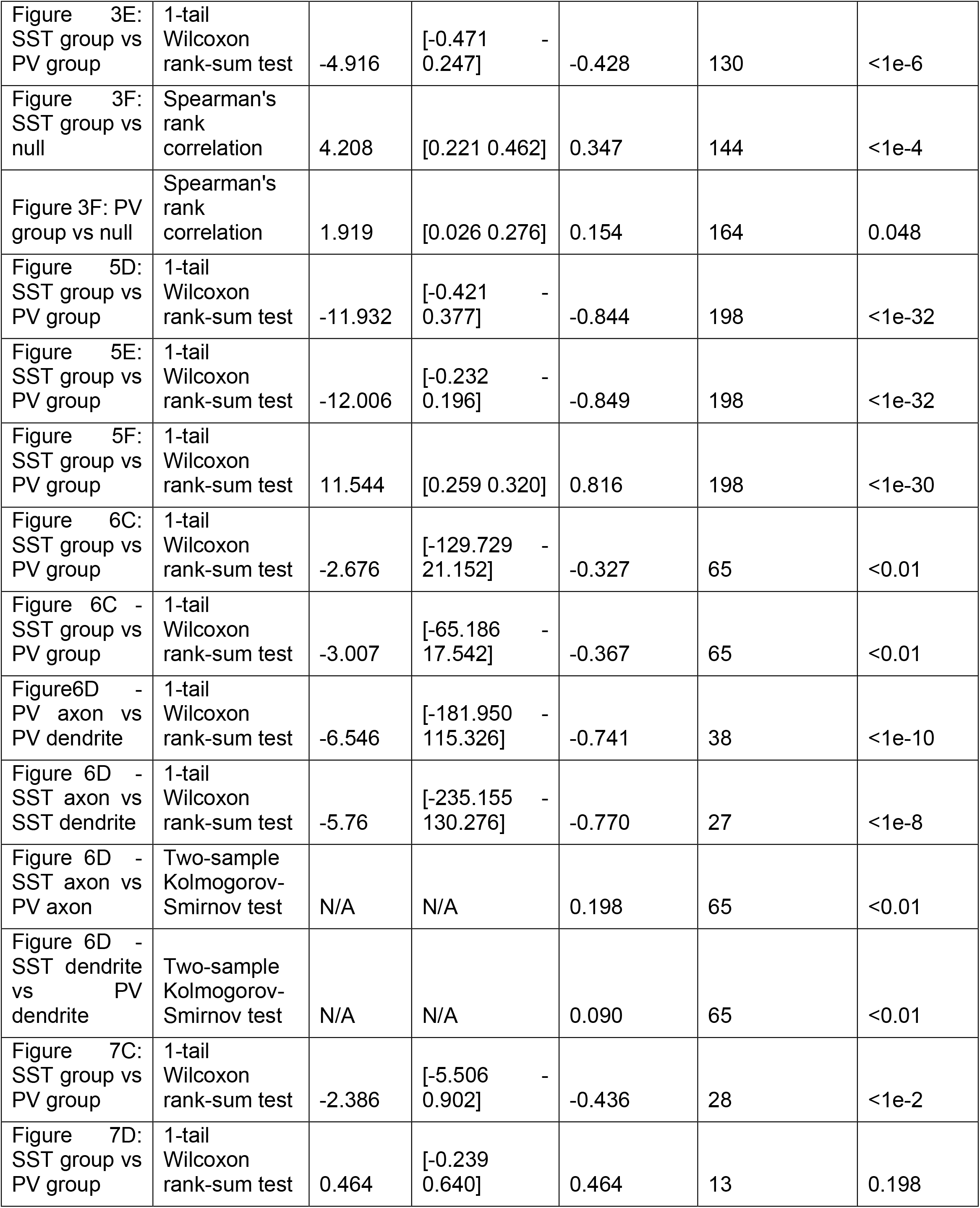
Precise targeting of retinotopic sites in V1. **A**. Left, illustration of experimental paradigm to record from a retinotopic site in V1 while distally activating PV or SST neurons at another retinotopic site. Middle, example ISI map showing targeted recording and stimulation sites corresponding to different visual retinotopic locations in V1 (same as Fig. 1). Right, the laser beam profile size was small (0.334 mm full-width half-max) allowing precise stimulation of specific retinotopic sites. **B**. The receptive fields of all neurons (1612 RS; 354 FS; 50 SST) were mapped by flashing bars (9° width, 0.1 s duration, 0.3 s inter-stimulus-interval) across multiple azimuth positions in the visual field (y-axis). Top, neurons in the targeted recording site in V1 had receptive fields at 10.2 ± 25.4° (peak ± σ), that largely overlapped with the task-relevant visual stimuli (Gabor Grating centered at 0°, σ = ∼10°). Bottom, neurons in the targeted laser stimulation site had receptive fields at 73.4 ± 25.6° and were not activated by and did not overlap with the task-relevant visual stimuli. Z-scored firing rate activity plotted.

**Figure S2.**
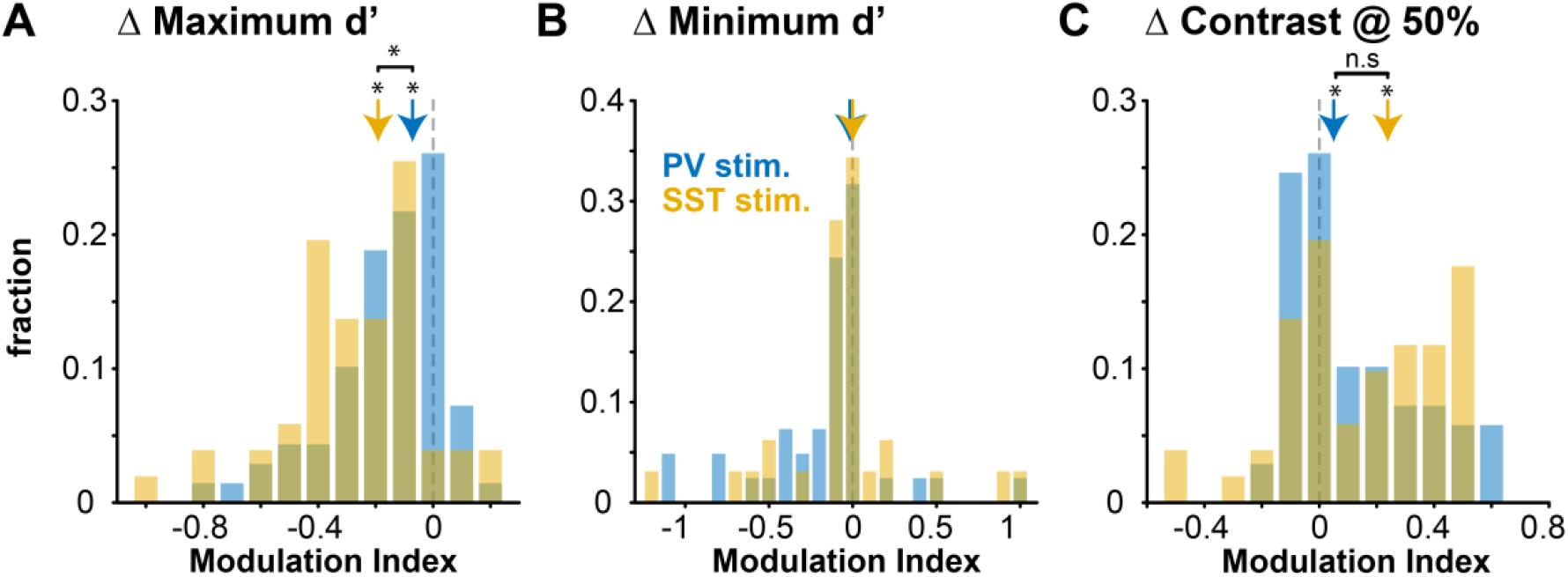
Changes in perceptual contrast response function (CRF) parameters. **A**. Both PV (−0.07 ± 0.15 MI, median ± mad, p < 1e-3, sign rank test) and SST stimulation (−0.19 ± 0.19 MI, p < 1 e-6) significantly decreased max d’, and this was greater for SST versus PV stimulation (p < 1e-2, rank sum test). **B**. No reduction in minimum d’ with PV (0.00 ± 0.41 MI, p = 0.064) or SST stimulation (0.00 ± 0.41 MI, p = 0.57), with no difference between PV or SST stimulation (p = 0.23). **C**. Both PV (0.05 ± 0.18 MI, p < 1e-6) and SST stimulation (0.24 ± 0.23 MI, p < 1e-4) significantly increase contrast needed for 50% performance (C_50_), with no difference between each other (p = 0.13). The increase in C_50_ in laser versus control trials (ΔC_50_, not normalized by sum as in MI) was significantly greater with SST vs PV stimulation (SST: 0.05 +/-0.06, PV: 0.01 +/-0.04; p =0.041).

**Figure S3.**
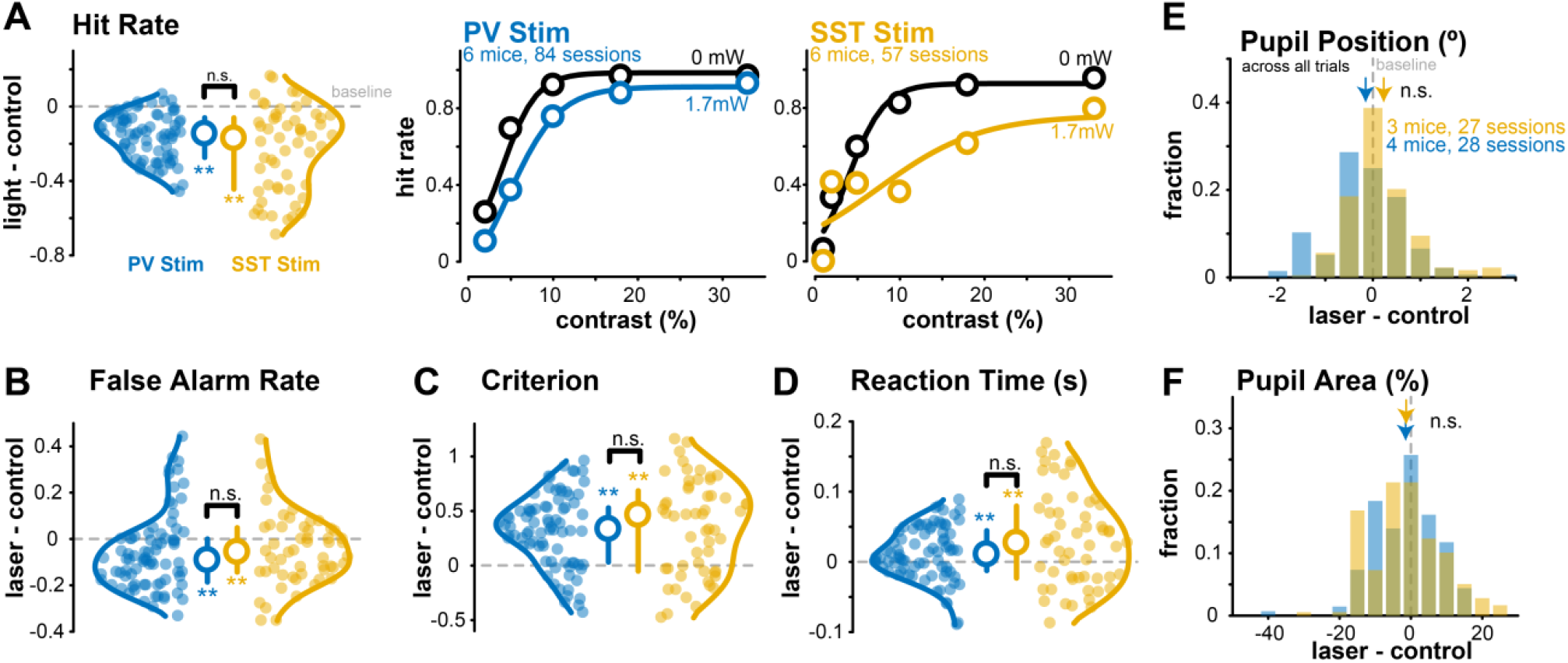
Behavioral metrics, pupil area, pupil position in PV and SST mice. **A**. Left, hit rate changes with distal PV stimulation (6 mice, 84 sessions) and distal SST stimulation (6 mice, 57 sessions), averaged over all contrasts (1 – 33%, Median ± IQR plotted). The hit rate decreases for both distal PV stimulation (Δ (laser – control) = –0.15 ± 0.11, median ± MAD, *p* < 1e–12, Wilcoxon signed-rank test) and distal SST stimulation (Δ (laser – control) = –0.17 ± 0.21, *p* < 1e–7). The overall magnitude of decrease in hit rate is not significantly different with distal PV vs SST stimulation (*p* = 0.07, Wilcoxon rank-sum test). Middle, hit rate sorted by contrasts in PV-ChR2 mice during control (0 mW) and distal PV stimulation sessions. Right, hit rate across contrasts for SST-ChR2 mice (6 mice, 57 sessions). Mean ± SEM, sigmoidal fits plotted. Distal SST stimulation decreases the hit rate slope (Δ (laser – control) = –0.06 ± 0.24, median ± MAD, *p* = 0.04), while PV stimulation slightly increases the slope (Δ = 0.03 ± 0.17, *p* = 0.02). **B**. The false alarm rate decreases with distal SST stimulation (Δ = –0.06 ± 0.12, *p* < 0.01) and distal PV stimulation (Δ = –0.09 ± 0.13, *p* < 1e–3) and is not significantly different across groups (*p* = 0.16). **C**. The criterion (likelihood to withhold from licking) increases with distal SST stimulation (Δ = 0.46 ± 0.37; *p* < 1e–5) and distal PV stimulation (Δ = 0.34 ± 0.26, *p* < 1e–9). The magnitude of increase is not significantly different (*p* = 0.30). **D**. The reaction time slows with distal SST stimulation (Δ = 0.03 ± 0.07, *p* < 0.01) and distal PV stimulation (Δ = 0.01 ± 0.03 s, *p* < 0.01) with no significant difference across groups (*p* = 0.11). **E**. The pupil position is similar between laser and control trials during distal PV stimulation (Δ (laser – control) = –0.17 ± 0.61°, *p* = 0.14) and SST stimulation (Δ = 0.15 ± 0.50°, *p* = 0.08). **F**. Pupil area (a proxy for arousal, measured as % change from mean) is not different in laser vs control trials during PV stimulation (Δ = –0.61 ± 7.10 %, *p* = 0.11) and SST stimulation (Δ = –1.30 ± 8.15 %, *p* = 0.06).

**Figure S4.**
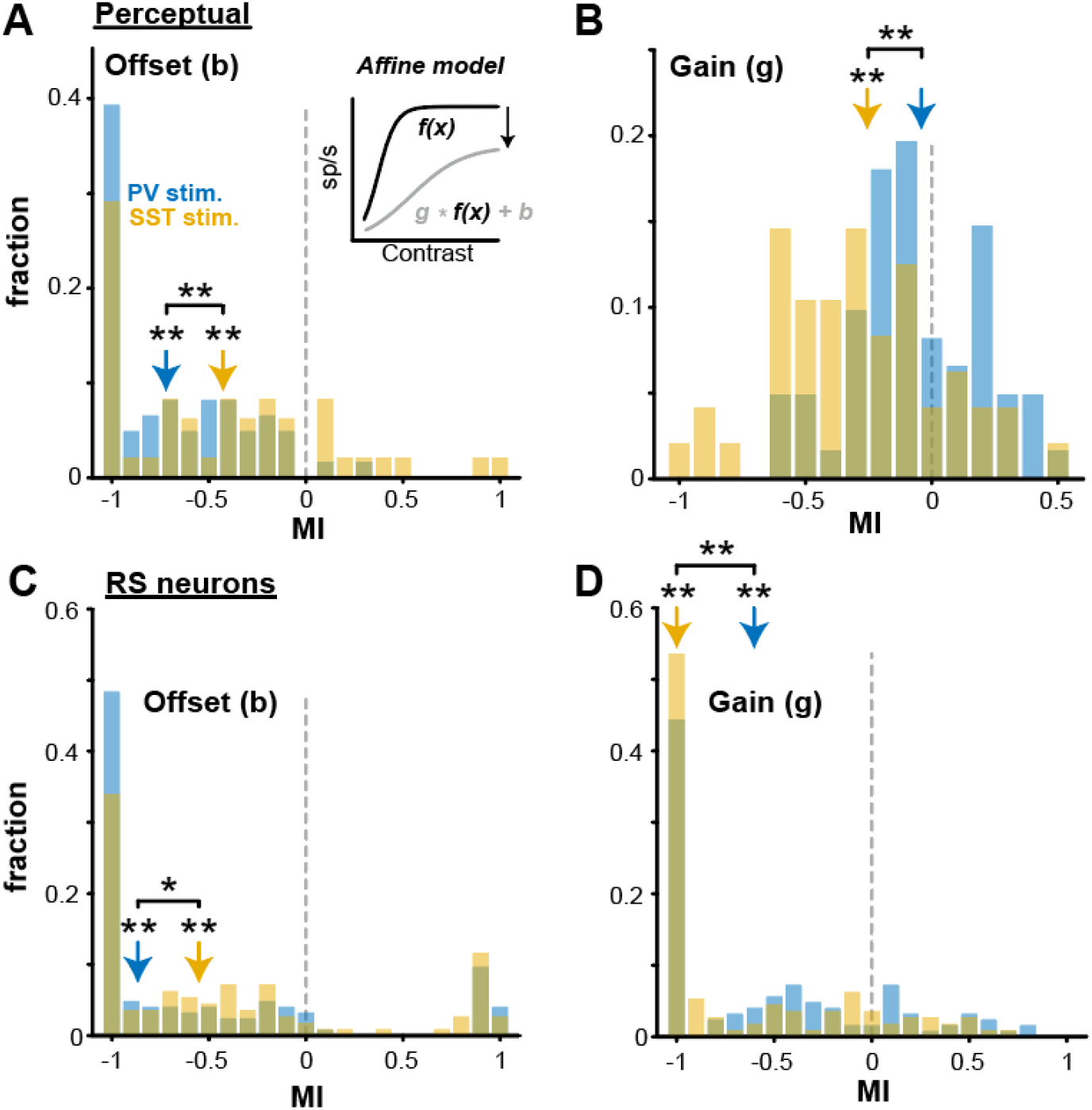
Affine model reveals divisive and subtractive effects. **A**. Affine model with gain (g) and offset (b) parameters acting on the control perceptual psychometric function (*f(x)*, inset). Both PV (–0.72 ± 0.28 MI, median ± MAD, 61 sessions, p < 1e-10, sign rank test) and SST stimulation (–0.43 ± 0.48, 48 sessions, p < 1e-4) decrease the offset (subtraction). Significantly larger decrease in offset (subtraction) with PV versus SST stimulation (p < 0.01, rank sum test). Only sessions with overall decreases in d’ relative to the control condition were analyzed. **B**. Same as A, but for change in gain (g). SST stimulation significantly decreased the gain (–0.26 ± 0.22 MI, p < 1e-4), but PV stimulation did not (gain –0.04 ± 0.17, p = 0.63). Significantly larger decreases in gain (division) with both SST vs PV stimulation (p < 1e-3). **C**. Affine model for contrast responses in RS neurons (124 neurons with PV stim.; 112 neurons with SST stim.). Both PV (–0.87 ± 0.13 MI, p <1e-8) and SST (–0.55 ± 0.45 MI, p <1e-4) stimulation decreases the offset. Significantly larger decreases in offset with PV versus SST stimulation (p = 0.04). Only RS neurons with overall decreases in spiking activity relative to the control condition were analyzed. **D**. Same as C, but for gain change. While both PV (–0.60 ± 0.40 MI, p < 1e-12) and SST (–1.00 ± 0.00 MI, p < 1e-14) stimulation decreased the gain, there is significantly larger decreases in gain with SST versus PV stimulation (p < 1e-2).

**Figure S5.**
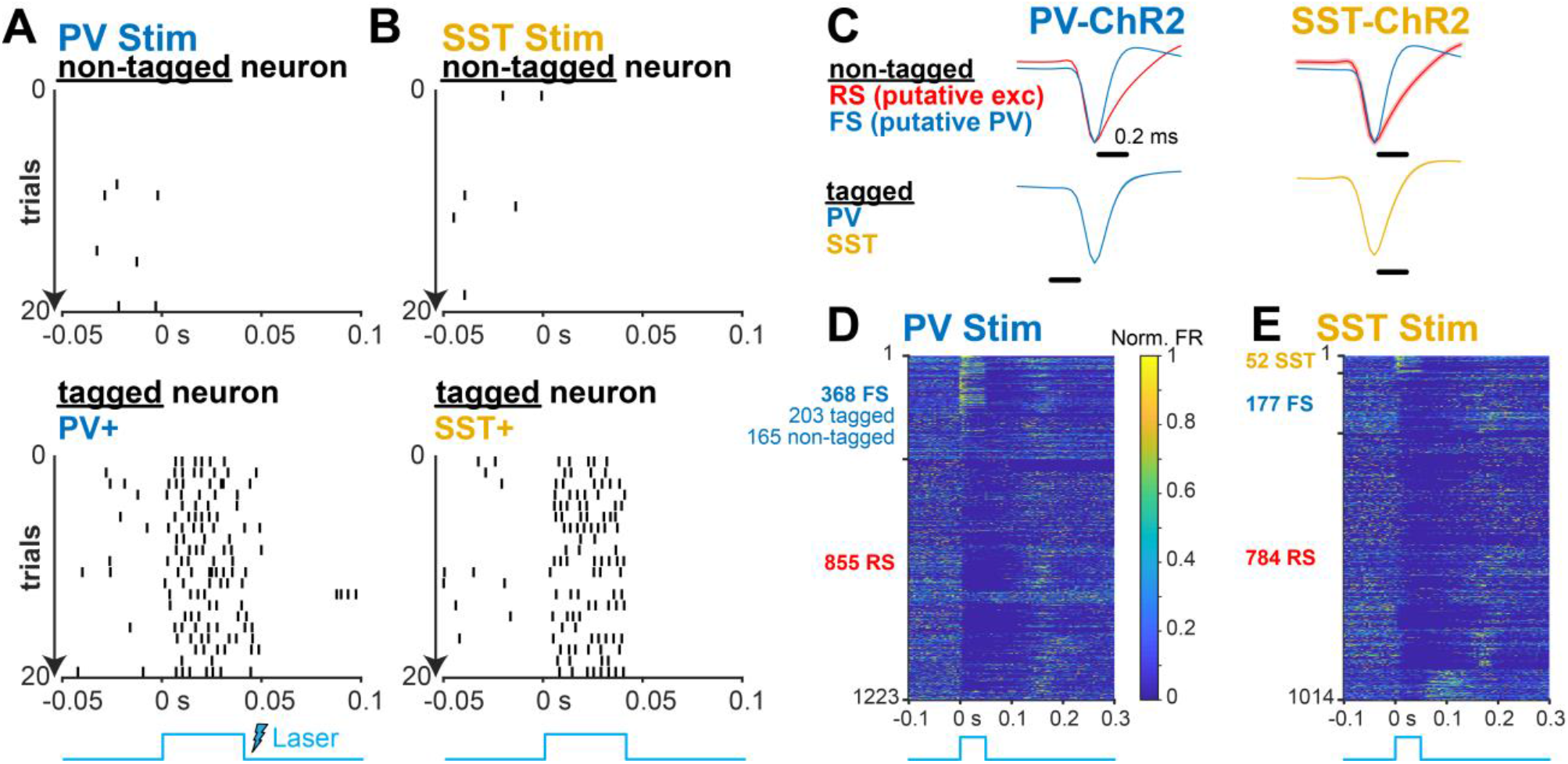
Neuron identification by optotagging and waveform profile. **A**. Example non-tagged (top) neuron and opto-tagged (bottom) neuron responses to a brief laser pulse (1.7 mW, 40 ms duration) positioned over the recording site (local stimulation, where recording and stimulation site were the same). In PV-ChR2 mice, neurons were classified as opto-tagged PV neurons if they statistically increased spiking activity rapidly (<10 ms) to a strong laser pulse. **B**. Same as A for SST-ChR2 mice. Bottom shows example opto-tagged SST neuron. **C**. Non-tagged neurons were classified as fast-spiking (FS) putative PV interneurons or regular-spiking (RS) putative excitatory neurons based on waveform width (FS < 0.57 ms, RS > 0.57 ms). Left, 855 RS neurons, and 368 FS neurons (203 opto-tagged PV neurons; 165 non-tagged) were identified in PV-ChR2 mice. All tagged and non-tagged FS neurons had narrow waveforms and were grouped together for analysis. Right, 785 RS, 177 FS, and 52 opto-tagged SST neurons were identified in SST-ChR2 mice. **D**. Responses of all neurons during PV stimulation (normalized to max firing rate). Most FS neurons (both tagged and non-tagged) increased activity to stimulation shortly after onset (203 neurons) while a smaller fraction decreased activity (165 neurons). RS neurons (855 neurons) typically decreased their responses. **E**.Same as C for SST stimulation. SST stimulation increases SST (opto-tagged) activity while simultaneously decreasing FS and RS activity.

**Figure S6.**
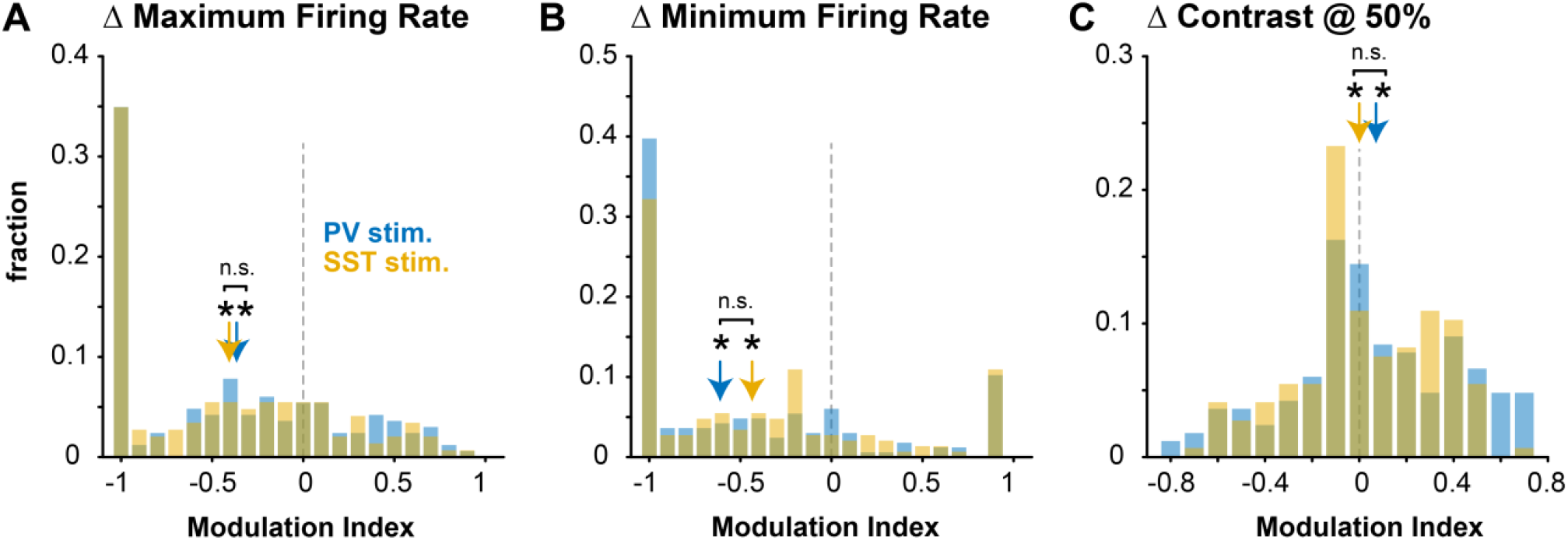
Changes in neural contrast response function (CRF) parameters. **A**. Both PV (−0.37 ± 0.51 MI, median ± mad, p < 1e-10, sign rank test) and SST (−0.41 ± 0.49 MI, p < 1e-11) stimulation significantly reduce maximum firing rate, with no difference between each other (p = 0.21, rank sum test). **B**. Both PV (−0.61 ± 0.55 MI, p < 1e-9) and SST (−0.44 ± 0.54 MI, p < 1e-6) stimulation significantly reduce the minimum firing rate, with no difference between each other (p = 0.19). **C**. Both PV (0.07 ± 0.28 MI; p < 1e-3) and SST stimulation (0.0001 ± 0.2384 MI; p = 0.01) significantly increase contrast needed for 50% firing (C_50_), with no difference between each other (p = 0.14). The increase in C_50_ in laser versus control trials (DC_50_, not normalized by sum as in MI) was not different between each other (PV: 0.02 ± 0.09, SST: 0 ± 0.08, p = 0.17).

**Figure S7.**
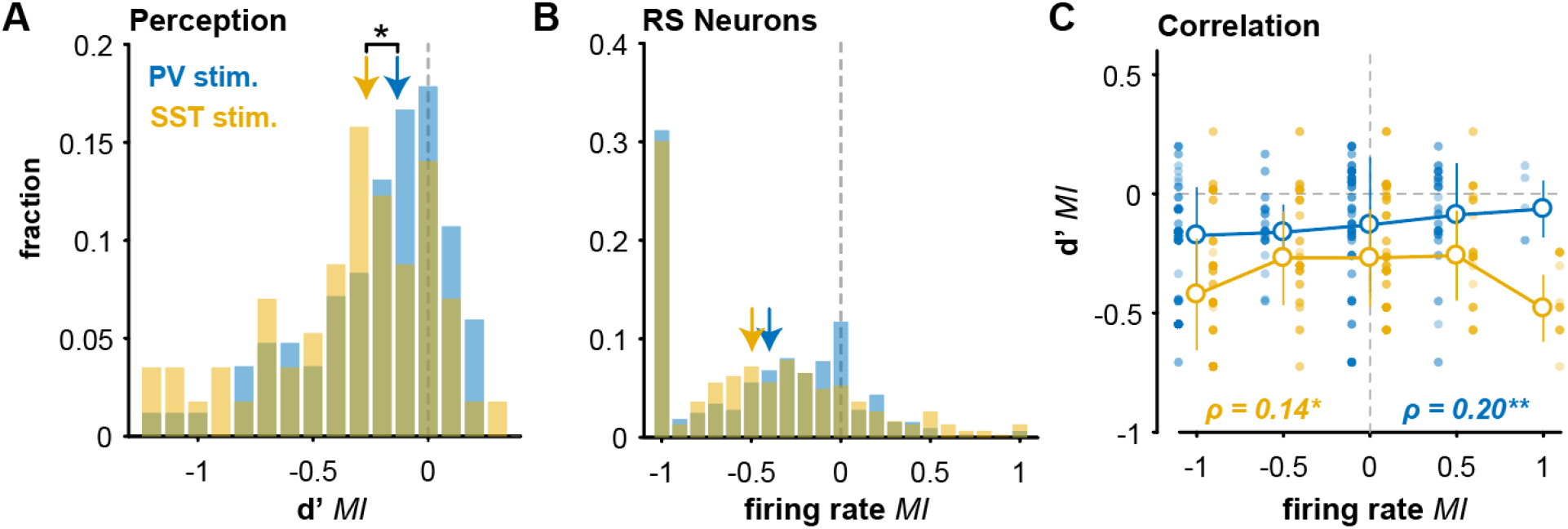
RS spiking activity and perceptual performance have correlated changes with distal PV or SST stimulation. **A**. The overall d’ decreases for both PV (−0.13 ± 0.25 MI, median ± mad, p <1e-7, 1-tail sign rank test) and SST stimulation (−0.27 ± 0.33 MI, p <1e-7), with a significantly greater reduction with SST stimulation (p = 0.02). **B**. RS activity is reduced with both PV (−0.40 ± 0.40 MI, p <1e-36) or SST stimulation (−0.50 ± 0.42 MI, p <1e-31), with no significant difference between groups (p = 0.29). **C**. Changes in RS spiking were correlated with perceptual changes for both PV (ρ = 0.20, p <1e-3, Spearman’s rank correlation) and SST stimulation (ρ = 0.14, p = 0.02).

**Figure S8.**
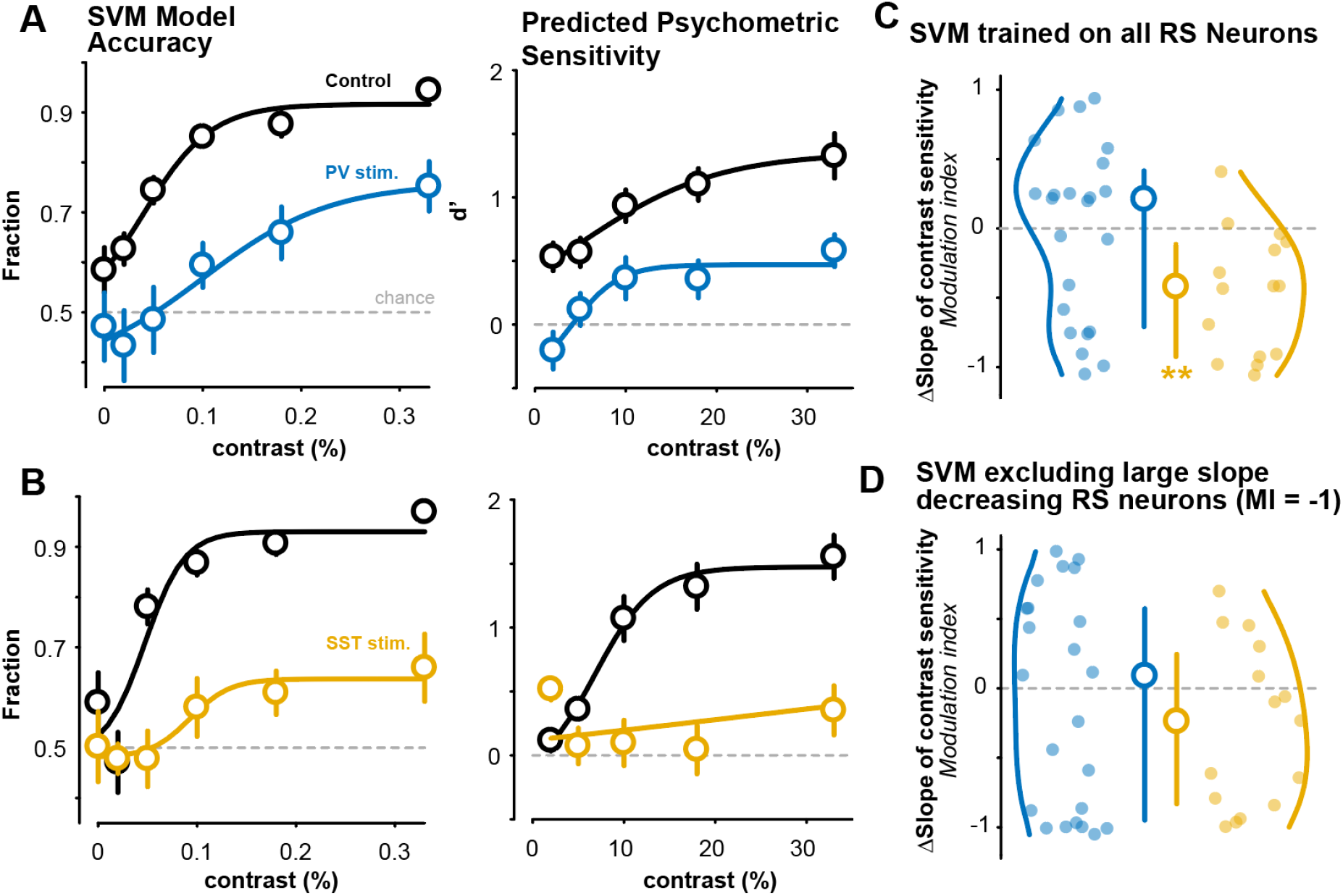
Decoder analysis using RS activity predicts psychometric contrast sensitivity changes during distal stimulation. **A**. *Left*, An SVM was trained for each session () only on control trials using all RS neurons (see Methods) and trial outcome, with no contrast information per trial. The SVM can predict behavioral responses above chance levels, and with emergent contrast dependency during the control condition (black). The SVM was then tested on the held-out PV stimulation trials within the same session (blue, left). Plots show *Right*, reconstruction of the psychometric response curve from the SVM model. **B**. Same as A, for SST stimulation. **C**. Predicted psychometric curves shows significant decrease in slope with SST stimulation (−0.42 ± 0.37 MI, median ± mad; p < 0.01, 1-tail sign rank test), but not PV stimulation (0.21 ± 0.54 MI; p = 0.98). Highly similar results were obtained with a single classifier trained on 50% of the control trials randomly sampled across all sessions, then tested on the 50% held out control trials, and 100% held out stimulation trials (slope MI: SST = -0.52 ± 0.31, median ± mad, p < 0.01, sign rank; PV = -0.32 ± 0.64, p = 0.26). There was no difference in the calculated slope MI for the within session versus across sessions SVM, for PV (p = 0.20) or SST (p = 0.42) stimulation groups. **D**. Removing RS neurons with complete contrast sensitivity loss (MI = -1) diminishes the change in slope (less divisive change) on SST stimulation trials (SST: -0.24 ± 0.51 MI, p = 0.09; PV: 0.09 ± 0.70 MI, p = 0.35).

**Figure S9.**
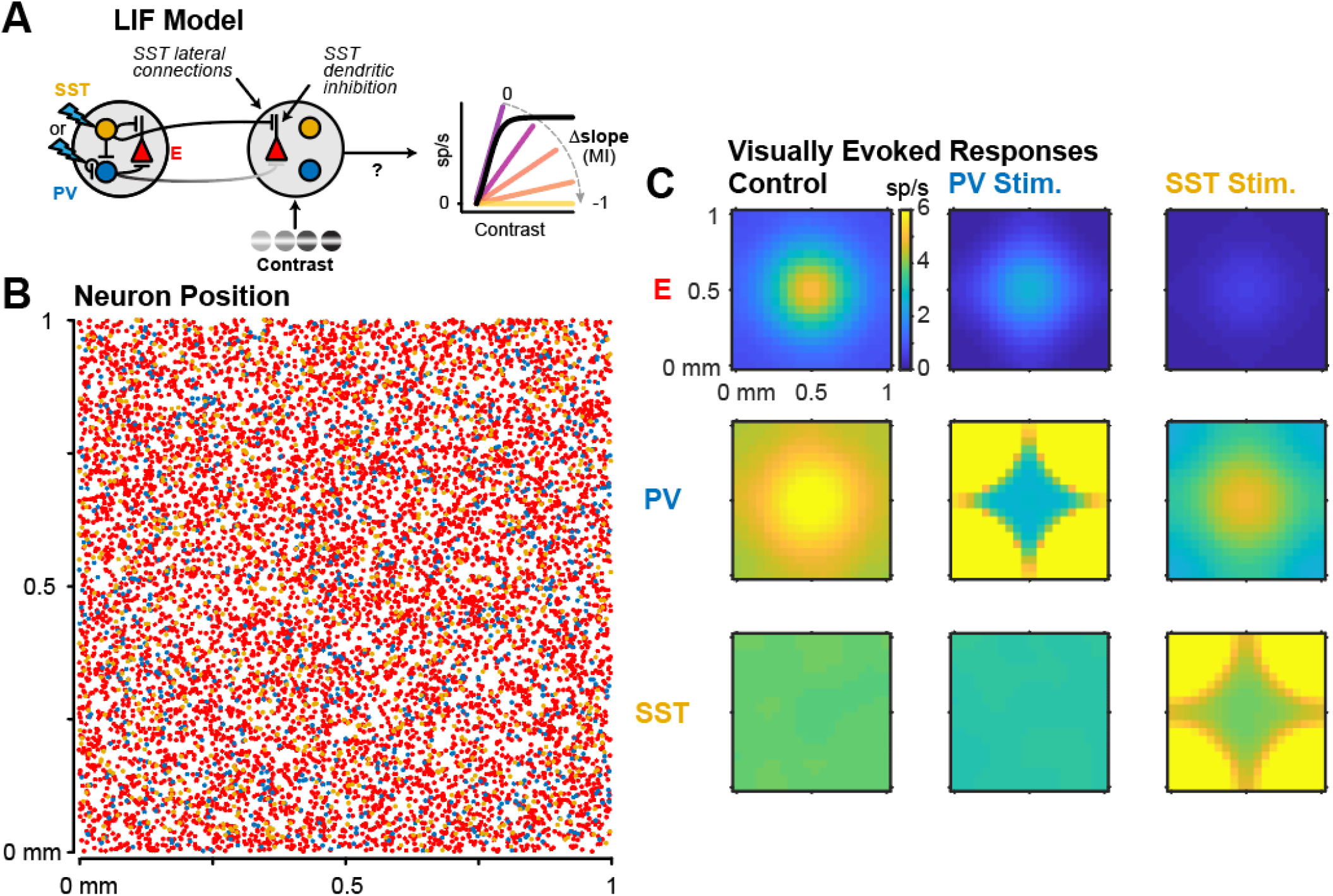
Spatial structure of LIF model. **A**. Schematic of the LIF model used to measure the relationship between lateral inhibitory connectivity and modulation of visually evoked responses across contrasts. **B**. E (red, 8k), PV (blue, 1k), and SST (orange, 1k) neurons within a 1mm^2^ grid. **C**. Spatial activity patterns of E, PV, and SST neurons during visual stimulus presentation for control (left), PV stimulation (middle), and SST stimulation (right).

**Figure S10.**
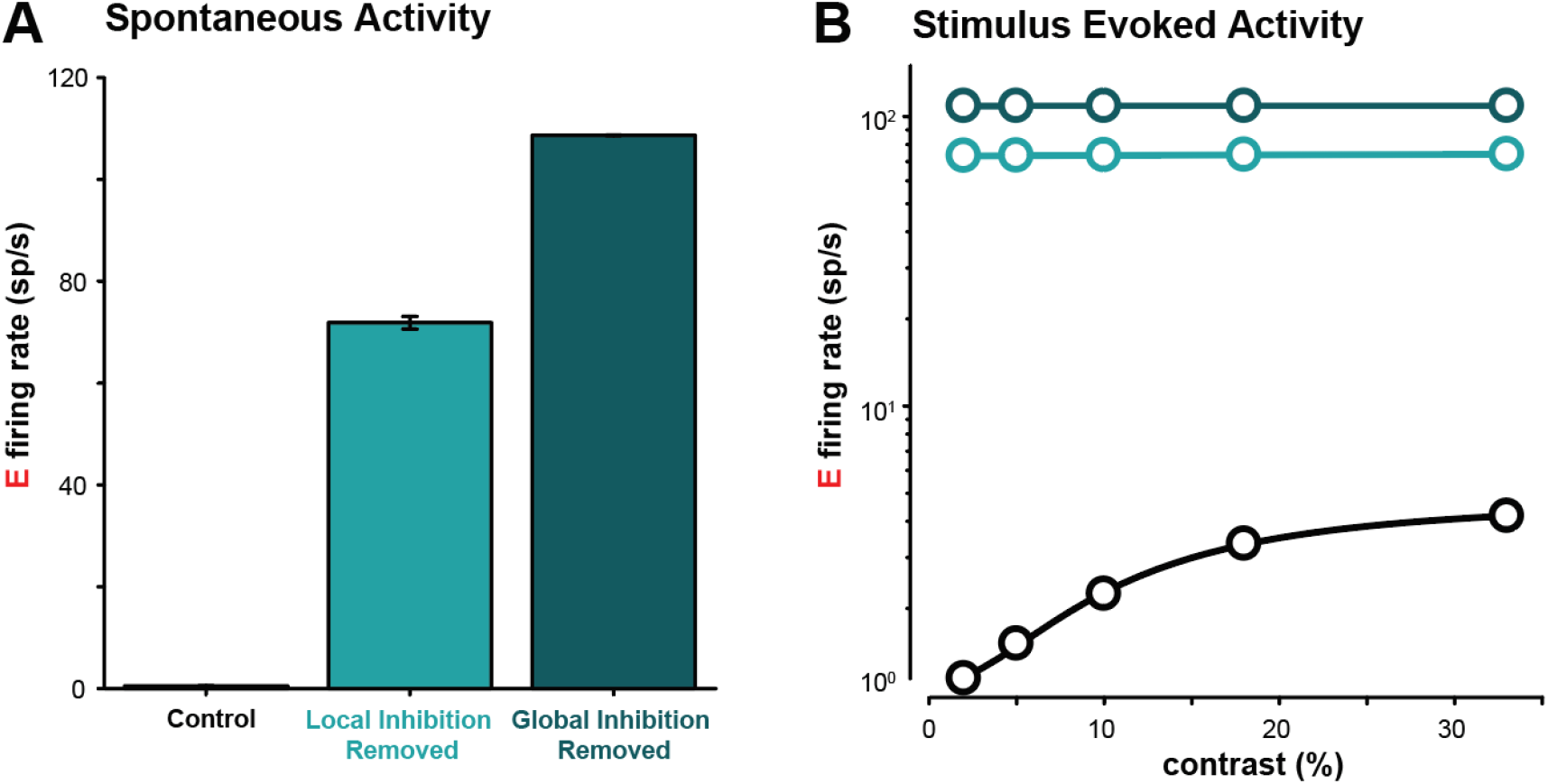
Removing inhibition de-stabilizes the LIF network. **A**. Excitatory activity increases >50-fold when inhibition is removed locally (site of visual input), or globally throughout the whole network. **B**. Visually evoked responses when inhibition is removed locally and globally. Compare to evoked activity rates in intact model (black).

**Figure S11.**
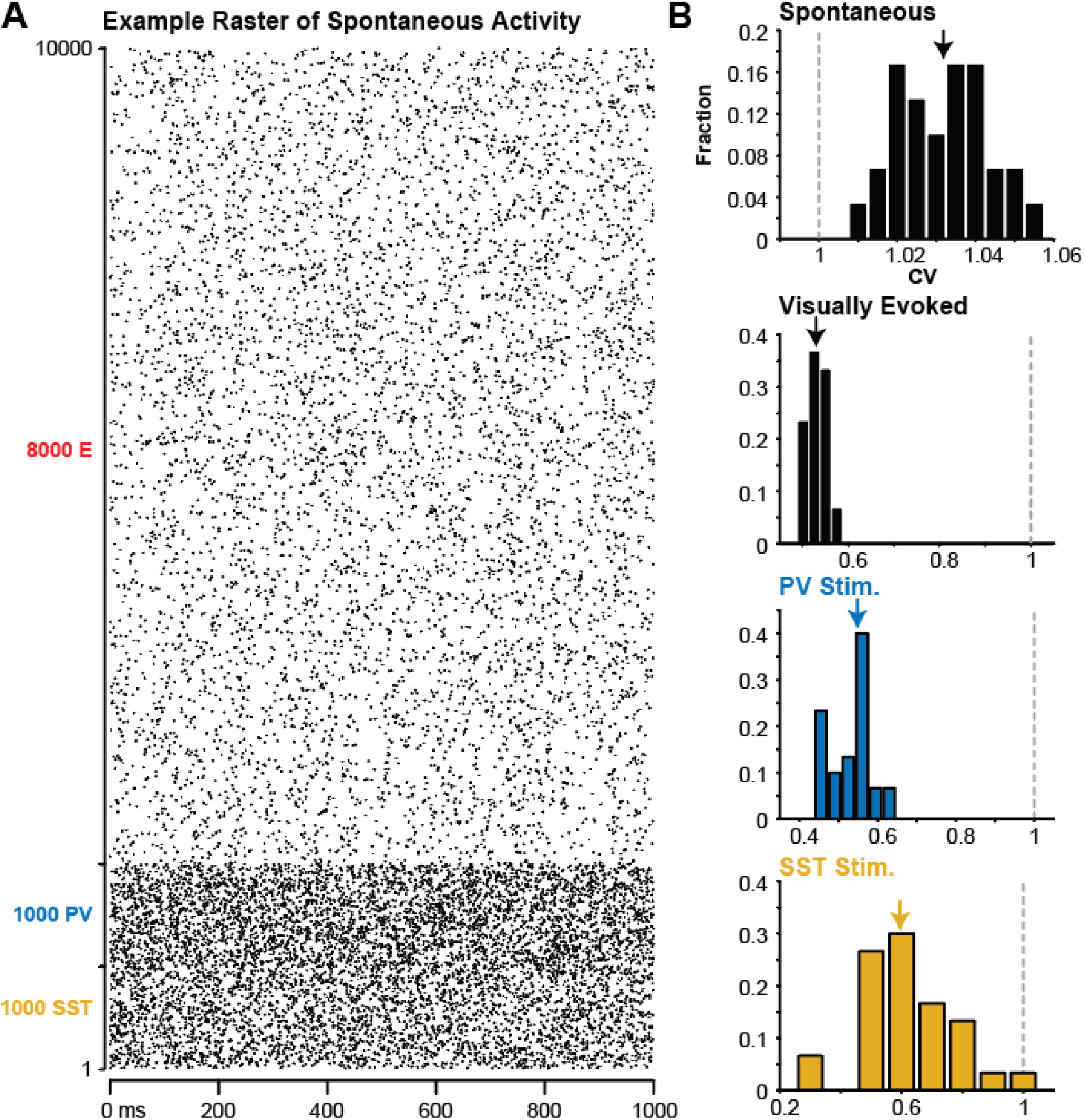
LIF model dynamics. **A**. Raster plot of E, PV, and SST neurons during spontaneous activity (1 s) in LIF **B**. *Top*, Coefficient of variation (CV) of inter-spike intervals for E neurons during spontaneous activity (30 simulations). CV > 1 typical for asynchronous irregular spiking across population (CV = 1.03 ± 0.01, median ± mad, p < 1e-5, sign rank), similar to CV of the experimentally recorded RS neurons (1.12 ± 0.58, p<1e-14). *Bottom*, RS activity becomes more synchronous during visual stimulation (Control: 0.53 ± 0.02, p < 1e-5; PV stim.: 0.55 ± 0.04, p < 1e-5; SST stim.: 0.60 ± 0.12, p < 1e-5).

**Figure S12.**
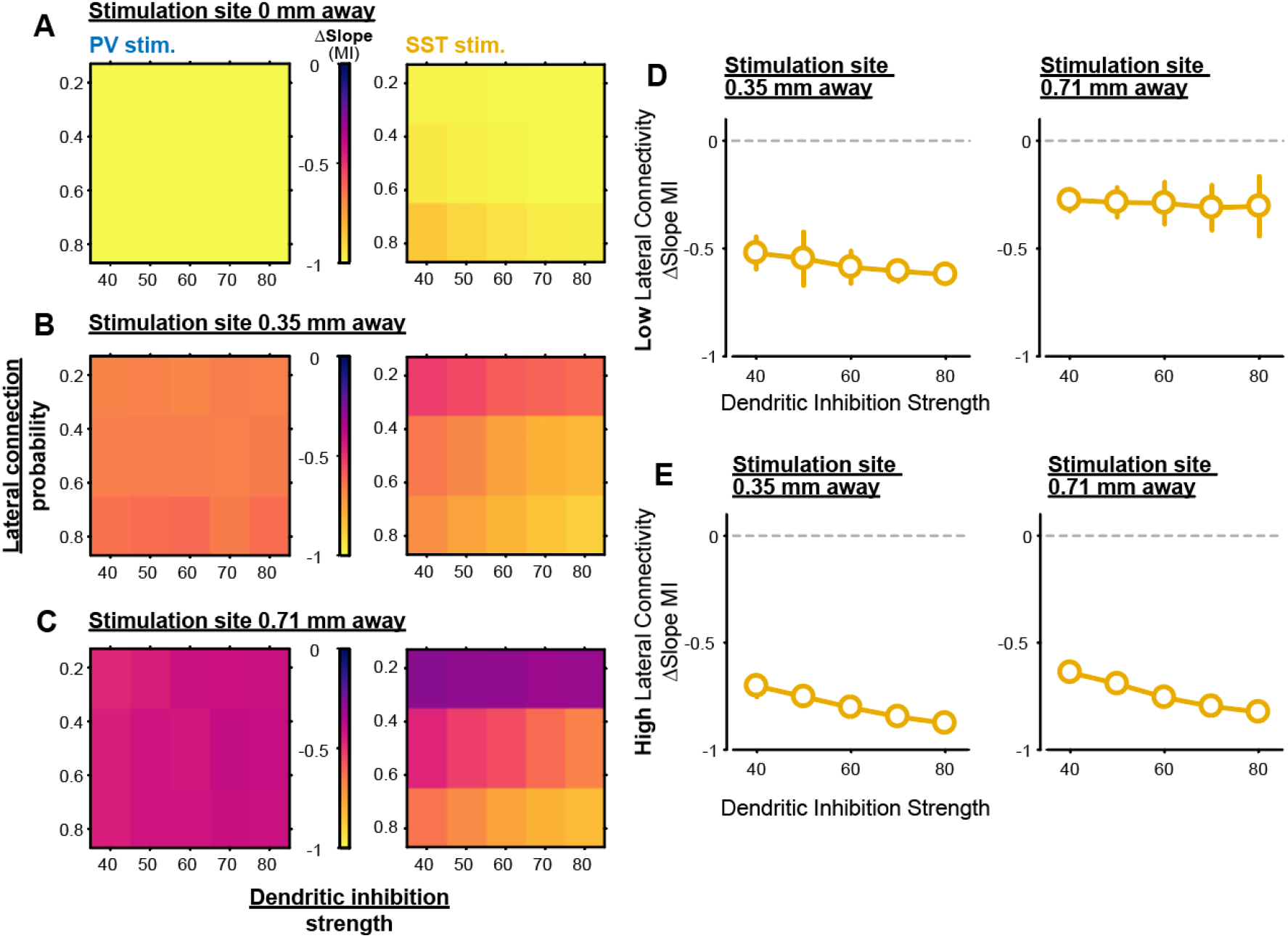
The effect of lateral connectivity and dendritic inhibition strength across stimulation distance in a LIF model. **A**. Heatmap showing the divisive effect on the excitatory contrast responses (slope MI) for different lateral SST connection probability (y axis)and SST dendritic inhibition strength (x axis), when the stimulation site (PV – left, SST – right) overlaps with the site of visual input (0mm away). **B**. Same as A, but when the stimulation site is 0.35 mm away from the site of visual input. **C**. Same as A, but when the stimulation site is 0.71 mm away from the site of visual input. **D**. When there is low lateral connectivity, dendritic inhibition strength by itself has little effect on the slope MI if the SST stimulation site is near (0.35 mm, left) or far away (0.71 mm) from the site of visual input. **E**. Same as D, but with high SST lateral connectivity. Increasing dendritic inhibition strength elicits larger decreases in the slope MI more evenly across stimulation locations in the presence of high SST lateral connectivity.

**Figure S13.**
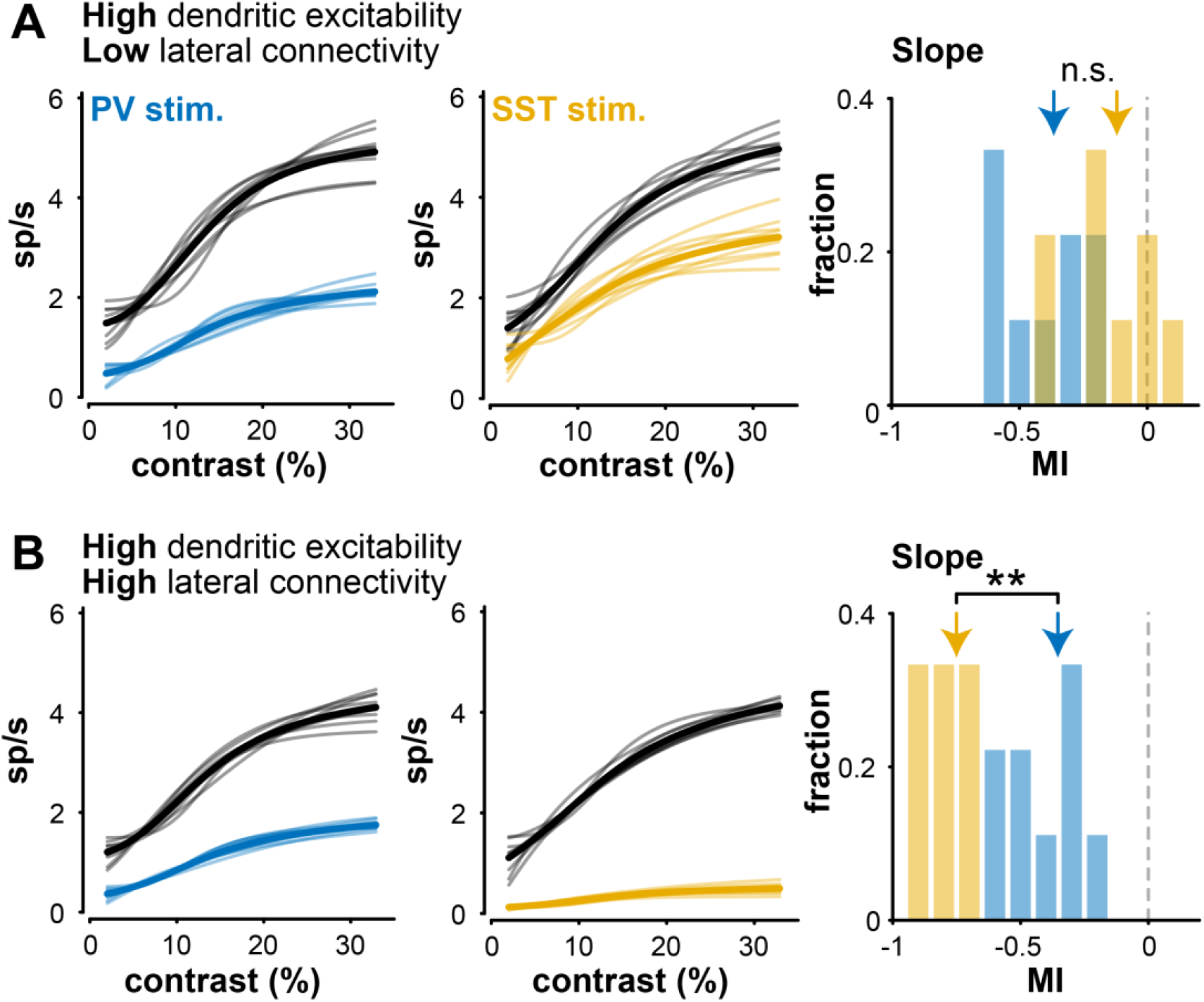
Increasing dendritic excitability is insufficient for driving divisive slope changes in LIF model. **A**. LIF model with high dendritic excitability but low lateral connectivity. High dendritic excitability activates a larger fraction of SST neurons, but is insufficient for driving larger divisive effects with SST stimulation (SST: –0.10 ± 0.11; PV: –0.31 ± 0.16; p = 0.99). **B**. Same as A, but restoring high lateral SST connectivity. Greater divisive changes in the contrast sensitivity curve emerge for SST versus PV stimulation (SST: –0.72 ± 0.03, median ± MAD; PV: –0.37 ± 0.14; p < 1e-3, 1-tail Wilcoxon rank sum test).

**Figure S14.**
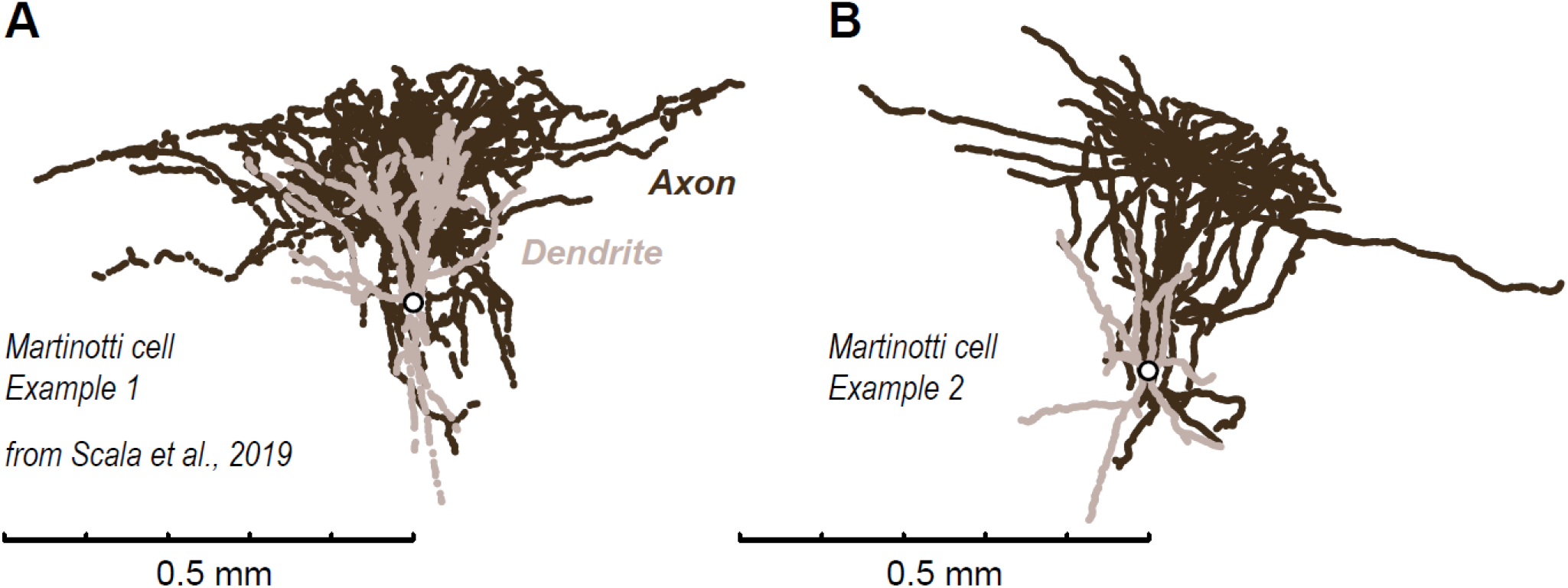
SST Martinotti cells in V1 show extensive lateral projections. **A, B**. Two example SST Martinotti cells from mouse V1, from Scala et al., 2019. Axons ramify laterally > 0.4 mm. Scale bars (0.5mm) aligned to soma. Data accessed and plotted from Zenodo link in paper^1^. ^1^Scala et al., Layer 4 of mouse neocortex differs in cell types and circuit organization between sensory areas. *Nat Commun* **10**, 4174 (2019). https://doi.org/10.1038/s41467-019-12058-z https://doi.org/10.5281/zenodo.3336165

**Figure S15.**
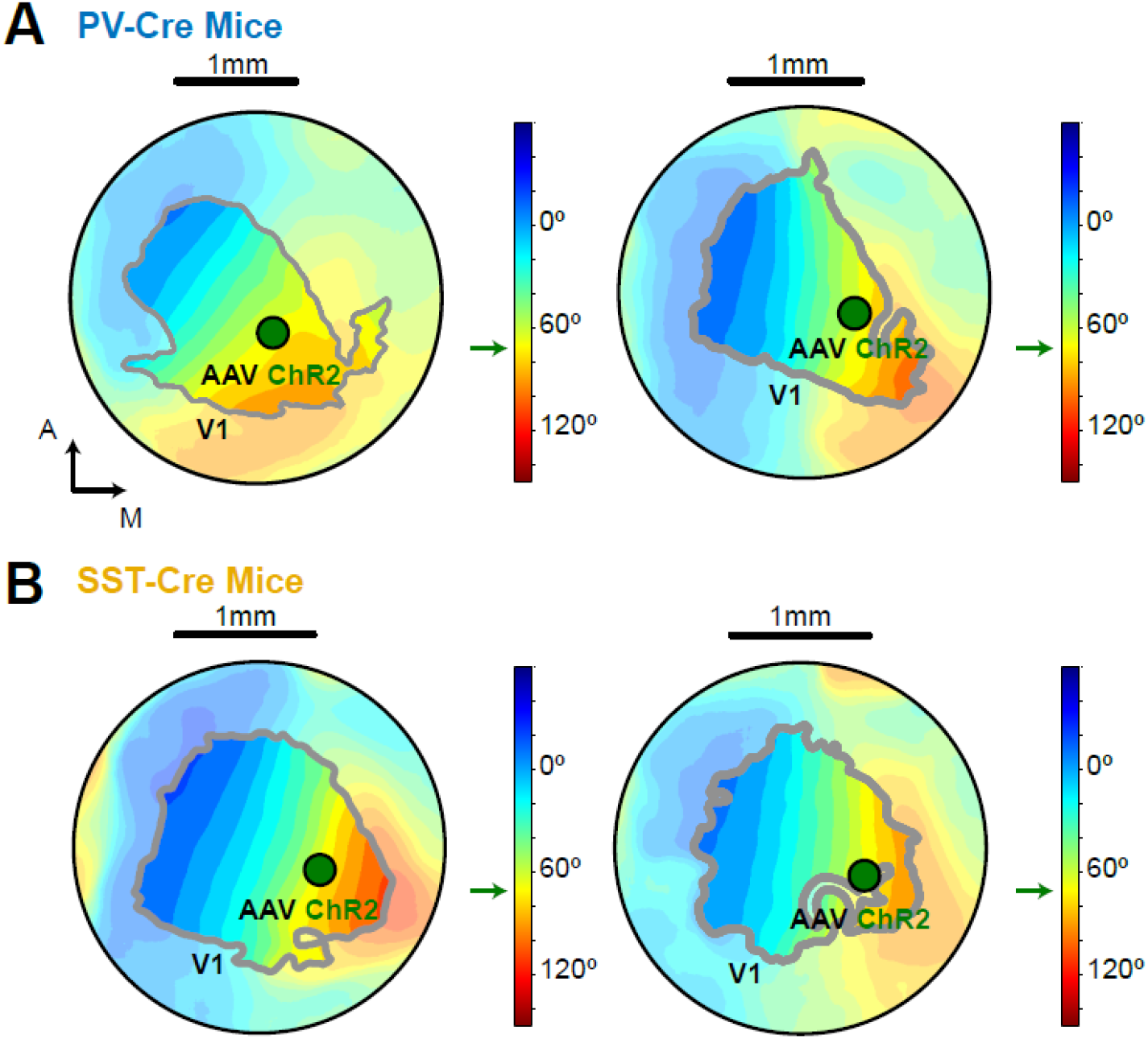
Retinotopically targeted viral injections. **A, B**. AAV-Flex-ChR2 injected into monocular V1 (∼70°, identified via intrinsic imaging of retinotopy) in PV- or SST-cre mice. Same mice with recordings in binocular V1 in Fig.S20, S23.

**Figure S16.**
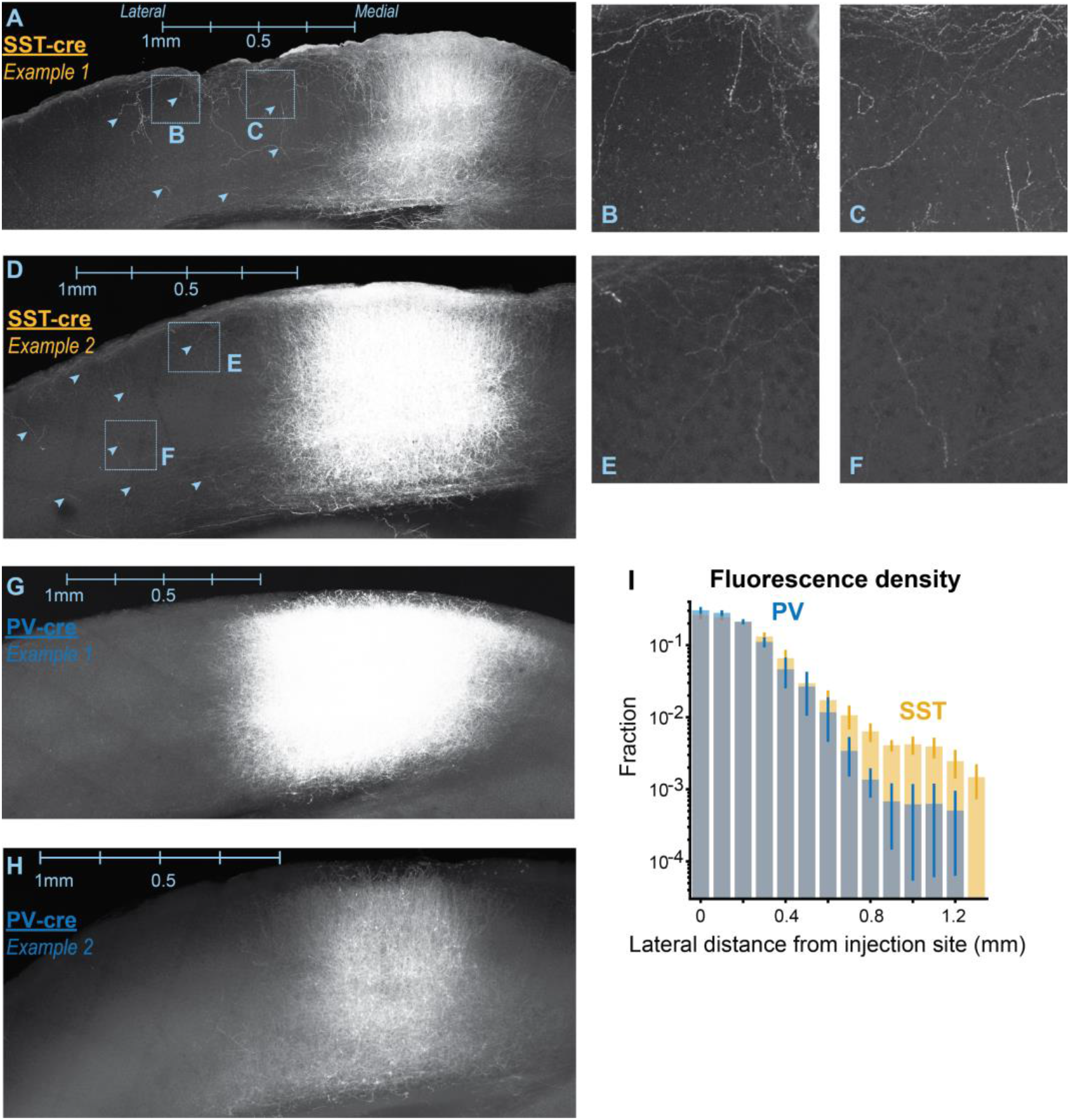
Lateral projections of SST neurons from monocular V1. **A**. Coronal section of AAV Flex EYFP injection in SST-Cre mouse, optically targeted to monocular region of V1 in left hemisphere (see Fig. S15; same area targeted for optogenetic stimulation *in vivo*). Note projections in all layers laterally towards binocular V1 (arrowheads) and in L1. Scale bar aligned to most lateral cell body near injection site. **B, C**. Expanded insets from A. Fibers show “beads on a string” consistent with axonal varicosities. Inset 0.5mm across. **D-F**. Same as A-C, in a second SST-cre mouse. **G, H**. Same as A, D, but in two example PV-cre mice. No evidence of lateral axonal projections beyond injection site, nor in L1. **I**. SST mice (n=2) show more extensive lateral fluorescence signal (likely from axons) than PV mice (n=2; *p < 1e-17*, Kolmogorov-Smirnov test). **I**. SST mice (n=2) show more extensive lateral fluorescence signal (likely from axons) than PV mice (n=2; *p < 1e-17*, Kolmogorov-Smirnov test).

**Figure S17.**
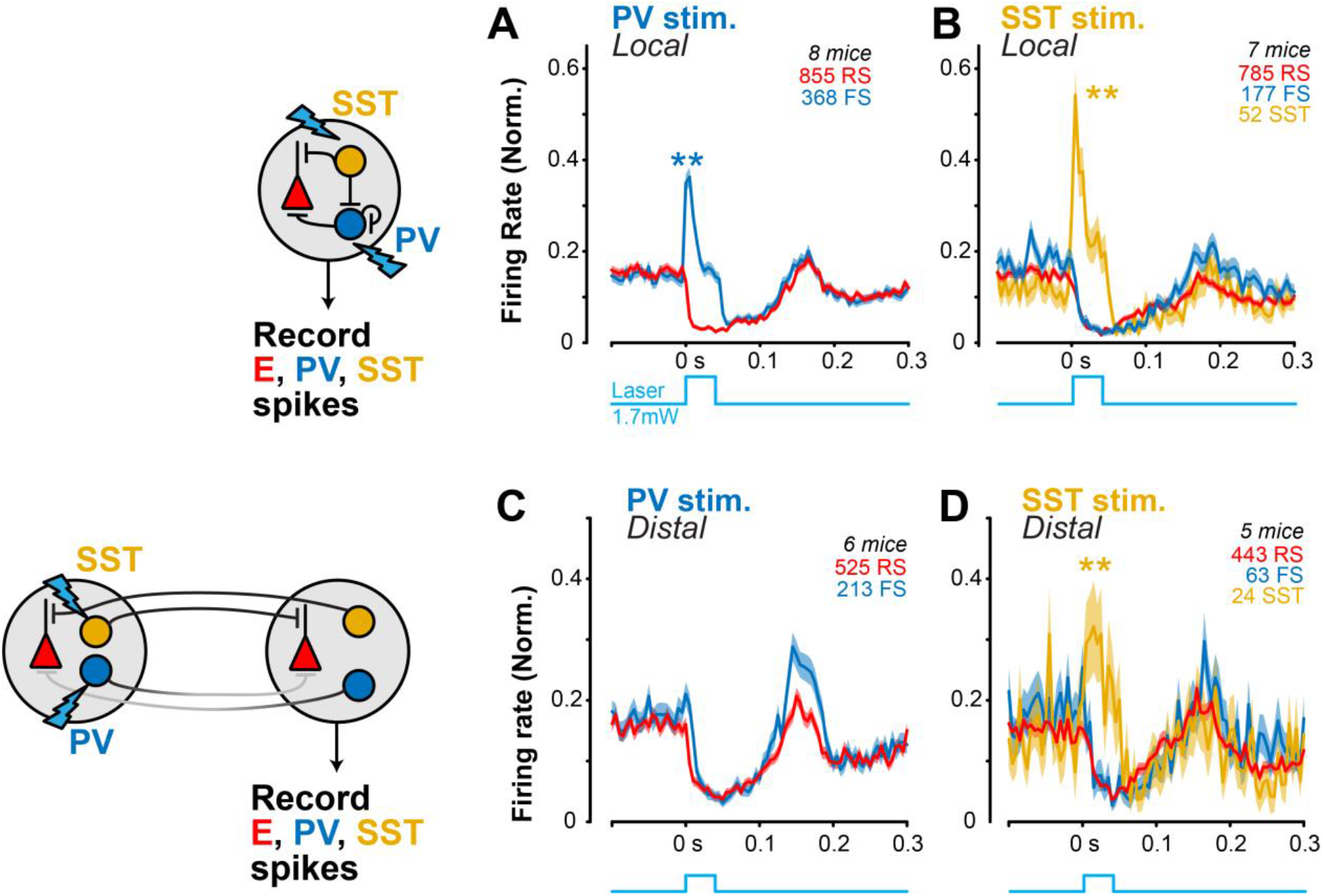
Single unit activity during local and distal PV and SST stimulation. **A**. Local PV stimulation (brief square pulse, 40 ms duration, 1.7 mW) rapidly increases FS activity (0.14 ± 0.04 MI, mean ± SEM, *p* < 0.01, Wilcoxon signed-rank test) while simultaneously decreasing RS activity (–0.63 ± 0.02 MI, *p* < 1e–93). **B**. Local SST stimulation increases SST activity (0.49 ± 0.08 MI, *p* < 1e–5), which inhibits FS (–0.45 ± 0.04 MI, *p* < 1e–14) and RS activity (–0.50 ± 0.02 MI, *p* < 1e–60). **C**. Distal PV stimulation decreases both FS (–0.28 ± 0.04 MI; *p* < 1e–7, 213 FS neurons) and RS activity (–0.46 ± 0.02 MI; *p* < 1e–40, 525 RS neurons). **D**. Distal SST stimulation decreases RS (–0.33 ± 0.03 MI; *p* < 1e–17, 443 RS neurons) and FS activity (–0.30 ± 0.07 MI; *p* < 1e–3, 63 FS neurons), but increases SST activity (0.26 ± 0.09 MI; *p* = 0.01, 24 SST neurons).

**Figure S18.**
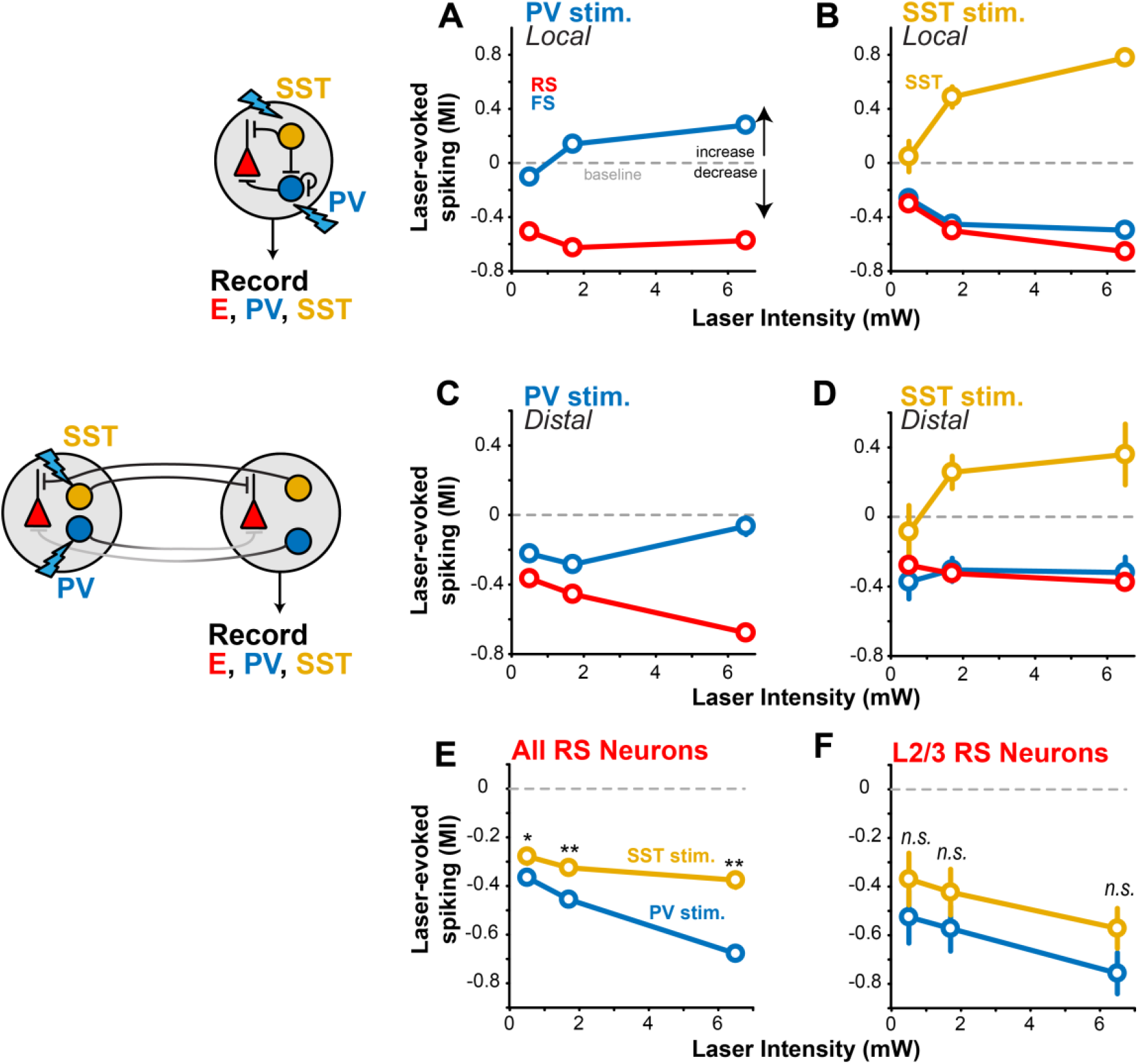
PV or SST stimulation effects across laser intensities. **A**. During local PV stimulation, neural activity scales with laser intensity (0.5 – 6.5 mW). FS activity increases and RS activity decreases with stronger laser intensities. Same neurons as in Fig. S17. Moderate power of 1.7mW was used for all main results in study. **B**. Same as A for local SST stimulation. SST activity increases with corresponding decreases in FS and RS activity during stronger laser intensities. **C**. Same as A for distal PV stimulation. Both FS and RS activity decreased with distal stimulation across laser intensities. **D**. Same as B for distal SST stimulation. Despite the laser stimulating a distal cortical site (0.8mm away), SST activity increased across laser intensities while FS and RS activity decreased with distal SST stimulation. **F**. Averaged across all layers, RS activity is more suppressed with distal PV than SST stimulation (p < 0.05 for all laser intensities). **G**. In L2/3 RS neurons, there is no significant difference between distal PV (n=36 RS) and SST (n=48 RS) stimulation (0.5 mW: p = 0.26; 1.7 mW: p = 0.32; 6.5 mW: p = 0.11). The firing rate decrease during stimulation (raw FR) was greater only at 6.5 mW (p = 0.048), but not at lower intensities (1.7 mW: p = 0.62; 0.5 mW; p = 0.38).

**Figure S19.**
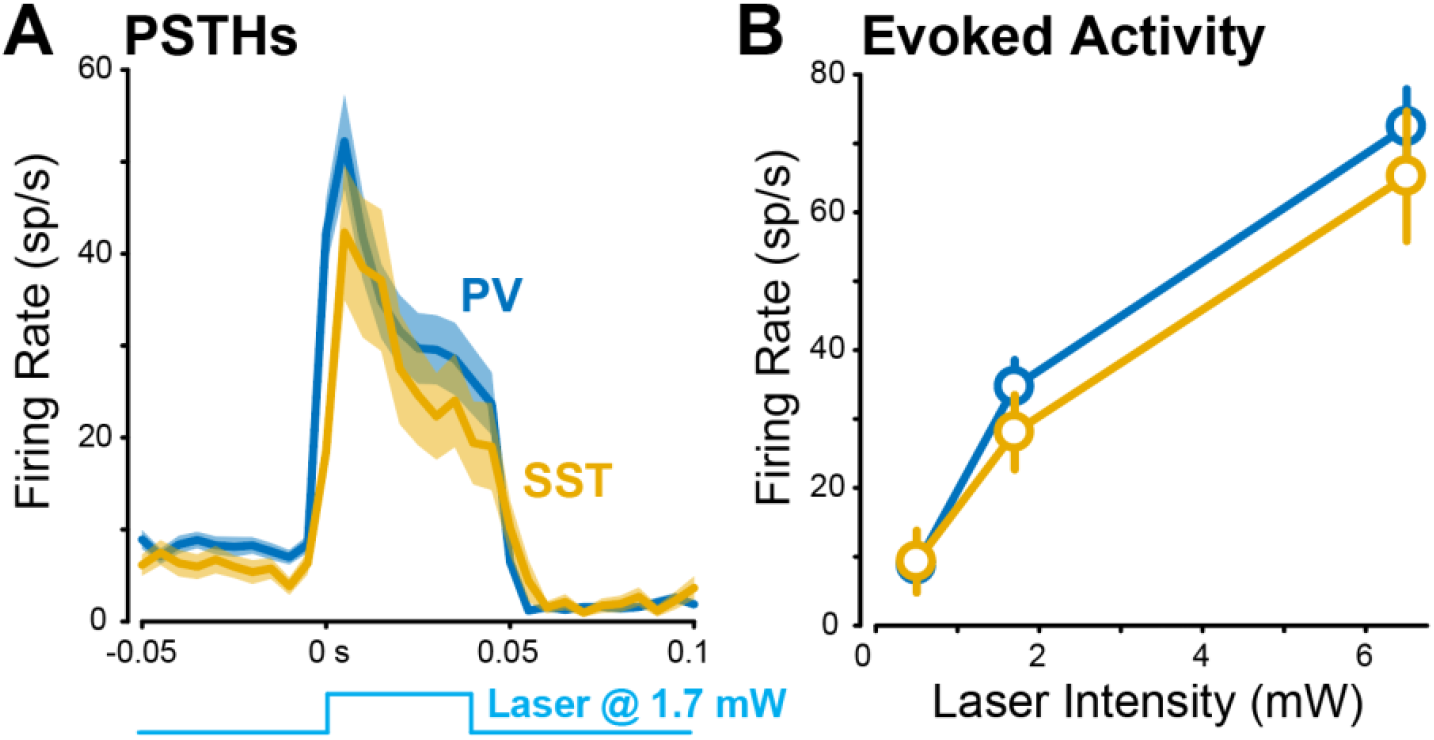
SST neurons not more photo-excitable than PV neurons. **A**. PSTHs of SST (n=52) or PV (n=203) neuron activity with local optogenetic stimulation (1.7 mW). Only “optotagged” neurons. Mean ± SEM. No difference in latency of activation (PV: 5.0 ± 6.4 ms; SST: 5.0 ± 7.1 ms; p = 0.24, 1-tail rank sum). **B**. Across intensities, SST neurons had similar evoked activity to PV neurons (0.5 mW: p = 0.237; 1.7 mW: p = 0.237; 6.5 mW: p = 0.485)

**Figure S20.**
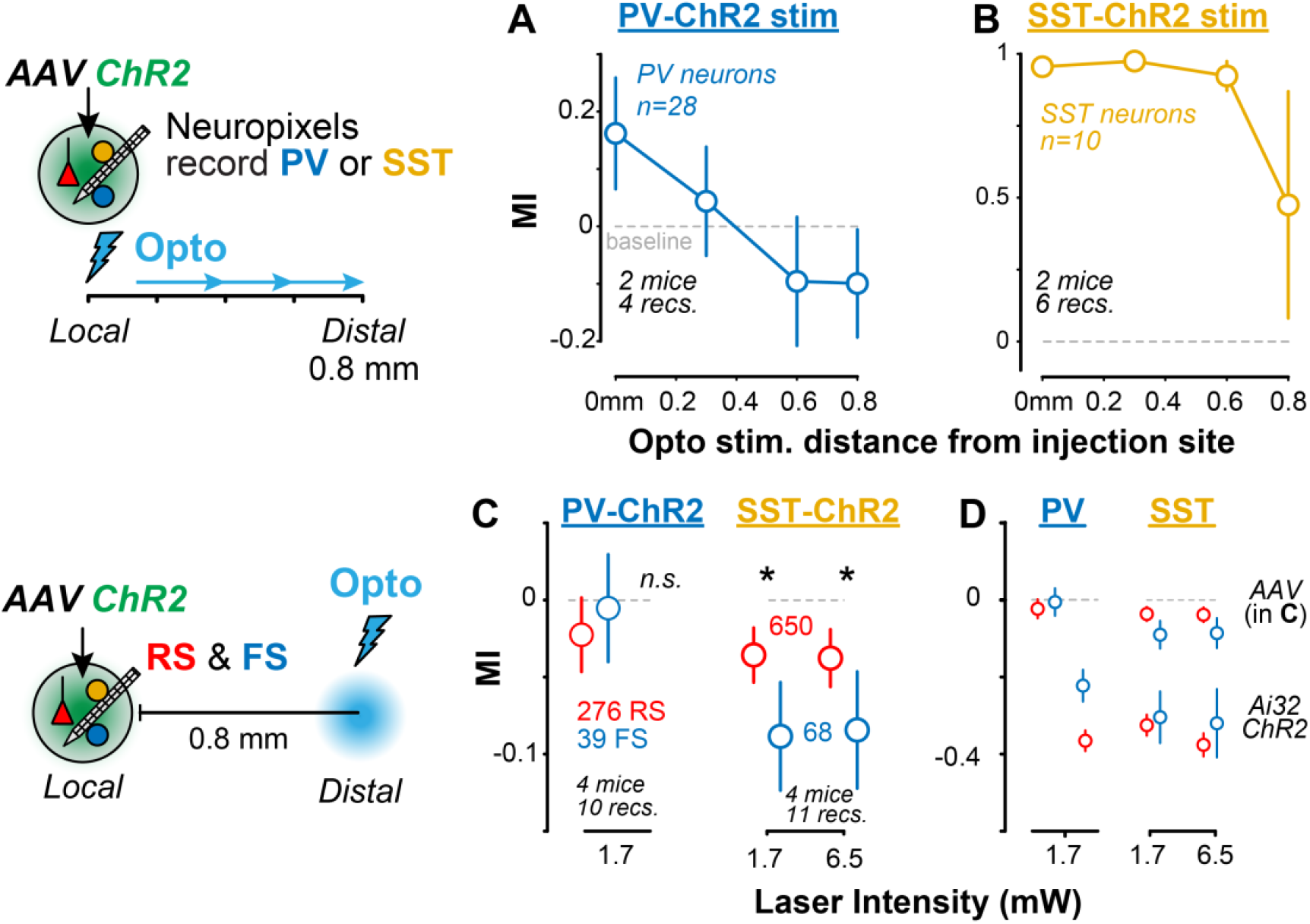
Focal ChR2 expression and distal photostimulation. **A**. AAV-Flex-ChR2 injection in PV- or SST-Cre mice, with Neuropixels recording at injection site and photostimulation (1.7mW) varying distances away. PV neurons are not activated with more distal photostimulation (y-axis, modulation index relative to control firing). **B**. In SST-Cre mice, distal photostimulation drives local SST neuron spiking. **C**. *Left*, no significant suppression in RS or FS firing with distal PV stimulation (1.7 mW, same as main experiments; p = 0.26). *Right*, significant suppression with distal SST stimulation at 1.7 mW (p = 0.01) and 6.5 mW (p = 0.045), but no difference across powers (p = 0.408). **D**. Same as C with comparison to experiments done in transgenic expressing ChR2 mice (Fig. S18).

**Figure S21.**
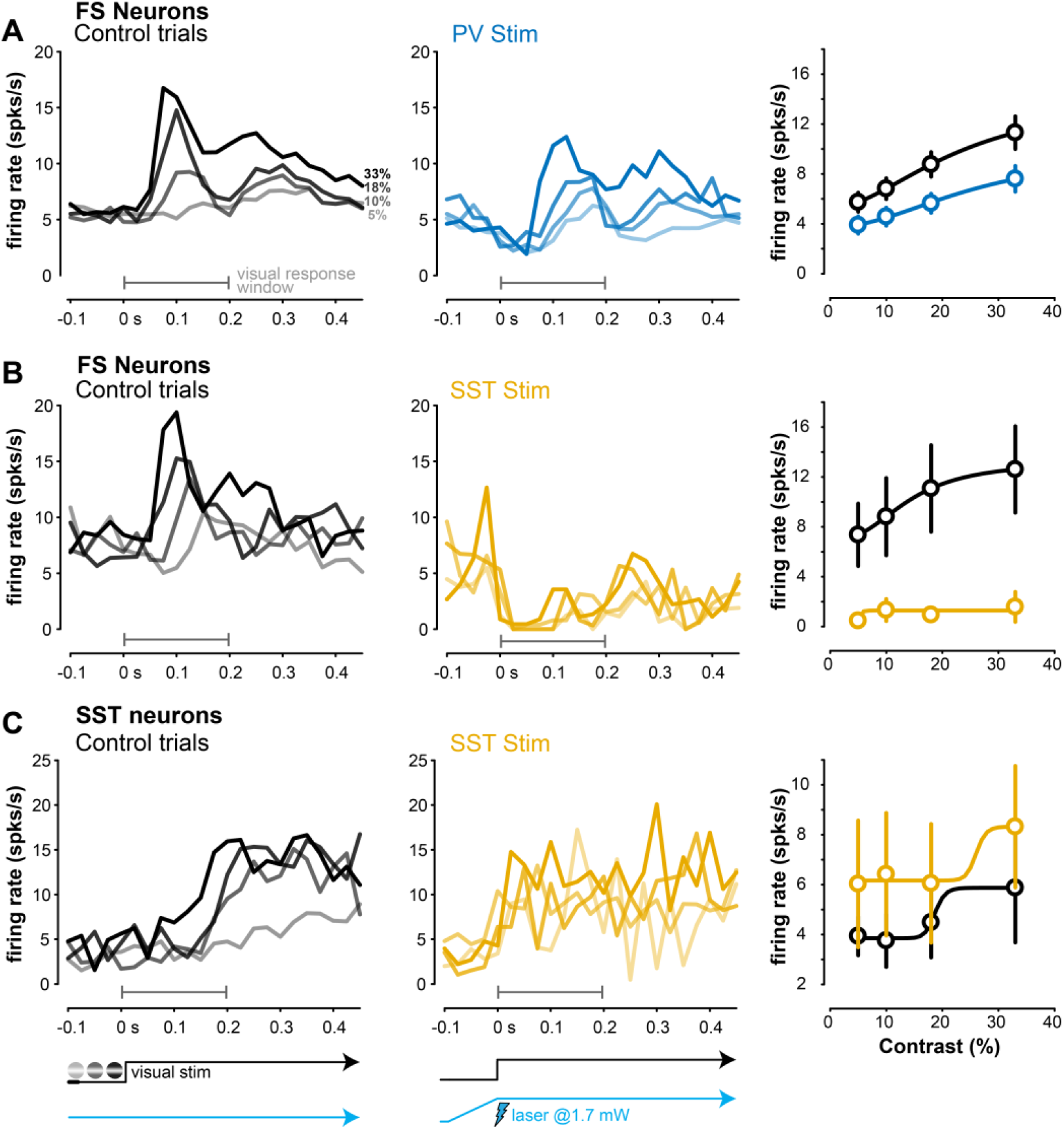
FS and SST activity during distal stimulation and perceptual behavior. **A**. FS neuron PSTHs during control (no laser, left) and distal PV stimulation (middle). Distal PV stimulation decreased the slope of FS neurons contrast response function (–0.26 ± 0.51 MI, median ± MAD; *p* < 1e-4, Wilcoxon signed rank test; 165 FS neurons). **B**. Same as for distal SST stimulation (. Distal SST stimulation strongly inhibited FS activity across all contrasts. Distal SST stimulation strongly decreased the slope of FS neuron contrast response functions (–0.71 ± 0.47 MI; *p* < 1e–2; 37 FS neurons). These effects were greater than with distal PV stimulation (*p* = 0.041, 1-tail Wilcoxon rank-sum test). **C**. PSTHs of SST neurons during control (left) and distal SST stimulation (middle). Distal SST stimulation increased SST activity across all contrasts (right; control = 4.91 ± 1.10 sp/s, mean ± SEM; SST stimulation = 8.26 ± 1.71 sp/s; *p* = 0.06; 18 SST neurons).

**Figure S22.**
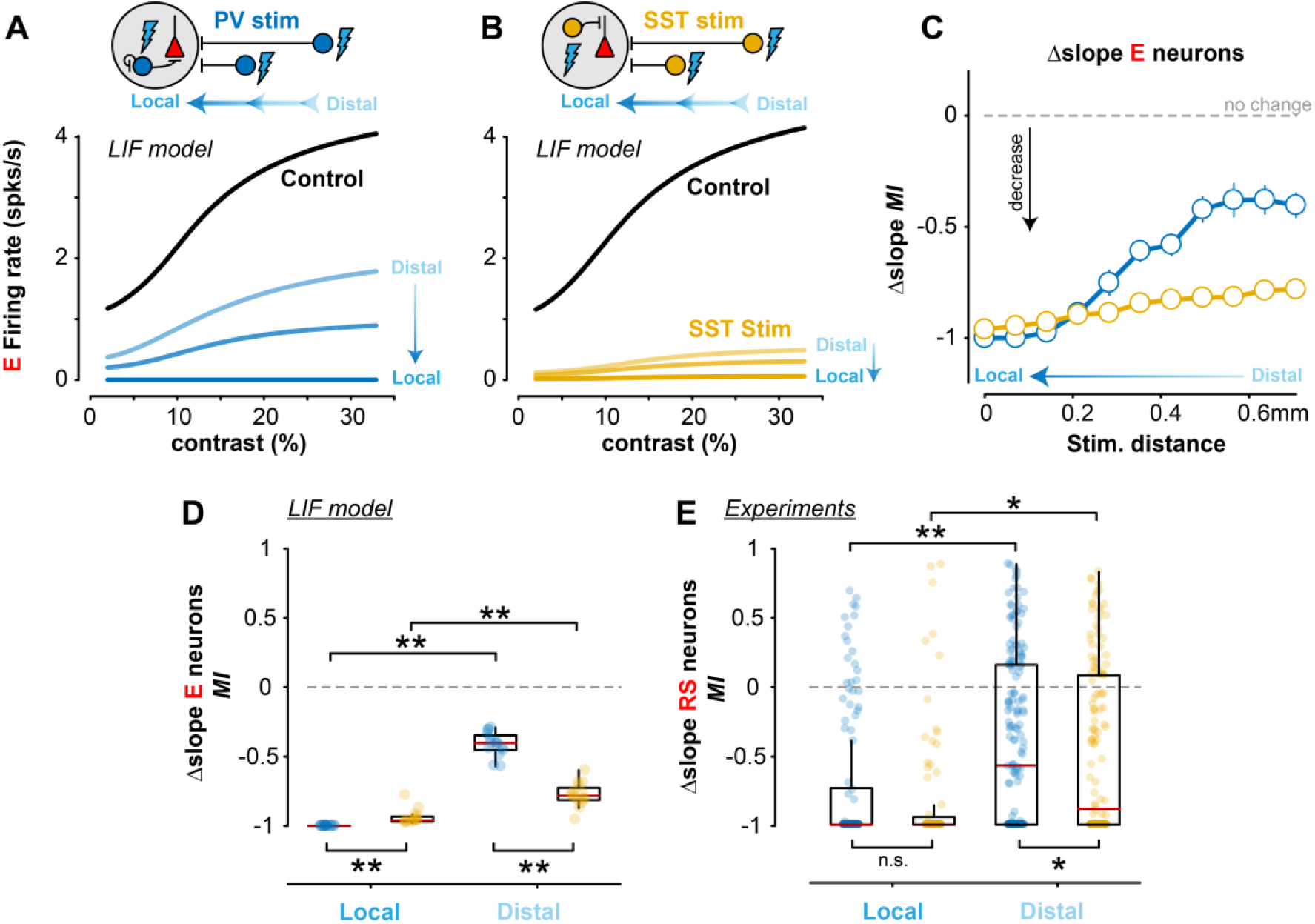
Modulation of contrast sensitivity as a function of stimulation distance. **A**. Contrast response curves of excitatory neurons in the LIF network model as the site of PV stimulation was applied at 3 varying distances away from the site of visual input. Median ± MAD plotted. **B**. Same as A for SST stimulation. **C**. Changes in the slope of E neuron contrast sensitivity function (MI, per all other analyses) as a function of stimulation distance. PV stimulation strongly reduced the slope during local stimulation, but the magnitude of effect diminished as the site of stimulation moved farther away. In comparison, SST stimulation exerted strong reductions in slope regardless of stimulation distance. 11 runs per stimulation site. Median ± MAD plotted. **D**. Changes in the slope of E neurons during local (stimulation distance = 0 mm) or distal stimulation (stimulation distance = 0.7 mm). Local PV stimulation (–1 ± 0 MI, median ± MAD, Wilcoxon sign-rank test) more strongly decreased the slope than local SST stimulation (–0.96 ± 0.04 MI; *p* < 1e–6, 1-tail Wilcoxon rank-sum test). The slope change was significantly smaller with distal vs local PV stimulation (*p* < 1e–6) as was the case with distal vs local SST stimulation (*p* < 1e–4). However, distal SST stimulation still reduced the slope significantly more (–0.78 ± 0.06 MI) than distal PV stimulation (–0.40 ± 0.06 MI, *p* < 1e–5). **E**. Experimentally recorded RS neuron contrast sensitivity slope changes during perceptual behaviors. Changes in contrast sensitivity slope were not significantly different with local PV stimulation (–1 ± 0.41 MI, 117 RS neurons) vs local SST stimulation (–1 ± 0.32 MI, 75 RS neurons; (*p* = 0.08). The slope decrease was significantly smaller with distal vs local PV stimulation (*p* < 1e–7) and with distal vs local SST stimulation (*p* = 0.021), but importantly, distal SST stimulation decreased the slope (–0.88 ± 0.54 MI, 146 RS neurons) significantly more than distal PV stimulation (–0.57 ± 0.55 MI, 166 RS neurons, *p* = 0.032), matching the model predictions.

**Figure S23.**
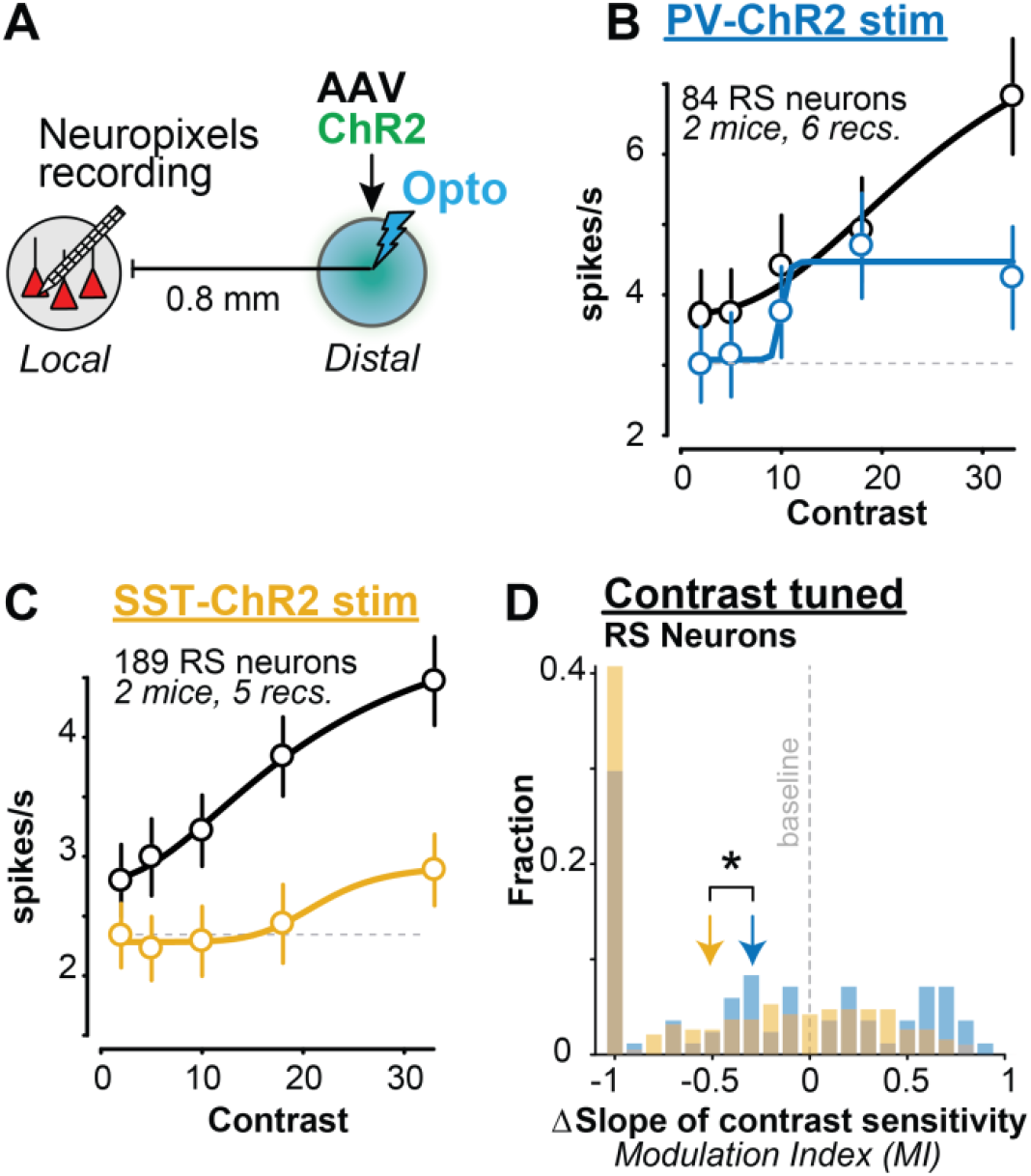
Changes in contrast sensitivity during distal stimulation with focal viral expression of ChR2. **A**. AAV-Flex-ChR2 targeted to monocular V1 (via intrinsic imaging of retinotopy) in PV- or SST-cre mice. Recordings retinotopically targeted to binocular V1. **B**. Average RS neuron contrast tuning (n=84) across during control (black) and distal PV stimulation. Only neurons with significant contrast tuning and decreases in spiking on laser vs control trials were analyzed. **C**. As in B, but for distal SST stimulation (189 RS neurons). **D**. SST stimulation evokes greater reduction in contrast sensitivity than PV stimulation (SST: -0.51 ± 0.52 MI, 189 neurons; PV: -0.29 ± 0.56 MI, 84 neurons; median ± MAD; p = 0.015). As in the Main results (Fig. 3), note the large fraction of RS neurons that lose all sensitivity to contrast with SST vs PV stimulation (MI = -1).

**Figure S24.**
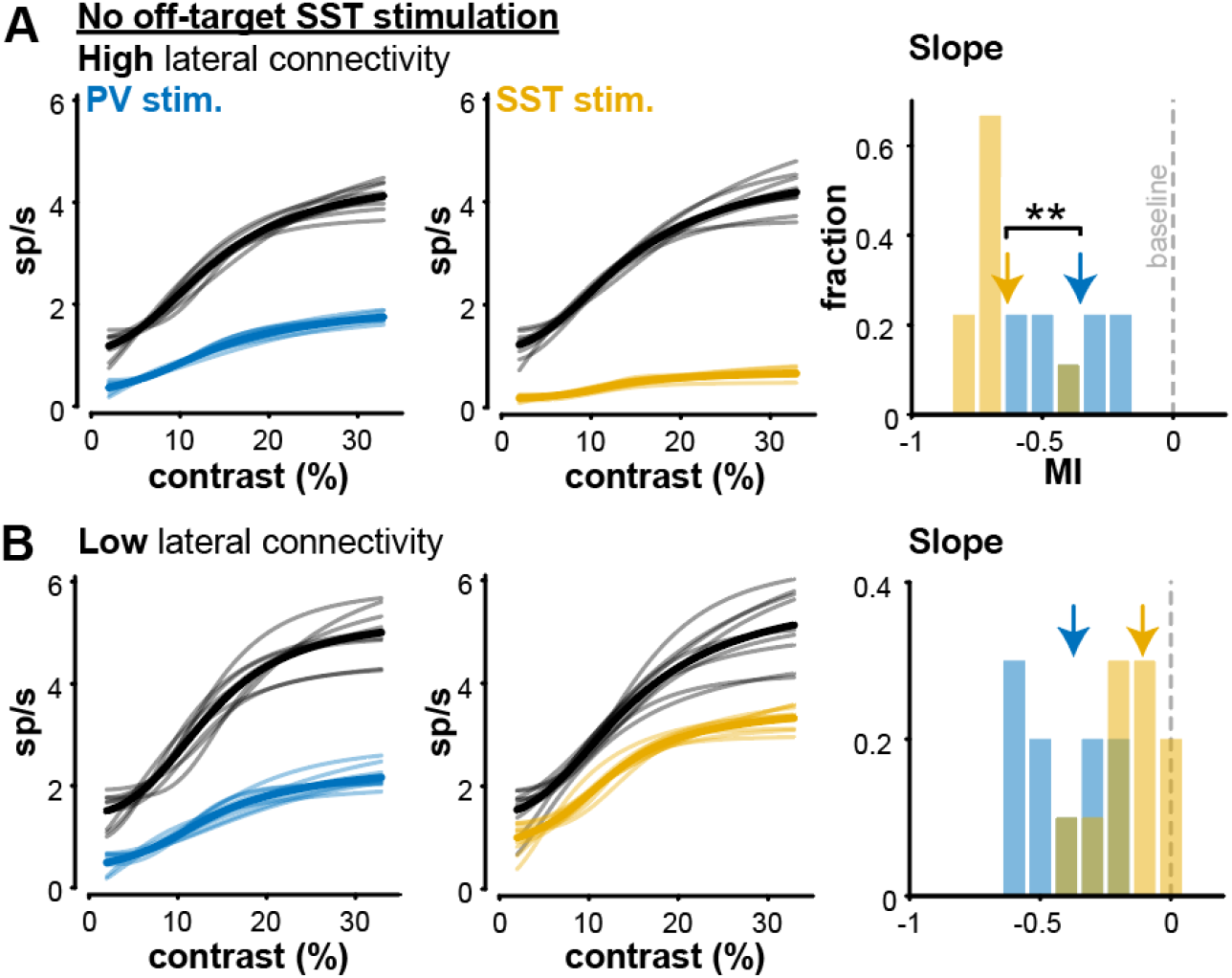
Antidromic photoactivation does not explain effects of SST stimulation. **A**. LIF circuit model (same as original) but without antidromic activation of a fraction (5%) of local SST neurons (Methods). Maintained significant decrease in slope of contrast sensitivity with SST but not PV stimulation (SST: –0.65 ± 0.03 MI, median ± mad; PV: –0.38 ± 0.14 MI; p < 1e-3, Wilcoxon rank sum test). Thin lines show model runs. **B**. Same model as A, but reducing SST lateral connectivity eliminates greater slope change with SST stimulation. (SST: –0.09 ± 0.07; PV: –0.31 ± 0.16; p = 0.998).

**Figure S25.**
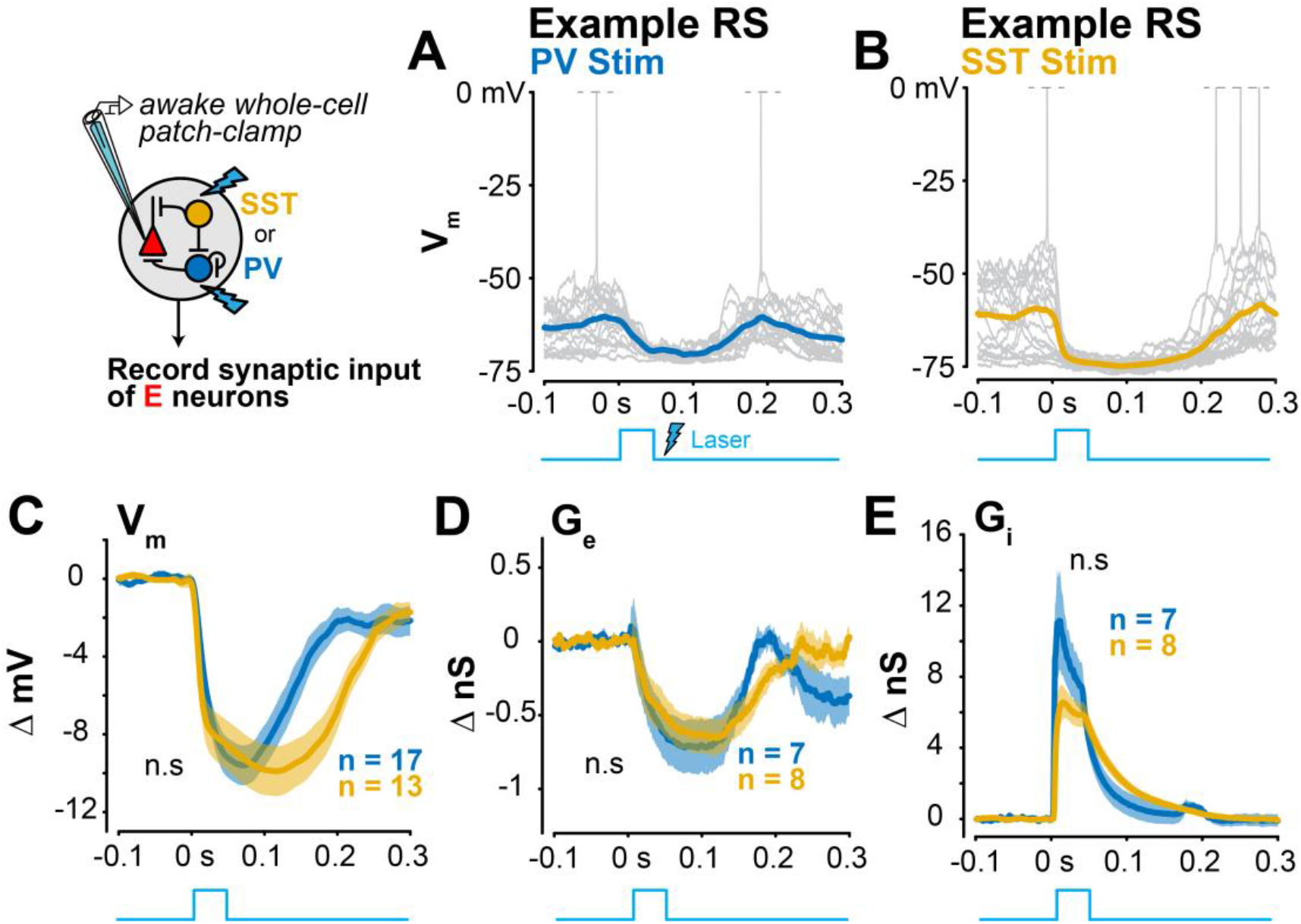
Changes in V_m_ and conductances in RS neurons during local PV or SST stimulation. **A**. Example whole-cell current clamp recording in awake V1 RS neuron during local PV stimulation (average across 20 trials plotted). Spikes truncated at 0 mV. **B**. Same as A for an example RS neuron during local SST stimulation. Spikes truncated at 0 mV. **C**. Hyperpolarization of RS neurons is not significantly different during local PV stimulation (ΔV_m_ = –6.35 ± 0.79 mV, mean ± SEM, 17 RS neurons) versus local SST stimulation (ΔV_m_ = –8.45 ± 1.16 mV, 13 neurons; *p* = 0.08, 1-tail Wilcoxon rank-sum test). **D**. Excitatory conductances are reduced with local PV stimulation (ΔG_e_ = –0.61 ± 0.18 nS, 7 RS neurons) and local SST stimulation (ΔG_e_ = –0.51 ± 0.13 nS, 8 RS neurons), but they are not significantly different from each other (*p* = 0.39). **E**. *Inhibitory conductances* increased with local PV stimulation (ΔG_i_ = 4.85 ± 0.92 nS, 7 RS neurons) and local SST stimulation (ΔG_i_ = 4.27 ± 0.56 nS, 8 RS neurons), but were not significantly different from one another (*p* = 0.43).

**Figure S26.**
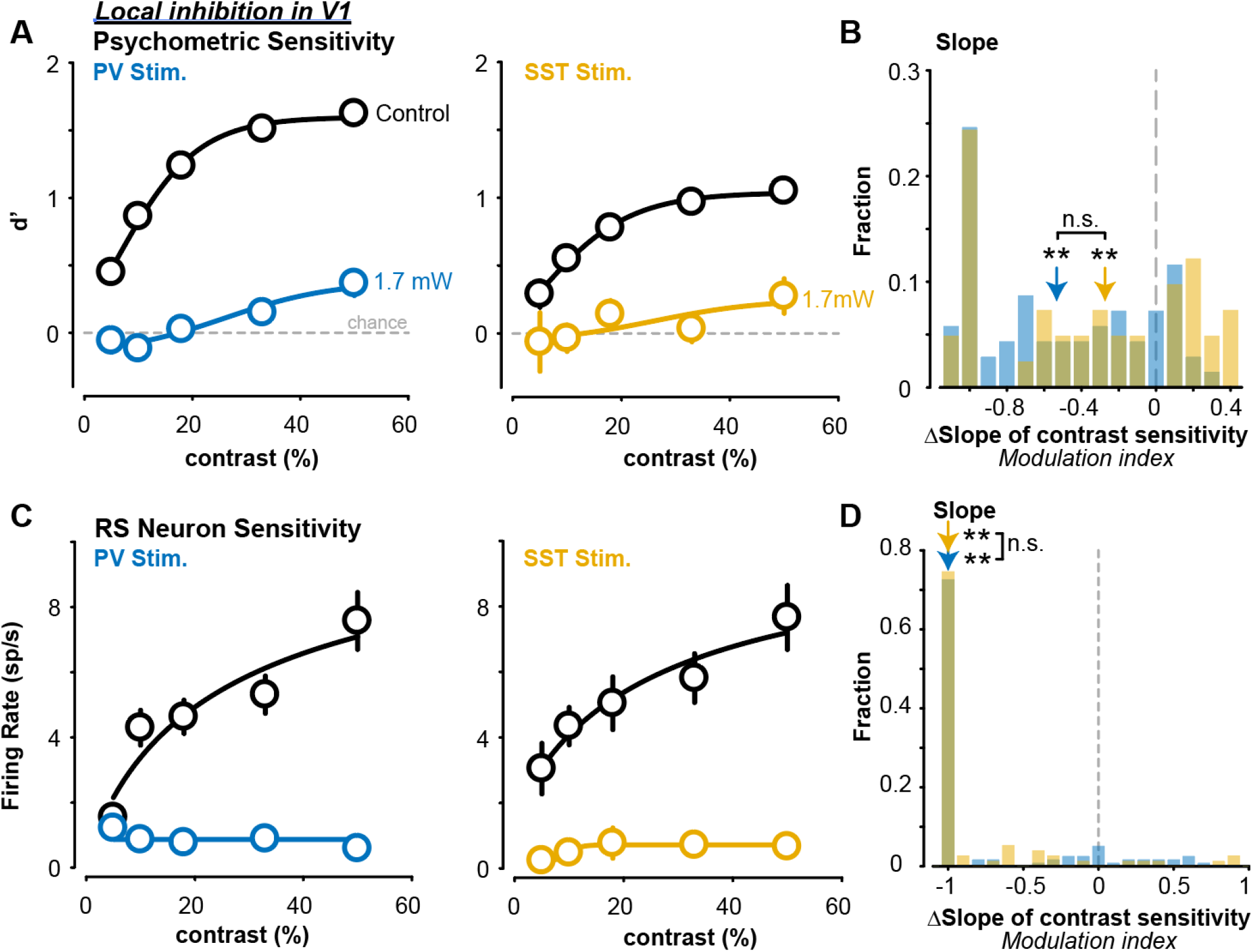
Contrast sensitivity changes during local activation of PV and SST neurons. **A**. Average psychometric sensitivity across contrasts during local PV stimulation (left) and SST stimulation (right). Only sessions with similar contrasts plotted for visualization purposes. **B**. Local PV (6 mice, 84 sessions) and SST stimulation (6 mice, 57 sessions) elicits similar reductions in the slope of psychometric contrast sensitivity (slope change MI: PV = -0.53 ± 0.41, median ± mad; SST = -0.27 ± 0.47; p = 0.062, 1 tail rank sum test). Both PV (p < 1e-8, sign rank test) and SST (p < 0.01) stimulation significantly decrease the slope. **C**. RS firing rate across contrasts during local PV stimulation (left) and SST stimulation (right). **D**. Change in RS neuron contrast sensitivity was not significantly different with local PV or SST stimulation (PV: –1 ± 0.41 MI, 117 RS neurons; SST: –1 ± 0.32 MI, 75 RS neurons; p = 0.08). Both PV (p < 1e-17) and SST (p < 1e-12) stimulation significantly decrease the RS neuron contrast sensitivity slope.

**Table S1.**
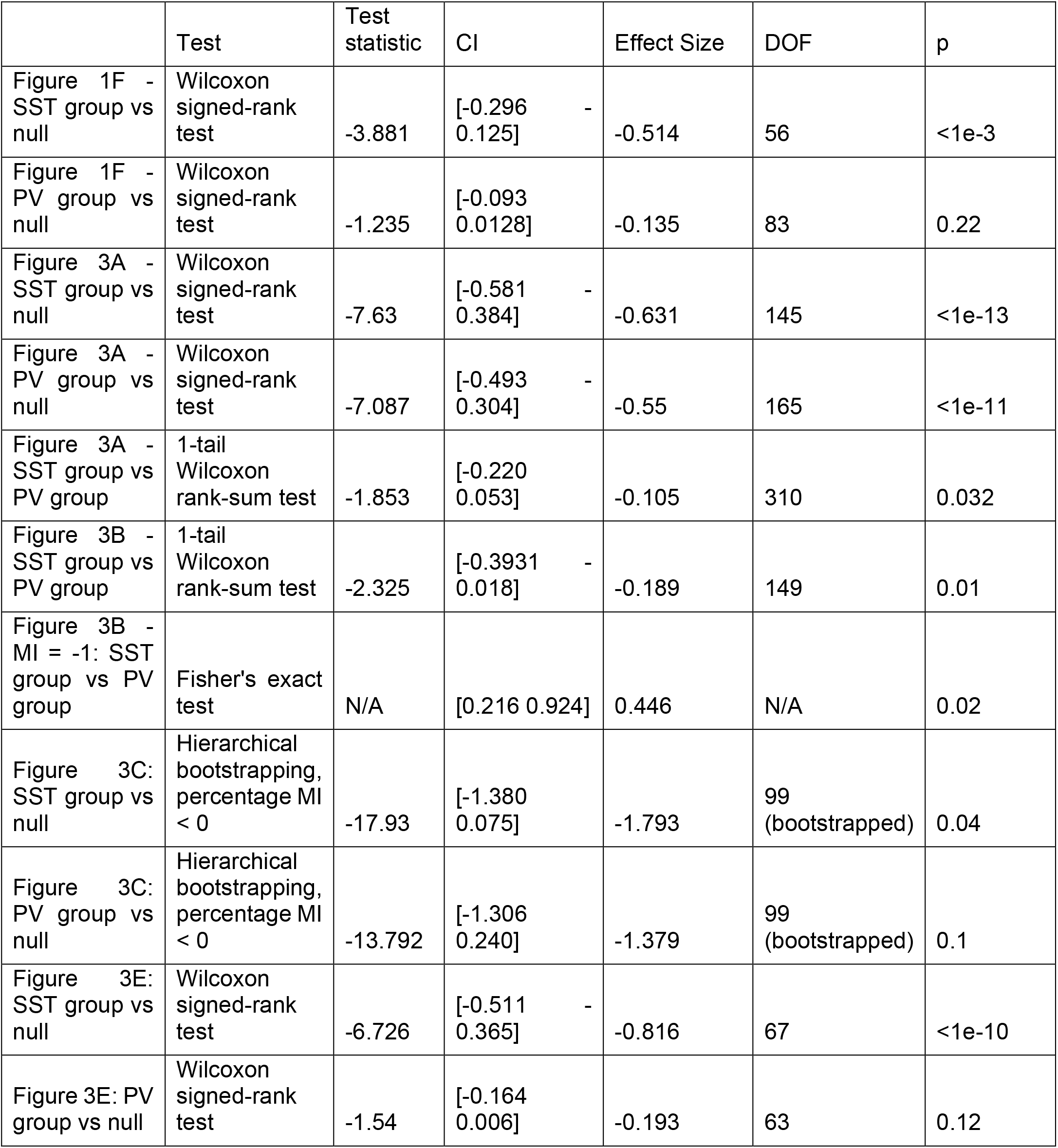
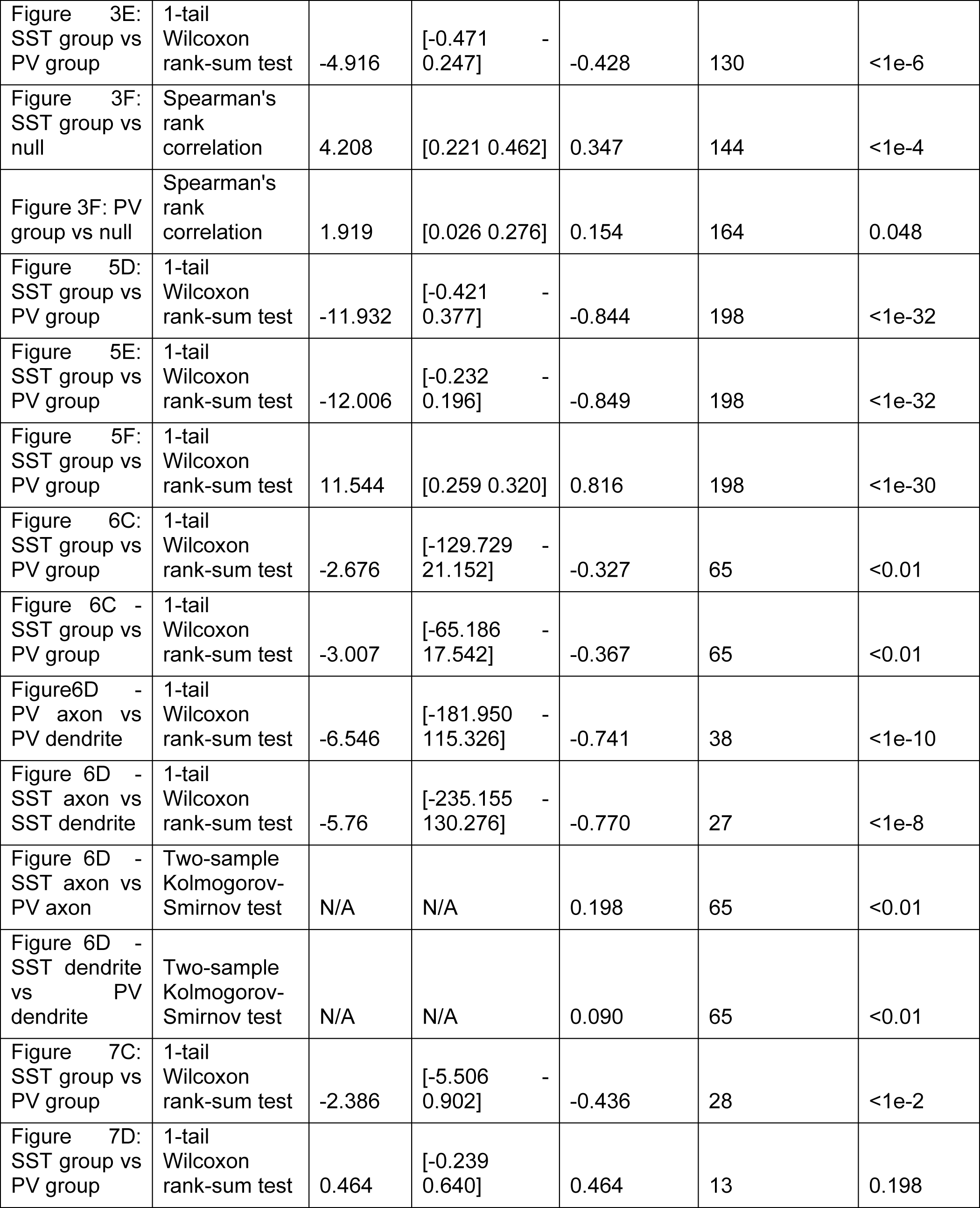
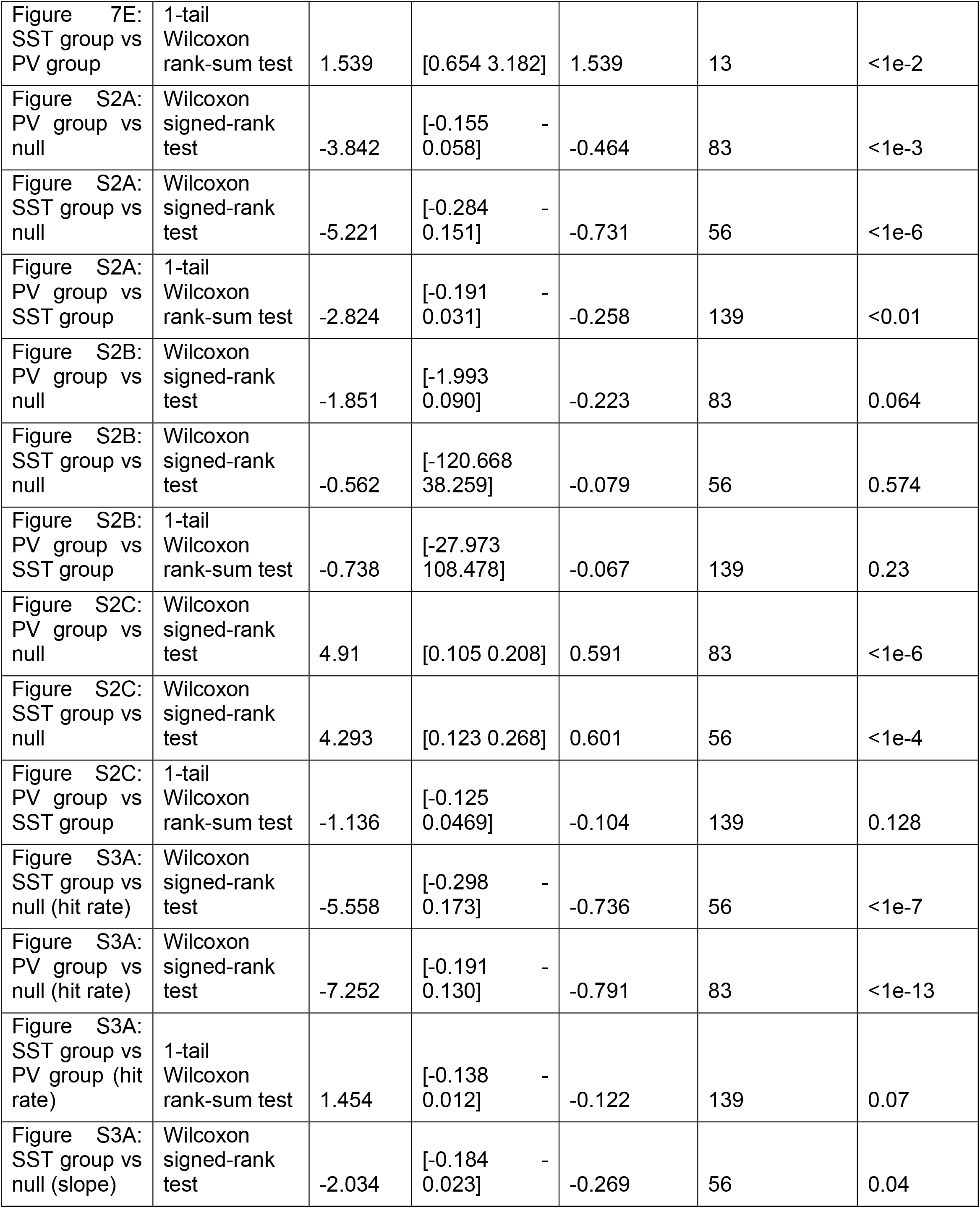
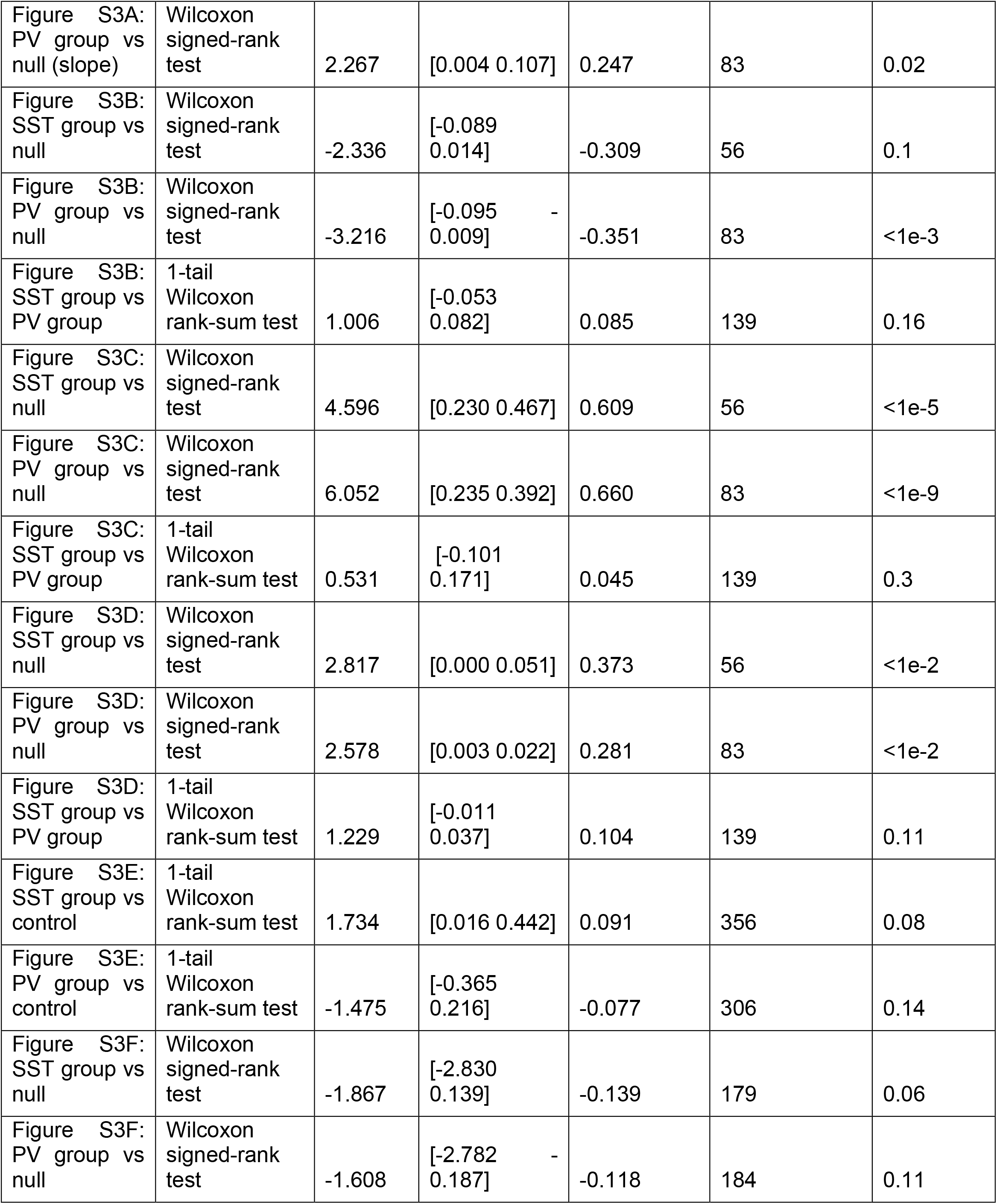
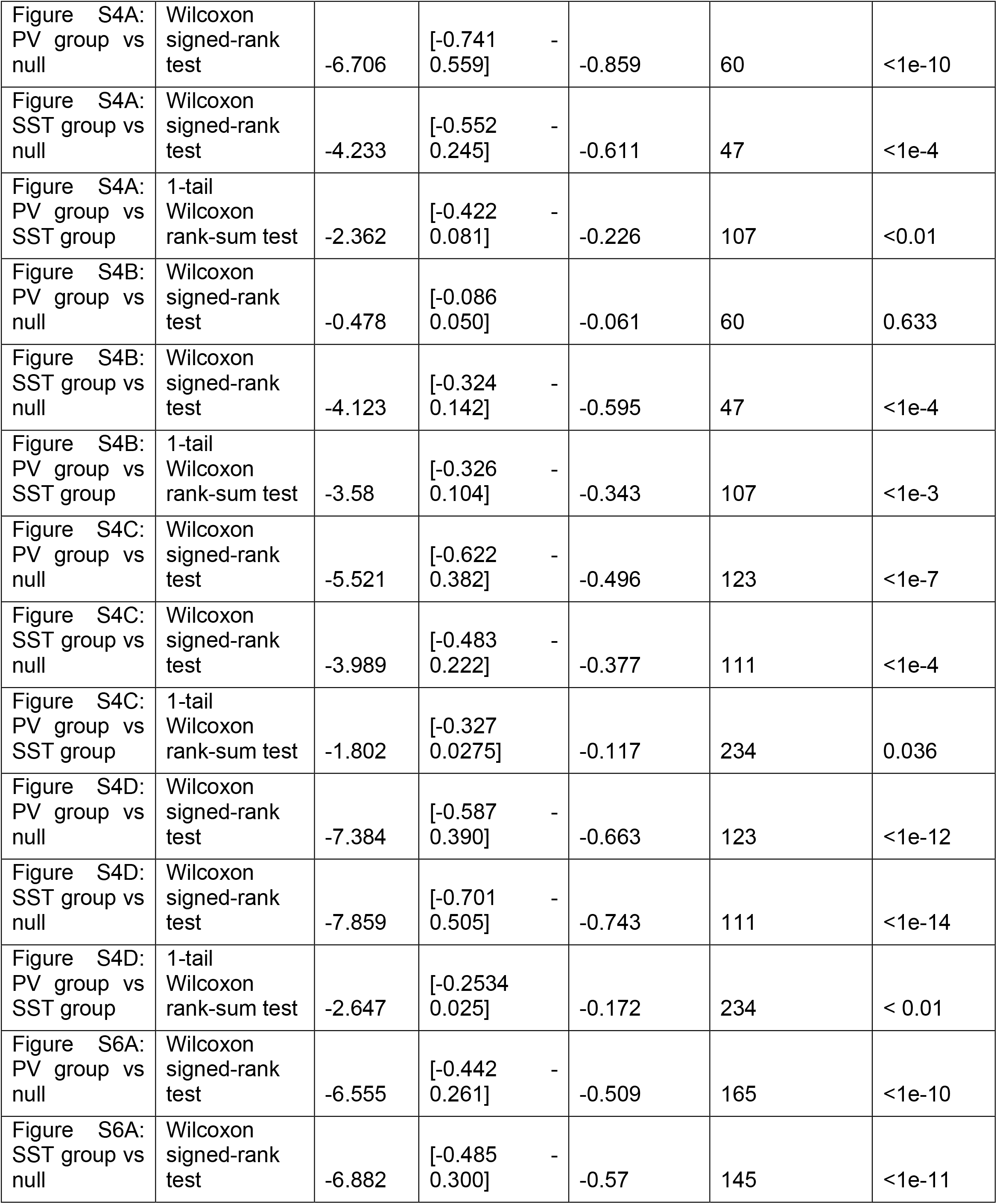
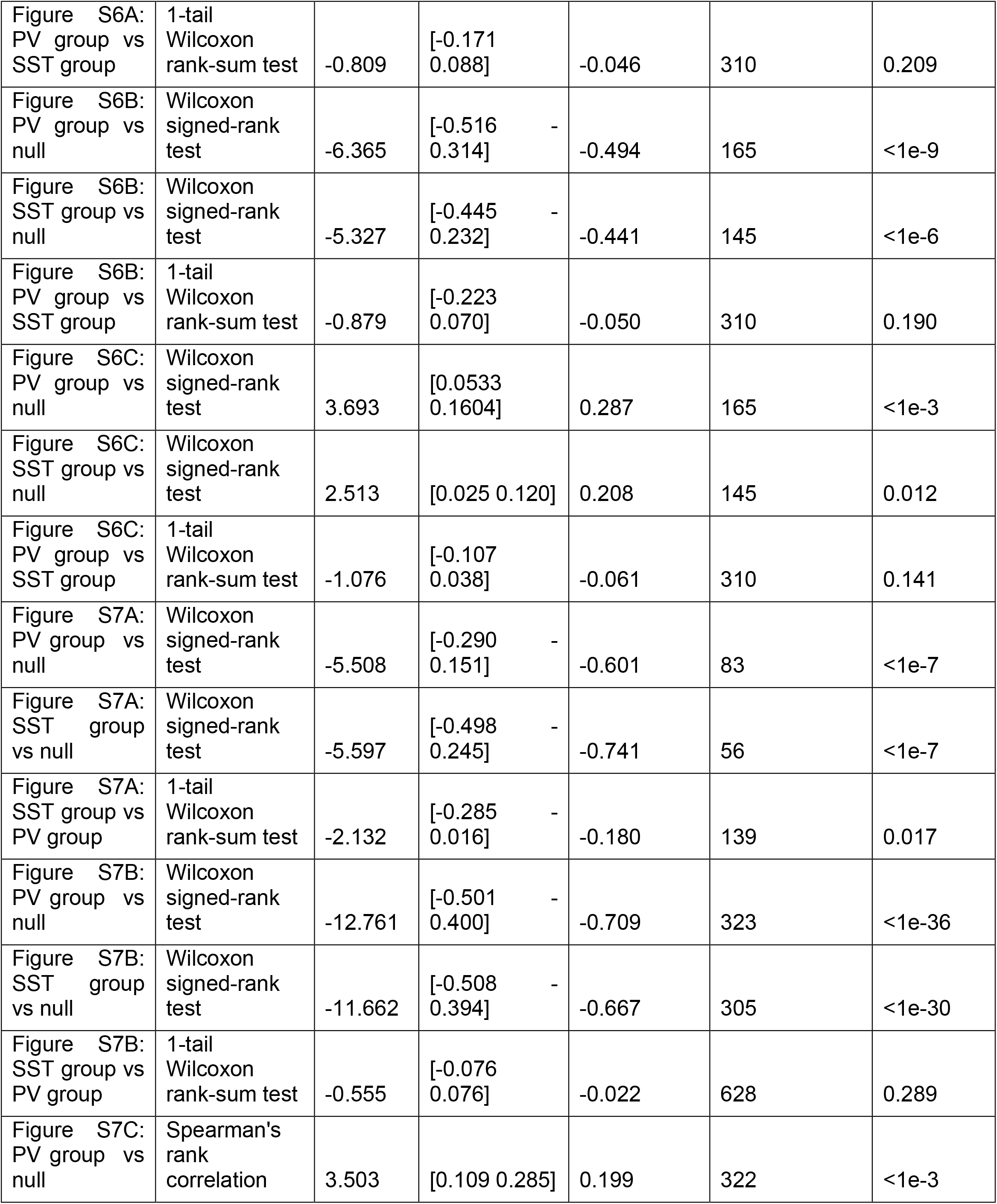
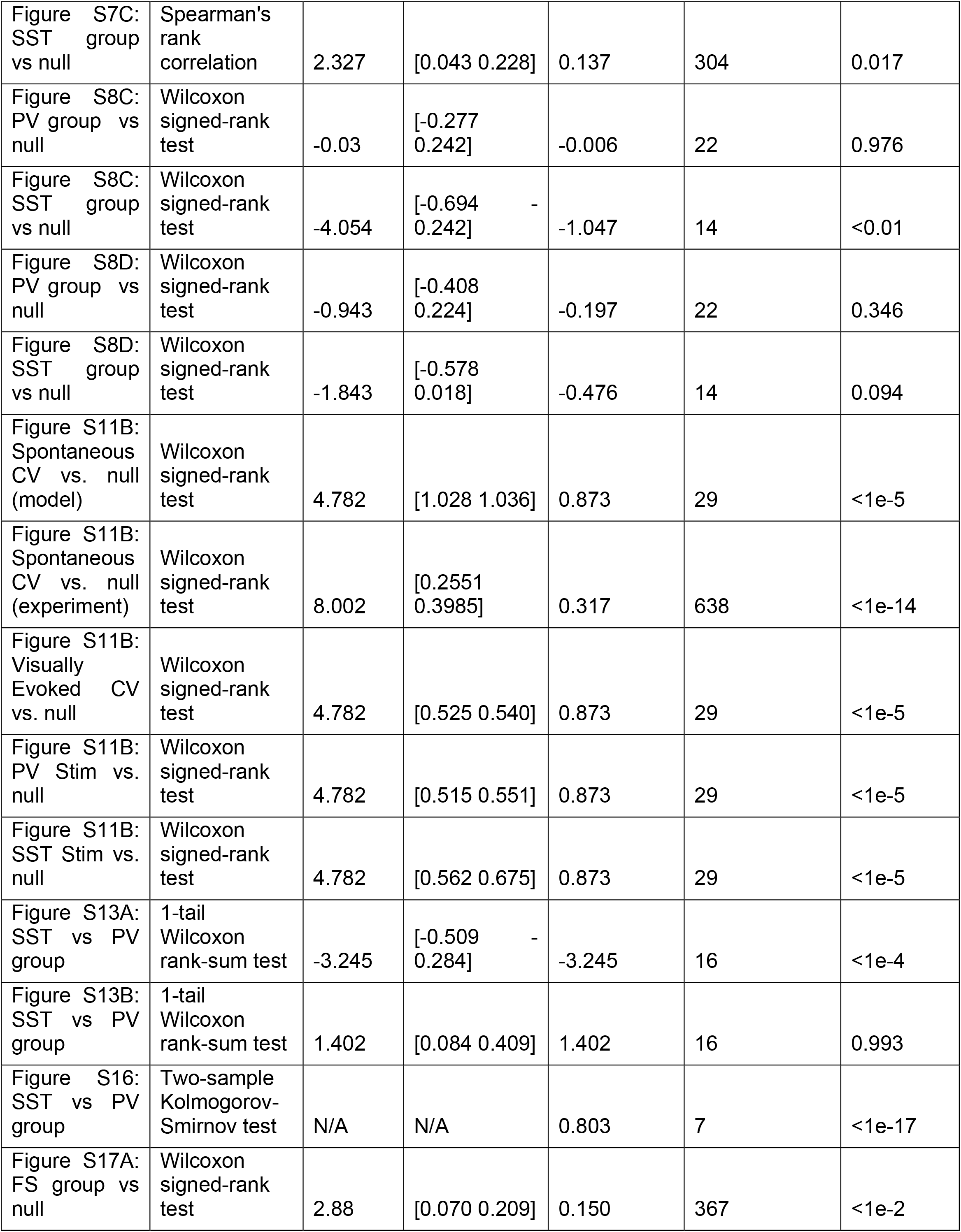
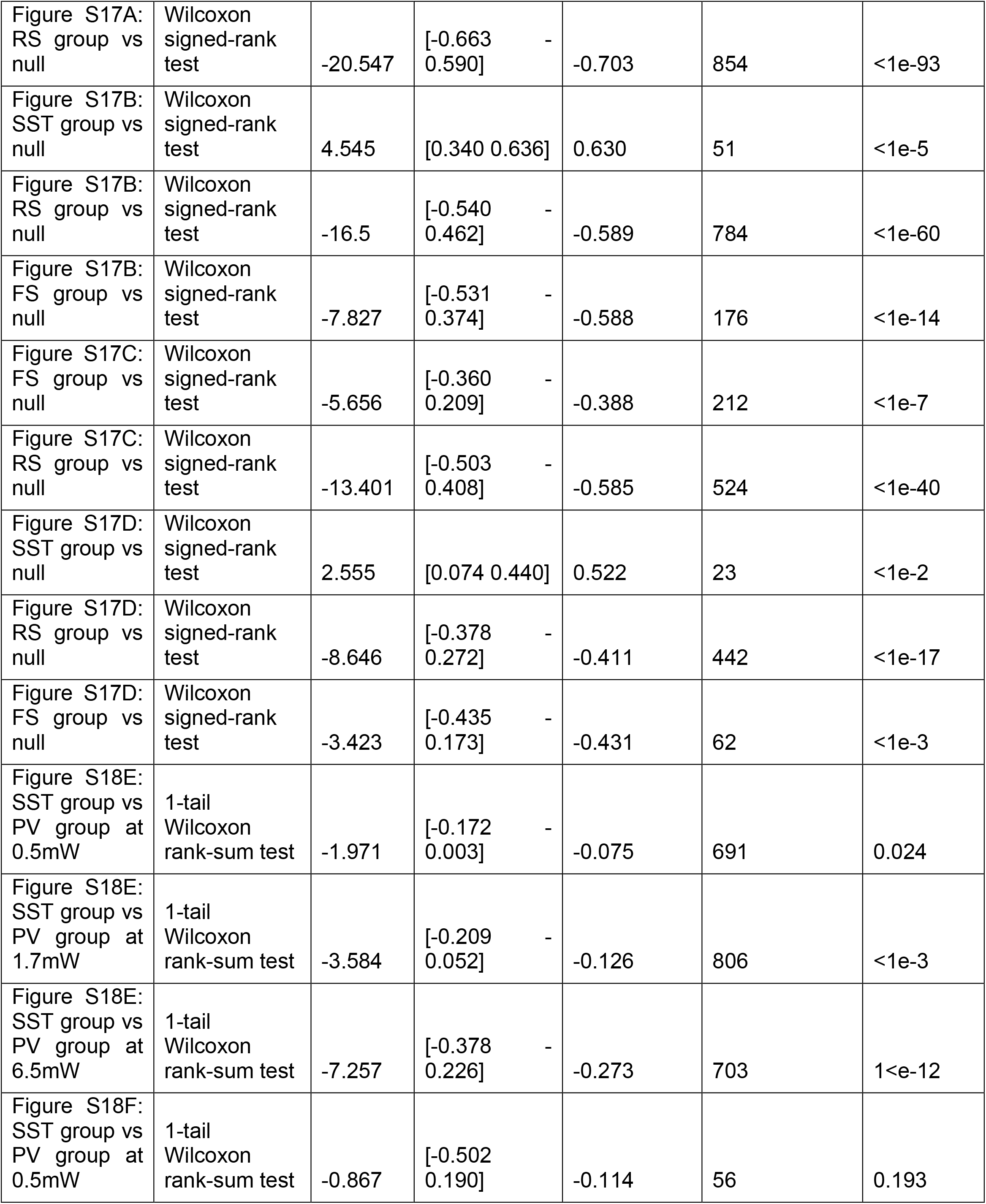
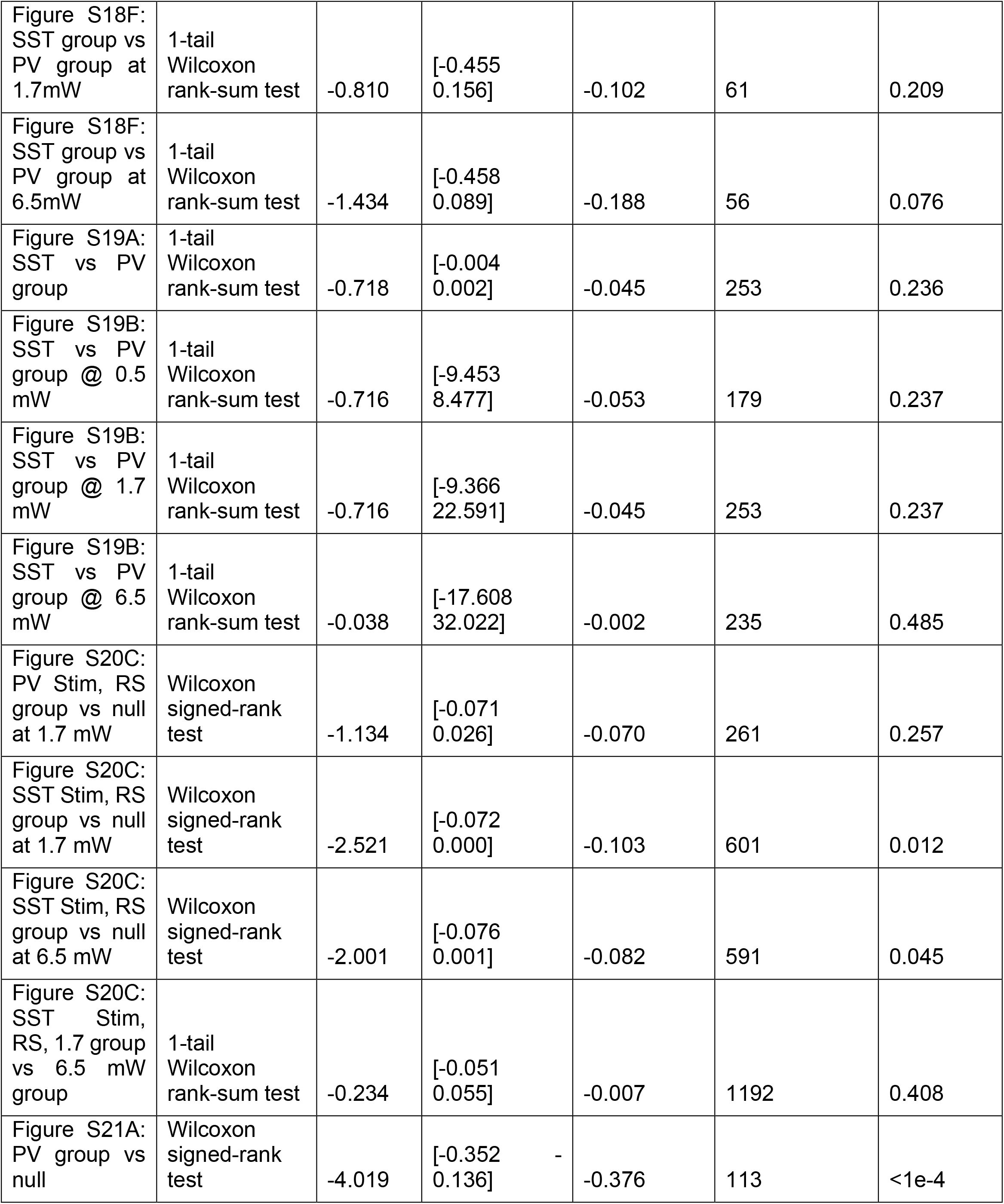
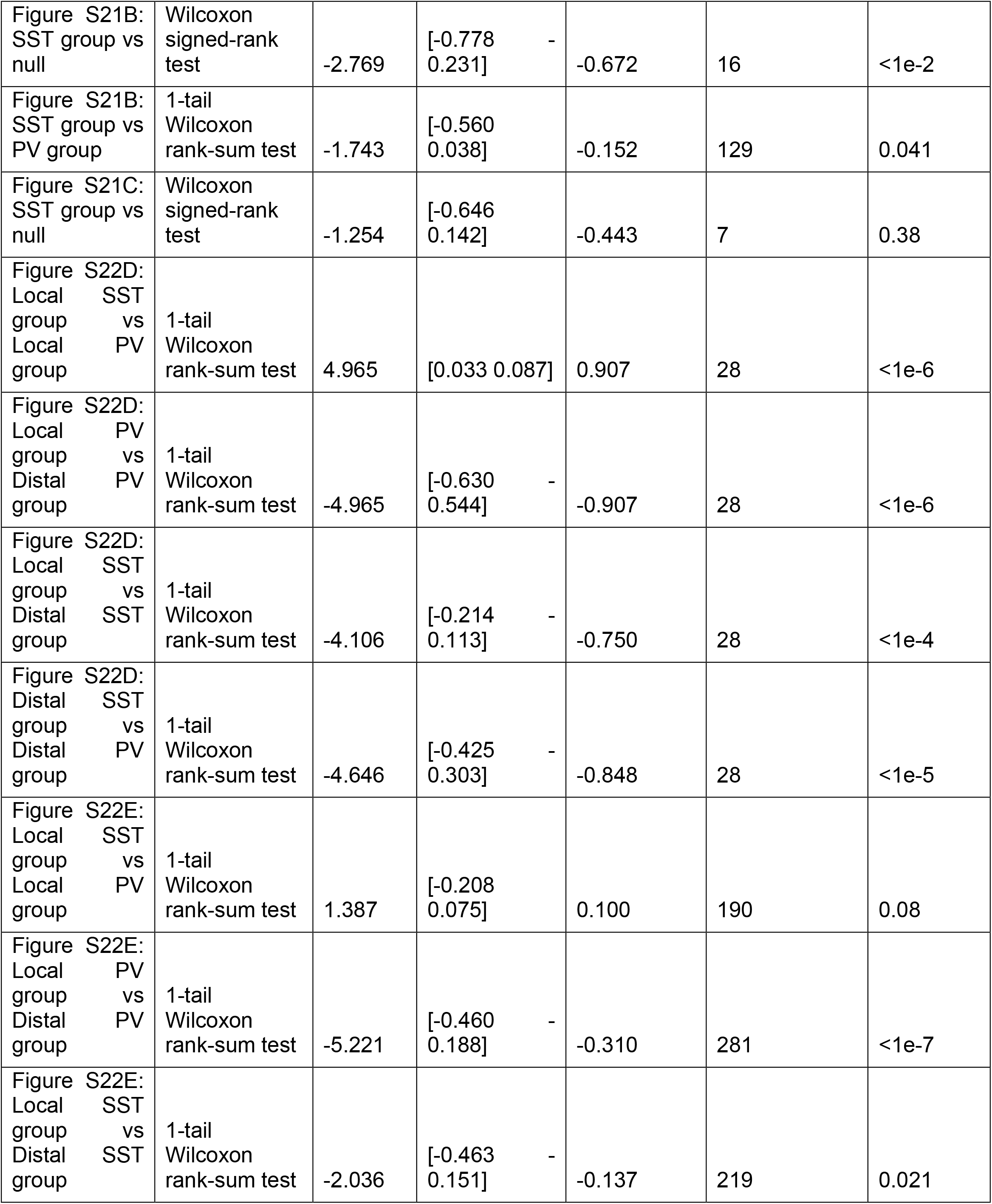
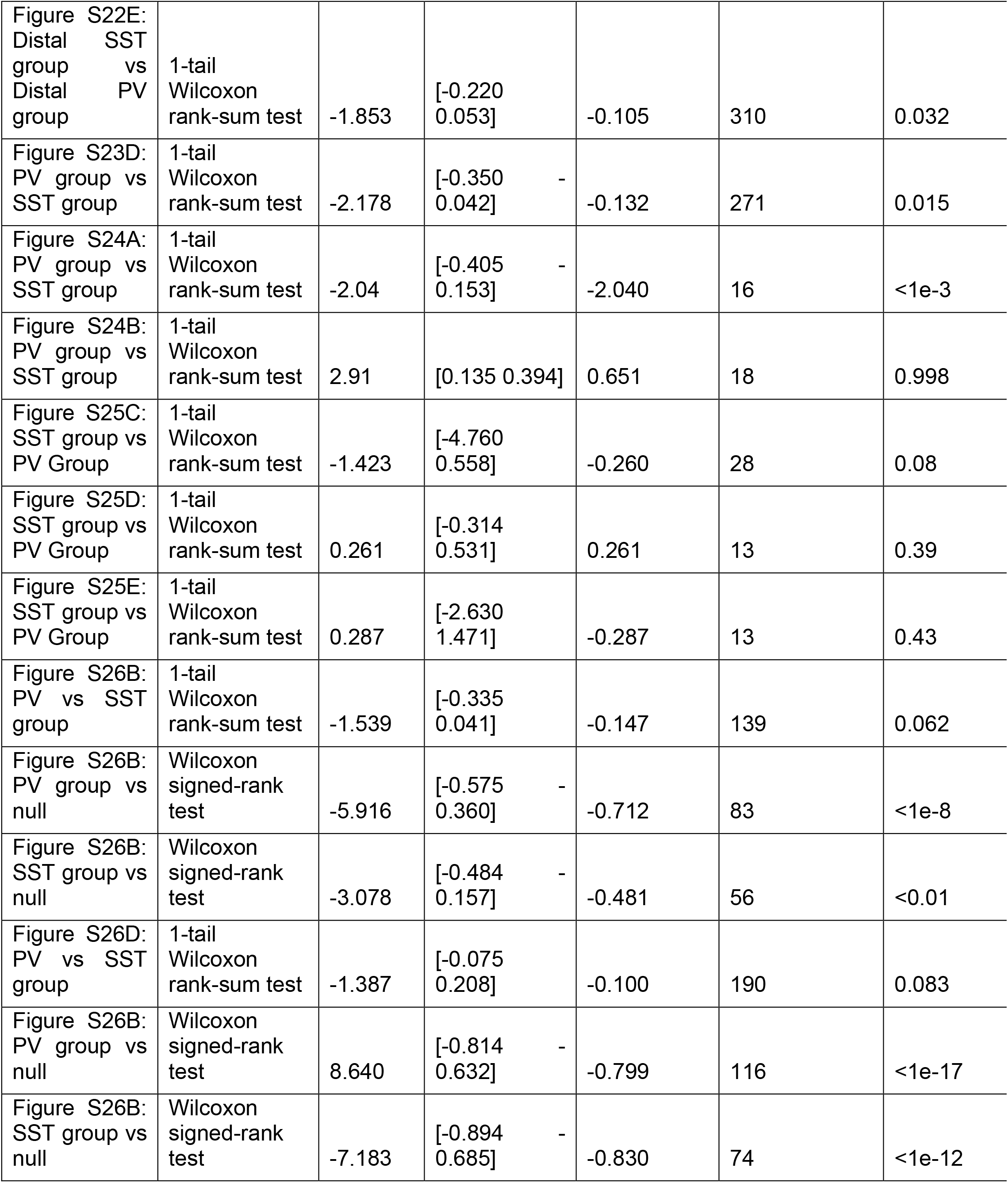
Statistical analysis. Detailed statistical results used in analysis. Test statistic calculated using z-statistic (or t-statistic when sample size was low, n < 30). Unless otherwise specified, effect size was calculated using r (or Cohen’s d for low sample size).

